# Interpretable Machine Learning Identifies an Emergent Absence Seizure Mechanism

**DOI:** 10.1101/2025.09.23.678032

**Authors:** Jacob M. Hull, Nicholas Denomme, Surya Ganguli, John R. Huguenard

## Abstract

Absence seizures are widespread spike-and-wave oscillations disrupting consciousness. Several consciousness frameworks emphasize intercortical/thalamocortical feedback dynamics, but we lack explicit dynamical mechanisms for how seizures disrupt these circuits. Using interpretable machine learning, we derived dynamical equations directly from seizure electrocorticogram data, reproducing seconds-long multi-regional local field potentials with precision matching inter-mouse variability. The model contained a low-dimensional chaotic seizure attractor emerging from between-region synchronization at preferred phase-lags. Unit recordings revealed corresponding spiking synchrony at single-neuron resolution, linking somatosensory and motor cortex with posterior thalamic nucleus (PO). Model coupling terms predicted tonic and burst firing spatiotemporal organization across regions and guided multisite-optogenetic stimulations. These stimulations showed PO controls corticocortical connectivity gain and that motor/somatosensory corticocortical functional connectivity varies at predicted phase-lags to drive seizures. Our results define absence seizures as dynamics confined to a chaotic attractor within a circuit implicated in anesthetic unconsciousness, linking distributed network chaos to loss of consciousness.

**Graphical Abstract:** 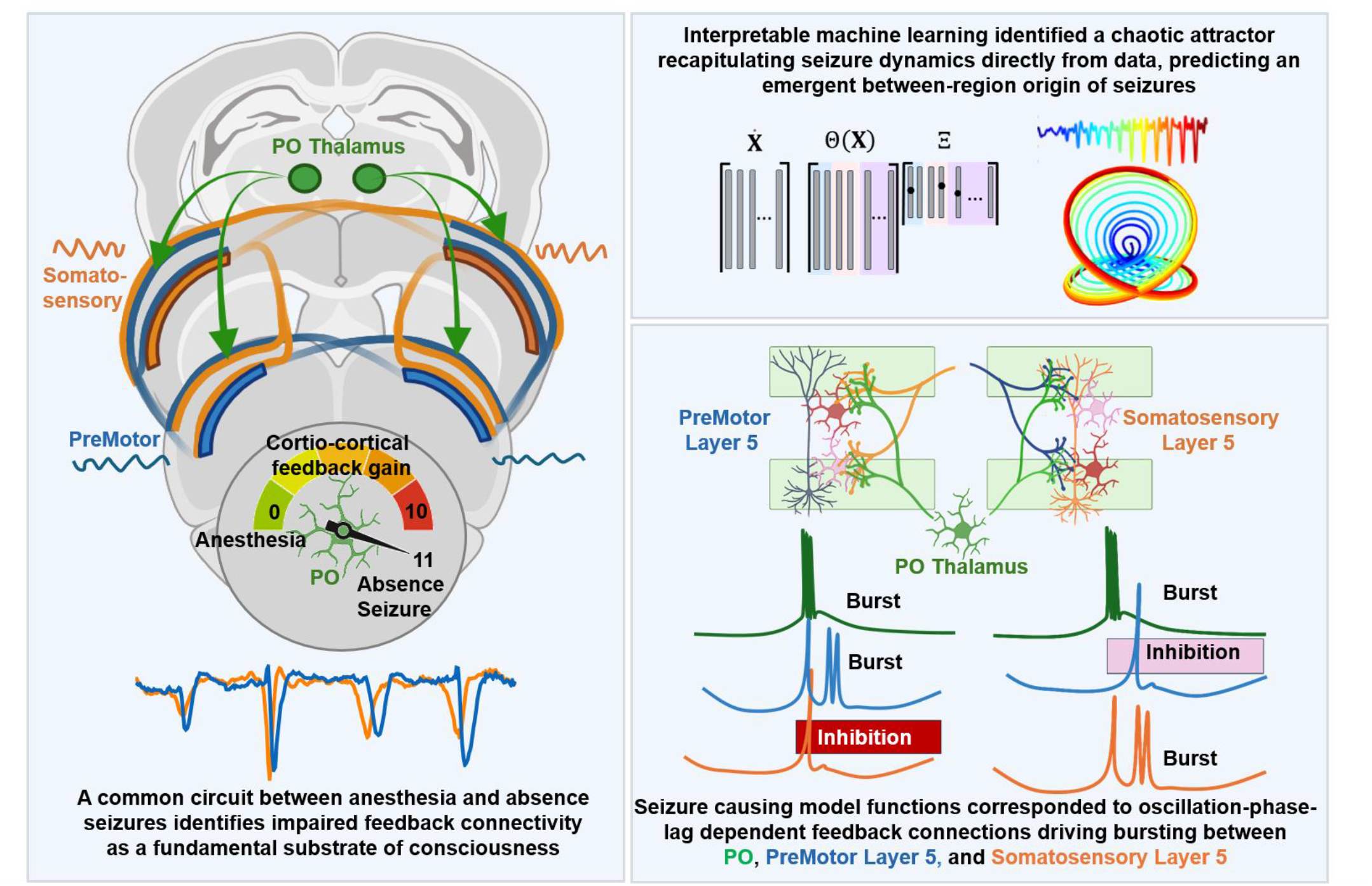

## Introduction

Absence seizures are generalized non-convulsive seizures characterized by impaired consciousness and widespread bilaterally symmetric 3 Hz spike-and-wave oscillations (SWD) on EEG. SWDs are multicomponent sequences of negative spikes accompanying lower frequency positive waves. Their onset is generalized across bilateral frontal/parietal regions, implying multi-regional mechanisms for their generation or rapid spread between regions. The cortical focus theory in rat absence models posits a single somatosensory cortical focus that rapidly spreads.^1^ A core feature of this theory is that somatosensory face/whisker cortical hyperexcitability drives initiation of each seizure, which then rapidly recruits other thalamocortical networks leading to near global cortical synchrony.^2^ Recent in vivo multielectrode and fMRI evidence however suggests little or no somatosensory cortical hyperexcitability in the transition to seizure leading to increased interest in how oscillatory firing and synchrony are coordinated.^3–6^ Absence seizure mechanisms involving hyperexcitability and oscillatory activity may however be interdependent. For example, focal cortical activation can recruit thalamic reticular neurons, inhibit thalamocortical cells and trigger rebound firing that completes an oscillatory cortico-thalamic feedback loop.^7–11^ Consistent with this mechanism, in vitro stimulation of cortical fibers projecting to the thalamus in a pattern mimicking high-frequency cortical bursting, transforms spindle-like (6–10 Hz) thalamic oscillations into hypersynchronous, SWD-like (3–4 Hz) thalamic bursts.^7, 8^ Acting in the opposite direction, thalamic stimulations in cat that induce spindle oscillations can be converted to SWDs when cortical excitability is increased with GABA_A_ blocker penicillin. This conversion from spindle to SWD oscillations occurs through progressive amplification of every second spindle peak, suggesting SWDs emerge from an underlying spindle-like cortico-thalamic oscillation at twice the SWD frequency.^12^

Long-range reciprocal feedback connections, can in principle lead to population level oscillations and synchrony, thereby coordinating sensation and action, and may support consciousness through cortical and subcortical mechanisms.^13–15^ Cortico-thalamic oscillations may be nonlinear, multi-regional, and high dimensional, suggesting the need for a quantitative framework. One framework for understanding large neuronal population dynamics is the concept of attractor states in which system dynamics are confined to a subset of neuronal activity patterns.^16^ Attractors have been identified in many circuits including those involving navigation, motor planning, perceptual decision making, and epilepsy.^17–23^ Such mechanisms may account for the recurrent and stereotyped activity patterns underlying absence epilepsy. While a dynamical systems perspective has greatly expanded our conception of seizure mechanisms beyond excitatory/inhibitory imbalance^23^, a central challenge is discovering how such abstract dynamics arise from specific cellular substrates. Here we combined recent advances in system identification via machine learning, high-density recording, and multisite optogenetic stimulation to systematically investigate the dynamics of absence seizure generation. We used interpretable machine learning to identify a highly accurate system of dynamical equations directly from seizure local field potential (LFP) data. This system accurately simulates seizures to within the high precision of inter-mouse variability. These equations gave rise to a chaotic attractor, where between-region oscillation coupling and phase-lags, rather than a region-specific deficit, drove absence seizures. High density multielectrode recordings and multisite optogenetics then allowed us to identify that this between-region phase-lag dependent seizure drive corresponded to the relative timing of burst firing in layer 5 pyramidal neurons of the motor and somatosensory systems and the amplification of cortico-cortical connectivity gain via higher order thalamic nucleus PO. A detailed summary of our results is given in the discussion section, which may be helpful to read before the results.

## Results

### Absence seizures are stereotyped by sensory-motor system relationships but not trajectories in time

We first aimed to uncover the dynamical laws governing absence seizures by examining seizures in a well-characterized mouse model (med mice, hemizygous for *Scn8a*, encoding Nav1.6).^24^ We developed a high-fidelity electrocorticogram (Ecog) approach with electrodes over 16 cortical regions (Fig. 1A). Individual spikes of a SWD complex in mice span ~133 ms (7.5 Hz), with additional rhythms shaping patterns within and between SWDs (Fig. 1B). Seizure power and onset times were quantified using our “seizure score” metric (Fig. S1 A-D). Median seizure onset times showed no detectable latency differences among premotor left/right (PreMotL/R), primary motor (PrimMotL/R) and somatosensory face (FaceL/R) cortices-all with onset within a single SWD interval (Fig. 1C).^25^ The somatosensory barrel cortex (BarL, BarR) onset was modestly delayed and greater delays were observed in other regions.

**Figure 1.**
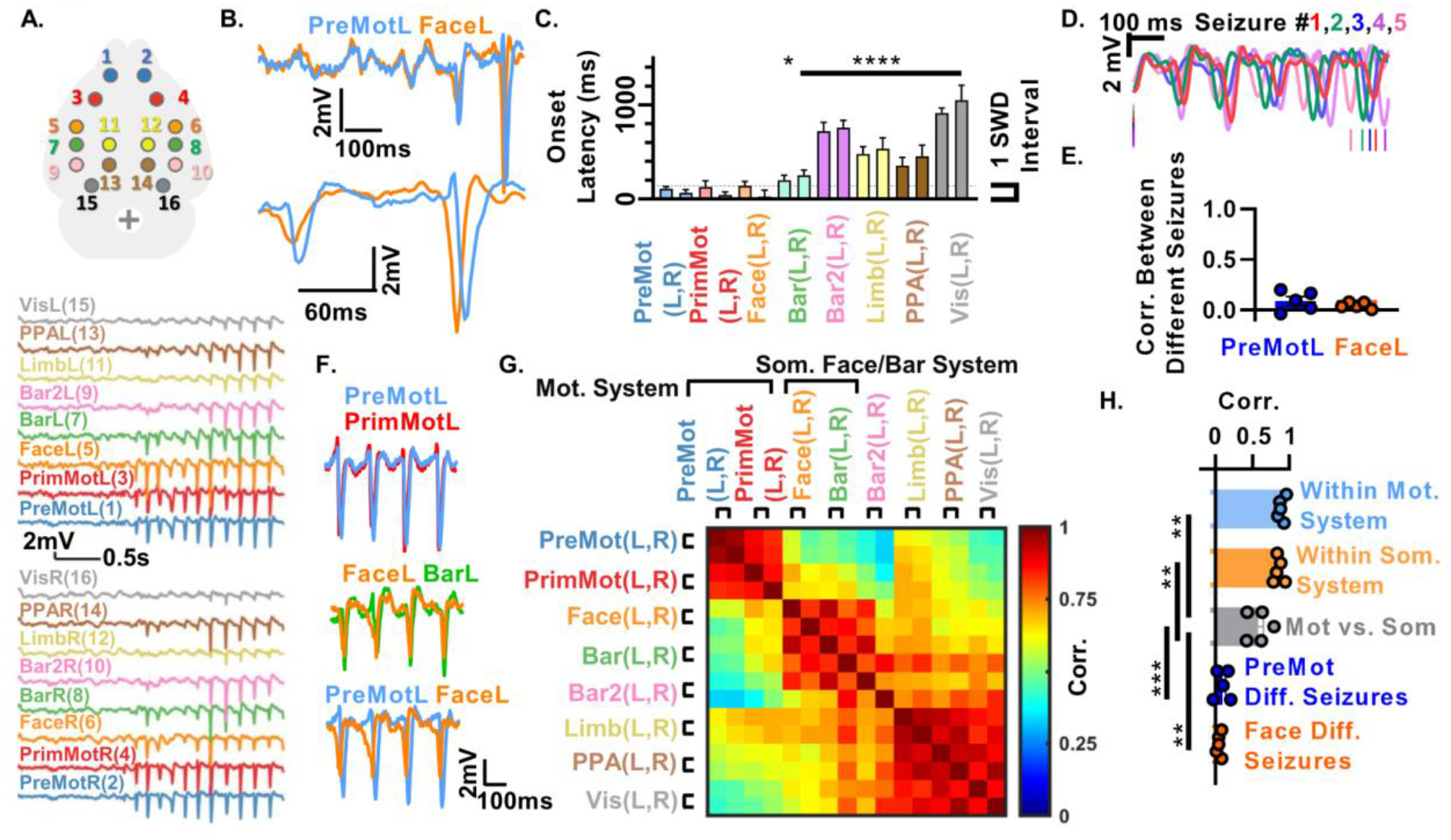
Between-region relationships describe absence seizure dynamics better than temporal trajectories. (A) Recording location schematic (top) and example Ecog traces (bottom). (B) Example spike-and-wave discharge (SWD) onset in premotor left (PreMotL) and somatosensory FaceL (FaceL) regions, expanded view in bottom. (C) Median onset latency ± 95% CI relative to the earliest onset site of each seizure (*N* = 5 mice, 30 seizures per mouse). (D) Example traces of five seizures from PreMotL in one mouse aligned to the peak of the first SWD-spike crossing (< −1000 mV); vertical lines mark SWD-spikes at first and sixth crossings. (E) Correlation between seizures within the same animal aligned as in (G): PreMotL 0.09 ± 0.04, FaceL 0.05 ± 0.01 AU. (F) Example motor (PreMotL, PrimMotL) and somatosensory (FaceL, BarL) system waveforms are similar within motor (top) and somatosensory (middle) systems but not between systems (bottom). (G) Average correlation between all region combinations (*N* = 5 mice, 5 seizures per mouse). (H) Within-system correlations: Mot 0.89 ± 0.02, Som 0.85 ± 0.03 vs between-system 0.58 ± 0.07 AU vs between different seizures from recorded from the same region of the same animal from E. Statistical analyses were performed by Cox proportional hazards model (C) and standard or Welch’s two-tailed Student’s *t* test (H); *N* = 5 mice. Data represent mean ± SEM except in C where data are median and 95% CI.

We next examined the temporal trajectories of seizures to identify regularities that can be used to understand seizure dynamics. Comparison of trajectories from the same region of the same mouse revealed that different individual seizures quickly diverge from each other in time, with SWD-spikes spanning the full SWD cycle within 1s (Fig. 1D) and with negligible signal correlation between different seizures (Fig. 1E). In contrast, relationships between regions were robustly maintained during seizures. Among early-onset areas, PreMotL and PrimMotL of the motor system showed highly similar SWD waveforms (Fig. 1F, top), as did FaceL and BarL of the somatosensory system (Fig. 1F, middle), while motor vs. somatosensory system signals were more distinct (Fig. 1F, bottom). We quantified similarity using pairwise correlations over the first 3 s of seizure across all region combinations (Fig. 1G). Motor and face/barrel somatosensory systems showed strong, nearly identical correlations within the respective systems and weaker but still strong between-system correlations (Fig. 1H). Comparing these between-system correlations to the between-different-seizure correlations of the same region of the same mouse (Fig. 1E) revealed that between-region relationships far better describe seizure dynamics than temporal trajectories (Fig. 1H).

Overall, these results revealed an early motor and somatosensory seizure network, with highly stereotyped signals within and between these systems, but with rapidly diverging dynamics in time between different seizures.

### Projection onto a toroidal coordinate system revealed highly constrained phase and amplitude dynamics during absence seizures

Fig. 1D and E suggest significant oscillatory phase drift as each seizure unfolds, while Fig. 1F-H suggests waveforms are stereotyped within-systems and that pairs of regions maintain robust phase locking during seizures, especially within and between somatosensory and motor systems. These results motivated a switch from time-voltage coordinates to coordinates that directly represent within- and between-region oscillatory phase-lags and dynamics. Hereafter, we refer to PreMot Ecog LFP signals as Mot and somatosensory face as Som unless otherwise specified. To gain a clear geometric view of within- and between-regional SWD oscillation dynamics, we utilized the dominant frequency bands in the SWD spectrogram. The SomL spectrogram (Fig. 2A) revealed a fundamental frequency in a ~7.5 Hz band (band 1) that slightly decays over time, accompanied by two substantial harmonics at 15 Hz and 22.5 Hz (bands 2 and 3). This 3-band pattern was seen across electrodes (PreMotL, FaceL, Visual L; Fig. S1A). We constructed three filters (Fig. 2A, right) to extract phases and amplitudes of these oscillations via the Hilbert transform (schematic, Fig. 2B).

**Figure 2.**
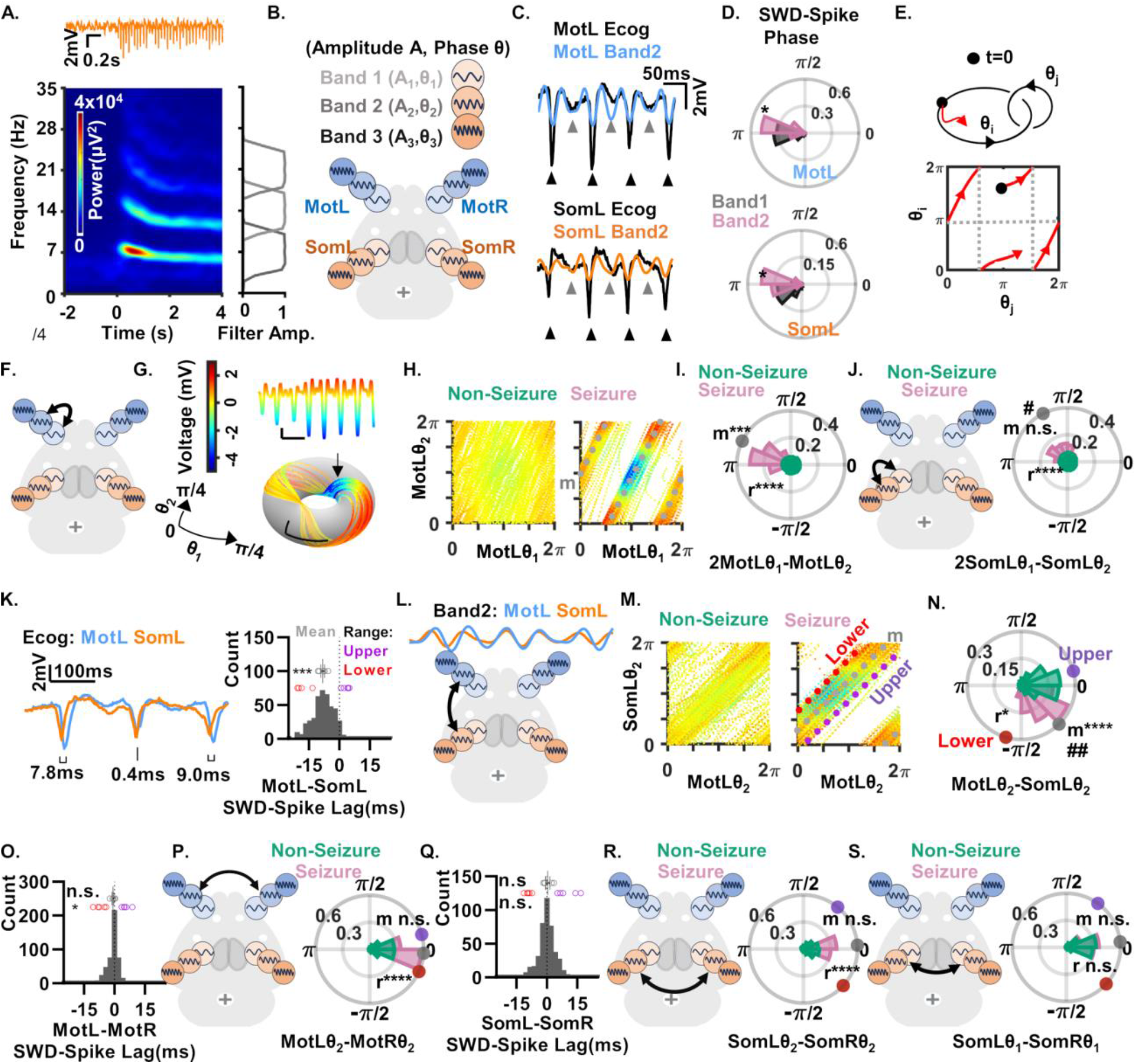
A toroidal state space defines absence seizures. (A and B) Average spectrogram of FaceL seizures (A, bottom-left) aligned to an example FaceL seizure (A, top) and corresponding band filters (A, right). Schematic (B) of Mot (L,R) and Som (L,R) signals composed of three harmonics with amplitudes A_i_ and phases θ_i_. (C) Example traces of full spectrum SWDs (black) and band 2 (blue, orange) in MotL (top) and SomL (bottom). Black arrows indicate negative SWD-spike troughs aligned to band 2 troughs (π rad). Gray arrows indicate Ecog troughs between SWD-spikes aligned with band 2 troughs. (D) Circular histograms of phase of SWD-spikes in bands 1 and 2. Asterisks indicate higher circular standard deviation (more variability) for band 1 than band 2, paired t-test on per mouse circular standard deviation. See Fig. S2A. (E) Toroidal coordinate system (top) and unwrapped representation with periodic boundary wrapping (bottom) illustrating an example phase trajectory (red) and phase wrapping at edges (dashed lines). (F and G) Schematic (F) of the MotL band 1 and 2 interaction. Experimental trajectories (G, bottom) on the toroidal manifold of bands 1 and 2 for MotL color denotes summed amplitudes of the three bands (G, top). Arrow indicates tight cluster of all SWD-spikes. Contrast to dispersed spikes in time in Fig. 1D. (H) MotL band 1:2 unfolded torus in nonseizure (left) and seizure (right) states from 5 seizures. Color as in G. Gray line indicates slope 2 seizure trajectory at phase-lag m. Contrast SWD-spike (blue region) regularity with Fig. 1D. (I) Phase-difference histogram as in H from all mice showing a phase-lag shift m and increased coherence r from the non-seizure to seizure state, paired t-tests. Gray circle indicates mean phase-lag m. (J) SomL band 1:2 interaction schematic (left) and phase histogram (right) showing increased coherence r from the non-seizure to seizure state, paired t-test. Seizure mean phase-lag m (gray circle) differs from MotL (I) # p<0.05, t-test n=5 mean phase-lags, 1 per animal. (K) Overlaid example MotL and SomL Ecog traces with variable SWD-spike timing delays (left). Histogram (right) of MotL-SomL SWD-spike timing delays. Gray circles indicate individual mouse time-lag averages (−8.2 ms, one-sample t-test vs 0), purple (upper: 3.3 ms) and red (lower: −19.6 ms) bounds of 95% range of SWD-spike time-lags. (L) MotL:SomL band 2 interaction schematic (bottom) and example band 2 traces (top) from K. (M) MotL:SomL band 2 torus, color as in H. Dashed lines indicate lower (~-π/2 rad, red) and upper (~+0, purple) phase-lag bounds containing 95% of phase-lags and mean phase-lag m (~-π/4 rad, gray). (N) Phase-difference histogram, showing shifted mean phase-lag m and increased coherence r in the seizure vs non-seizure state, paired t-tests. Dots indicate mean phase-lag (gray) and 95% range of phase-lag distribution respectively as in M, right. Phase-lag differed from 0 in the seizure state (#, one-sample t-test vs 0). Note similarity with non-zero SWD-spike time-lags in K. (O-R) SWD-spike time-lags (O and Q) showing SWD-spike time-lags ~0 ms for MotL-MotR (n.s. one-sample t-test vs 0) and SomL-SomR (n.s. one-sample t-test vs 0). Gray, purple, red circles as in K. 95% range reduced in P relative to K, t-test. Corresponding band interaction schematics and phase-difference histograms (P and R) showing ~0 mean phase-lags m, n.s. one-sample t-test vs 0. There was increased coherence r but not phase-lag m in the seizure vs non-seizure state, paired t-test. Note similarity between 0 rad phase-lags to 0 ms time-lags in O and Q. (S) Schematic SomL/SomR band 1 interaction (left) and corresponding phase-difference histogram (right) showing no difference in phase-lag m or coherence r between seizure and non-seizure, paired t-tests. Histograms contain observations from all mice, N=5 mice, 5 seizures/mouse. Statistical comparisons were performed with values computed per mouse from 5 seizures/mouse.

While the fundamental SWD frequency is ~7.5 Hz, the timing of the SWD-spike (Fig. 2C, black arrows) aligned more strongly with the trough of band 2 than band 1. The trough of band 2 is at π rad by definition and the SWD-spike occurred near the trough in both MotL (2.97 ± 0.02 rad) and SomL (3.10 ± 0.04 rad). To quantify how reliably SWD-spikes are reflected by the phase of the different bands, we compared the circular standard deviation of the phase where SWD-spikes occurred. Standard deviation was 43% and 34% greater (i.e. more variable) for band 1 than 2 in SomL and MotL respectively, indicating tighter locking of SWD-spike timing to band 2 phase over band 1 phase (Fig. 2D, Fig. S2A). Similarly, band 3 standard deviation was 37% greater than band 2 in MotL, indicating tighter locking of SWD-spike to band 2 than band 3, while SomL band 2 and 3 were not different (Fig. S2A). The unfiltered Ecog trace (Fig. 2C black trace) also exhibited an additional peak and trough (gray arrows) between the SWD-spikes which also aligned with band 2 peaks and troughs (Fig. 2C blue and orange traces, gray arrows).

The substantial association of SWD-spike timing across bands suggests understanding both within- and between-region oscillation phase relationships is necessary to understand seizure dynamics and their origin. A standard method to examine how two oscillations evolve relative to each other, is to observe their joint dynamics on a phase torus. Fig. 2E shows a schematic of an arbitrary phase trajectory (red) on a torus defined by two oscillation phases, θ_i_ and θ_j_, where the phase torus is unfolded into a square with a periodic wraparound in both directions. Mapping MotL voltages onto the MotL band 1 phase versus band 2 phase torus (connection schematic Fig. 2F), revealed that SWD occurrences are confined to a specific region of the phase torus across many SWD cycles. For example, the dark blue SWD-spike troughs Fig. 2G, top, occur primarily at a specific point on the torus (Fig. 2G, bottom).

More generally, confinement of the entire trajectory along a limited region on the phase torus (Fig. 2G) suggests the presence of a dynamic attractor. To anticipate the quantitative interpretation of seizure dynamics in the toroidal coordinate system, we outline several theoretical possibilities and how to quantitatively detect them. For two oscillator phases (θ_i_, θ_j_), identical oscillation frequencies produce a slope-1 trajectory that repeats indefinitely, while differing frequencies cause diverging trajectories, filling the torus for irrational slopes (i.e. irrational oscillation frequency ratios, Fig. S2B). This stability of phase relationships is captured by phase-difference histograms, quantified by the coherence r ranging from 0 to 1 (Fig. S2C, Supplemental Methods, Equation 3). High coherence implies tight phase locking and narrow phase-difference histograms, while low coherence indicates the opposite. Note when computing phase-difference histograms and coherence for two oscillators with integer frequency ratios (e.g., 1 or 2), we first multiply the phase of the slower oscillator by the frequency ratio, otherwise the phase locking is not reflected (Fig. S2D and E). Finally, a constant phase difference or phase-lag between two phases reflects an offset of the trajectory without affecting slope, and it is captured by the mean m of the histogram of joint phase-differences (Fig. S2F and G).

Seizure trajectories exhibited numerous band- and region-specific changes in phase-lag and coherence when transitioning from non-seizure to seizure states. For example, plotting five seizures on the unfolded MotL band 1-2 torus as in Fig. 2G, revealed a lack of association between phases outside of seizures (Fig. 2H, left) but confinement to a narrow region within seizures (Fig. 2H, right). Thus, plotting voltage as a function of band 1 and band 2 phases revealed stereotyped structure in seizure dynamics that is not apparent in the temporally diverging trajectories shown in Fig. 1D and E. In particular, during seizures, dynamics condensed along a narrow slope-2 path (gray dotted trajectory), with SWD-spikes (blue) clustered in a specific torus region (Fig 2H, right). A circular histogram of 2θ_1_ − θ_2_ from all mice (Fig. 2I) revealed increased coherence and a shifted phase-lag from the nonseizure to seizure state (Fig. 2I gray circle). The SomL band 1-2 torus also showed increased coherence and a shifted phase-lag (Fig. 2J gray circle) from the non-seizure to seizure state, with a SomL phase-lag that was distinct from the MotL phase-lag. These examples represent a small subset of within-brain-region but between-frequency-band phase relationships, showing that within-region phase-lags have coherences and mean phase-lags that vary substantially across Mot (L/R) and Som (L/R) region pairs (schematic in Fig. S2H, left; histograms for bands 1:2, 1:3, and 2:3 across regions in Fig. S2I).

Overall, these results show that while there is significant temporal drift of SWD-timing when comparing different seizures (Fig. 1D and E), the toroidal coordinate system captures strong regularities in SWD waveforms across many oscillation cycles (Fig. 2F-J).

### Projection onto a toroidal coordinate system describes between-region SWD time-lags

Fig. 1G and H showed robust correlations between-regions during seizures. To examine how well the toroidal coordinate system captures between-region oscillatory features, we compared phase-lag measures in the toroidal system to a specific between-region feature of SWDs: the time-lag of SWD-spikes between-regions. The tight within-region locking of the SWD-spike time with band 2 phase in Fig. 2D suggested that between-region band 2 phase-lags may distinctly inform the SWD-spike time-lags. Indeed, with our toroidal coordinate system we found a strong quantitative correspondence between SWD-spike time-lags and the corresponding band 2 phase-lags. Individual MotL-SomL SWD-spike lags were strongly correlated with the corresponding phase-lags (Corr=0.70) between MotL and SomL band 2 (MotLθ_2_-SomLθ_2_, schematic and example traces, Fig. 2L). Bands 1 and 3 were more-weakly correlated (Corr = 0.45 and 0.46 respectively) (Fig. S2J and K). Additionally, there was a tight correspondence between the mean and ranges of SWD-spike time-lags with the mean and ranges of band 2 phase-lags. Mean MotL-SomL SWD-spike time-lag was −8.2 ± 0.9 ms, indicating SomL spikes typically occur before MotL spikes (example traces Fig. 2K, left and mean time-lags Fig. 2K, right gray circles). Since band 2’s frequency is 15 Hz (a period of 67 ms), SWD-spike time-lags of −8.2 ms divided by the period of 67 ms times 2π, should correspond to a phase-lag of −0.77 ± 0.1 rad or approximately −π/4 rad. Similarly, the 95% range (~two standard deviations) of SWD-spike time-lags between −19.6 ms to +3.3 ms (purple and red circles respectively, Fig. 2K, right), corresponds to band 2 phase-lags ranging from −1.84 rad to + 0.31 rad, or equivalently from slightly less than −π/2 rad to slightly more than 0 rad. As predicted, the directly measured MotLθ_2_-SomLθ_2_ phase-lag was near −π/4 rad (−0.80 ± 0.10 rad) during seizures, with a 95% interval between −1.86 ± 0.29 rad and 0.27 ± 0.15 rad (gray, red, and purple trajectories and corresponding circles in Fig. 2M and N). We will see below that these values correspond to specific dynamical drives of absence seizures and region-specific burst firing at the single neuron level. As observed above for within-region band 1 and 2 tori (Fig 2F-I), the MotLθ_2_-SomLθ_2_ trajectories were confined to a specific torus region, here confined to the 95% phase-lag interval described above (Fig. 2M and N red and purple trajectories). Moreover, the MotLθ_2_-SomLθ_2_ phase histogram captured seizure vs. non-seizure features including increased coherence during seizure relative to nonseizure and a phase-lag shift to near −π/4 rad (−0.80 ± 0.10 rad) from the near 0 rad (0.09 ± 0.03 rad) phase-lag outside of seizure (Fig. 2N).

We observed a similarly tight correspondence between band 2 phase-lags and SWD-spike time-lags in SomL-SomR and MotL-MotR (Fig. 2O-R, Fig. S2K, see calculations in Supplemental Methods). In contrast to MotL-SomL, mean SWD-spike time-lags in MotL-MotR and SomL-SomR showed no timing delay between either transcallosal pair during seizures (MotL: −0.7 ± 0.4 ms and SomL: 0.42 ± 1.05 ms, Fig. 2O and 2Q Gray circles). This was correspondingly reflected by a ~ 0 rad mean phase-lag in band 2 for MotLθ_2_- MotRθ_2_ (−0.07 ± 0.05 rad, Fig. 2P schematic and histogram) and SomLθ_2_-SomRθ_2_ (0.07 ± 0.11 rad, Fig. 2R schematic and histogram). Coherence increased from the non-seizure to seizure state in both transcallosal region pairs with no change in mean phase-lag (Fig. 2P and 2R). In contrast to band 2, the SomLθ_1_-SomRθ_1_ band 1 phase difference showed no change in mean phase-lag or coherence from seizure to non-seizure (Fig. 2T), indicating a lack of substantial changes during the seizure state in band 1.

Overall, these are examples of one particular SWD feature (the SWD-spike time-lags) and the relation to a small subset of all possible phase-difference relationships which have numerous band- and region-specific changes (schematic in Fig. S2H; histograms Fig. S2I and L). The switch to a toroidal coordinate system provides a rich quantitative phenomenology of mean phase-lags and coherences between frequency bands and regions with accompanying amplitude variations that determine the relative timing and waveform of SWDs. The large number of possible phase-difference and amplitude relationships, make it extremely difficult in general to understand individual pairs of oscillatory relationships and their causes in isolation. However, when considered in aggregate, they form a powerful set of metric constraints for identifying dynamical models that evolve to a putative absence seizure attractor described by these relationships.

### Identifying a sparse dynamical system model that recapitulates seizure dynamics

To discover a dynamical mechanism capable of generating these numerous complex relationships, we applied a system identification approach: Sparse Identification of Nonlinear Dynamics (SINDy).^26^ SINDy expresses how the system evolves toward future states as a function of the current state (Supplemental Methods, equation 4), selecting model terms from a candidate function library (Supplemental Methods, equation 5). Model complexity is limited by using sparsity-promoting regularization (see Supplemental Methods). We focused on modeling the earliest SWD onset regions observed in Fig. 1, limiting the model to bilateral exemplars of the motor and somatosensory systems: MotL (PreMotL), MotR (PreMotR), SomL (FaceL), and SomR (FaceR). We learned a model that tracks the real (x) and imaginary (y) components of the Hilbert transform of the 3 frequency bands as in (Fig. 2B) across 4 regions, forming a 2*3*4=24 dimensional dynamical system.

We trained the model on a single-step prediction task using data from 400 ms before to 800 ms after seizure onset using 25 seizures from each of 5 mice. We then tested its ability to simulate 3 s of seizure activity from 25 different pre-seizure initial conditions (5 from each mouse) that were not in the training set. This held-out test performance was evaluated by errors in predicting multi-second amplitude dynamics across 3 bands × 4 regions (12 metrics, Supplemental Methods equation 6), phase-lag (Supplemental Methods equation 7) and coherence (Supplemental Methods equation 8) across bands-within regions (2* 4 regions * 3 frequency pairs = 24 metrics), and within-bands across 4 region pairs: 2 transcallosal (MotL–MotR, SomL–SomR) and 2 ipsilateral somatosensory motor (MotL–SomL, MotR–SomR), yielding another 24 metrics (Fig. S2H). Together, these 60 metrics provide a strong basis for assessing how well our model recapitulates the state space of held-out seizure dynamics over seconds long timescales of future state prediction. We define an overall error metric composed of the average of these 60 metrics to quantify model performance (Supplemental Methods equation 9).

### The identified system of dynamic equations defines a chaotic absence seizure attractor

Using these metrics to guide model selection (see Supplemental Methods), our model closely reproduced experimental data across regions, capturing both voltage waveforms (Fig. 3A) and toroidal dynamics (example band 1–2 torus dynamics shown in Fig. 3B). The model reproduced seizure features even outside the toroidal coordinate system, with principle component analysis (PCA) projections of experimental and simulated data showing strong similarity across the first six components, which explain 90.1% and 91.3% of total variance, in experiment and the model, respectively (Fig. 3C). These components contained oscillatory trajectories progressing from low-amplitude pre-seizure states (blue) toward a common region in PC space that is consistent between experimental data and the model (Fig. 3D).

**Figure 3.**
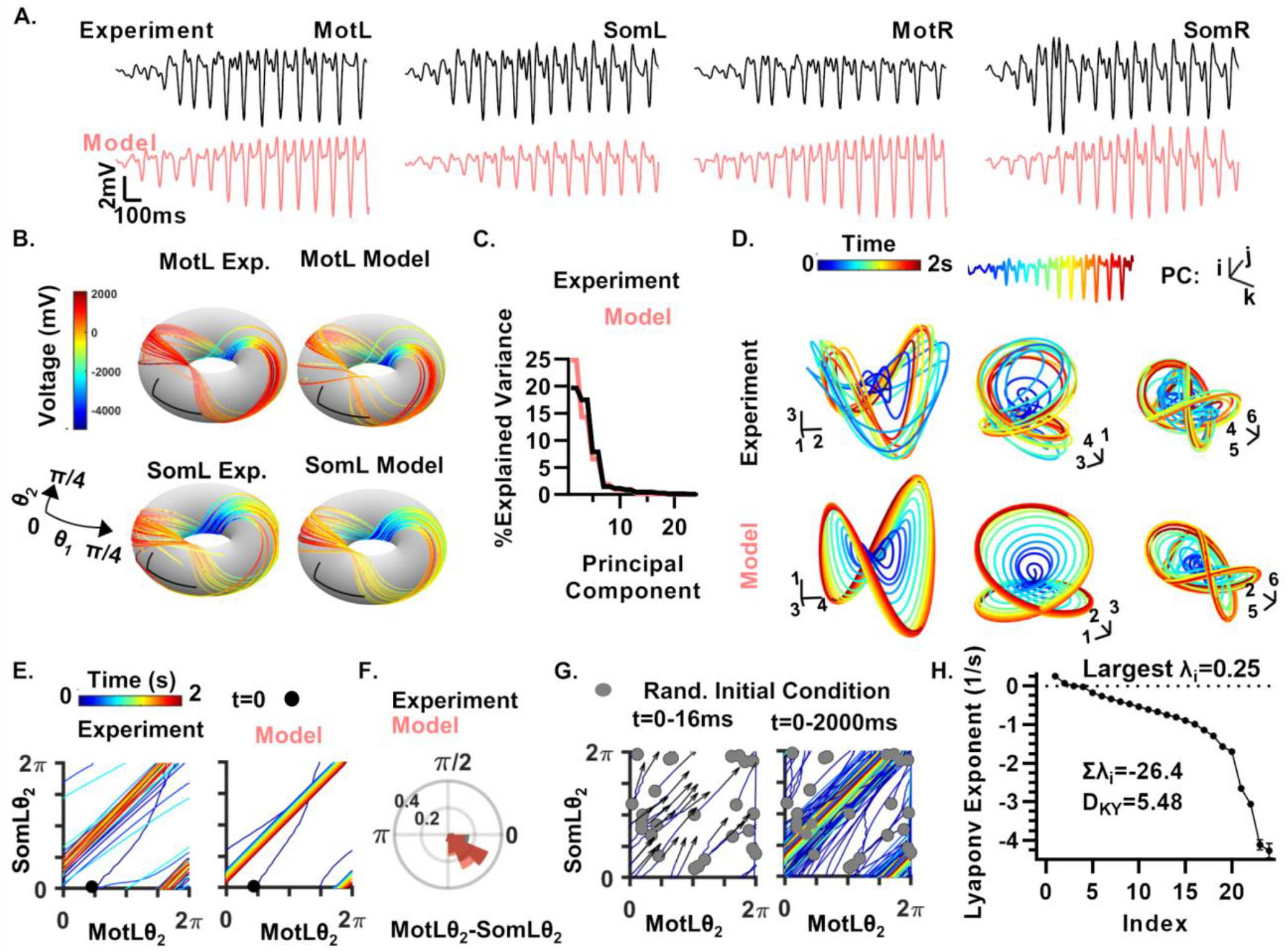
Sparse identification of nonlinear dynamics revealed governing equations for an absence seizure chaotic attractor. (A) Experimental and simulated voltage traces during seizures. (B) Experimental and simulated dynamics on band 1:2 tori for MotL and SomL. Color indicates voltage as in Fig. 2G. (C) Percent variance explained by principal components (PCs) in experiment and model. (D) Example PC traces showing model simulation (bottom) and experimental (top) dynamics across the first six PC components; trajectories colored by time. Scalebars 1000 μV, numbered by PC number. (E) Time-colored trajectory on the band 2 Mot/Som torus from an initial condition outside the seizure state space. (F) Phase-difference histogram comparing model and experiment across 25 pre-seizure initial conditions as in E. See Fig. S3 I and L for all between-band and between-region comparisons. (G) Simulations from 25 random initial conditions, initially disorganized during the first 16 ms (left) converge to the seizure state space (right); trajectories colored by time as in E. (H) Lyapunov exponent spectrum from simulations as in Fig. S3F. Largest exponent λ>0 indicating a chaotic system. Sum of λs being negative indicative of an attractor. Kaplan–Yorke dimension (D_KY_) of 5.48 indicates the attractor is a fractal occupying a between 5 and 6 dimensional subspace.

We next examined trajectories toward the seizure state space from the pre-seizure initial conditions. An example torus of band 2 MotLθ_2_ and SomLθ_2_ phases is shown in Fig. 3E; both experimental and model trajectories followed a similar curved path towards the seizure state space, afterward following straight-line trajectories as described in Fig. 2M, right. The corresponding phase-difference histogram showed high agreement between model and experiment, both in mean phase-lag and coherence (Fig. 3F). Our model performed well across 60 such phase and amplitude metrics (Fig. S3A-D). We will see below that the interpretable nature of the model allowed us to uncover a mechanistic basis for this and other phase-difference relationships. Total model error did not increase in simulating data from within the training set (0.19) vs held out test data (0.19). Remarkably, the model aligned so closely with experiment that replacing the model predicted future dynamics of seizures from an individual mouse with seizures from another experimental mouse yields higher total error (0.24) than simulation from the specific pre-seizure state (0.19), an effect reflected across phase and amplitude metrics (Fig S3C and E). Thus, our model describes seizure properties with greater accuracy than the level of between-subject variability in these properties.

Moreover, late-time seizure dynamics indicated an attractor state. Random initial conditions (real and imaginary components sampled between ± 150 μV) all converged to a limited seizure state space. An example torus is shown in Fig. 3G during the first 16 ms (left) and after 2 s (right). Replacing seizure-specific initial conditions with random initial conditions increased prediction error only slightly from 0.19 to 0.23, indicating specific pre-seizure states, while tracked, were not necessary for reaching the attractor state.

To assess long-term stability, we analyzed 1200 s long model simulations which showed bounded oscillatory SWD dynamics (first 25 s in Fig. S3F). A recurrence plot of the final 120 s (Fig S3G) revealed strong aperiodicity across long timescales, indicating that the system does not settle into a simple limit cycle oscillation. However, on short time scales on the order of the SWD-spike interval of 133 ms, the model exhibits approximately periodic oscillations with cycle-to-cycle variations in amplitude and phase that differ across regions. This strong aperiodicity over long time scales suggests the learned dynamical system exhibits chaotic dynamics within an attractor region. We confirmed this by computing the Lyapunov exponent spectrum which indicated a 5.48 dimensional (Kaplan-Yorke dimension: 5.48), chaotic (largest exponent: 0.25/s), and dissipative system (sum of exponents= −26.43/s), indicating the presence of a chaotic attractor (Fig. 3H).

Overall, these findings show that our dynamical equations, learned directly from data both: (1) quantitatively account for 60 different metrics characterizing held-out seizure dynamics, not in the training data, at a high level of accuracy, comparable to that of inter-subject variability between individual mice, and (2) reveals that absence seizure dynamics are well modelled by a dynamical system containing a 5.48 dimensional chaotic attractor.

### Our dynamical systems model for absence seizures is highly sparse and interpretable

A priori, the final dynamical system could have been highly complex, with 1134 candidate functions across 24 variables (27,216 parameters). However, our iterative expansion and sparsification method of model selection yielded a much smaller model (see Supplemental Methods). The model was built by iteratively expanding simpler candidate-function libraries and incrementally testing terms for removal (Schematic: Fig. S4A). Several quantal drops in prediction error for our 60 seizure metrics occurred with the addition of specific libraries (Fig. S4B). In contrast, single-step cross-validated mean squared error poorly distinguished between libraries (Fig. S4C). This dichotomy reveals the fundamental discriminative power of our 60 seizure state-space metrics in performing model selection, rather than simply following the widely used method of single-step prediction performance. In particular, improvements in specific metrics appeared at different library stages - e.g., autonomous, between-band, or between-regional terms (Fig. S4D).

Of 25 possible function libraries, only 11 contained selected (i.e. non-zero) terms. Each band/region included 43 terms on average. Many identified coefficients were near-identical multiples (see Supplemental Methods), enabling an analytic expression in the complex variable z=x+iy and conversion to polar coordinates z=Ae^iθ^. This left only 2 classes of autonomous terms (Fig. 4A), 2 classes of within-regional between-band interaction functions (Fig. 4B, left and middle), and 6 classes of between-regional interaction functions (one is shown in Fig. 4B, right and five are shown Fig S4E). The terms that most effectively improved model performance (see Fig. S4B) were in libraries 1, 2, 3, and 7, corresponding to the 5 functions shown in Fig.4A and B, while many between-region coupling terms shown in Fig. S4E (accounting for 46% of model terms) provided minor improvements individually, and removing them all increased the overall error metric slightly from 0.19 to 0.21 (Fig. S4F).

**Figure 4.**
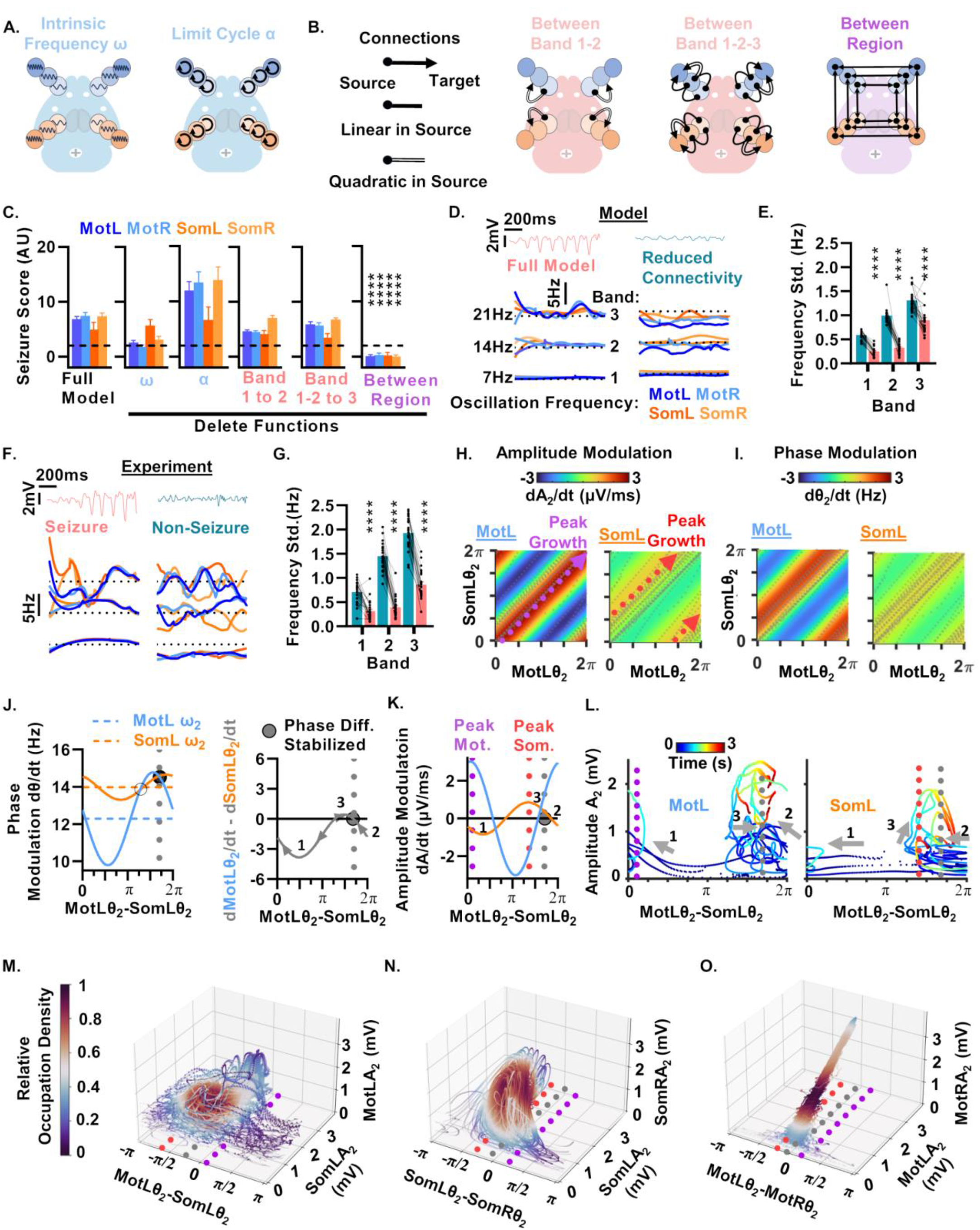
Between-regional coupling drives seizure generation through frequency locking driven amplitude growth. (A and B) Schematics of dominant model terms. (A) Band- and region-specific autonomous terms combine analytically to form oscillators with intrinsic frequency ω and amplitude-damping limit cycle α. (B) Coupling terms include between-band-within-region band 1 to 2 coupling (left), band 1 and 2 to 3 coupling (middle), and linear between-regional-within-band coupling (right). See also Fig S4E for additional terms and Supplemental Methods for analytic simplification steps. (C) Effect on seizure score of removing term classes from A and B by setting learned coefficients to 0. Asterisks indicate significant reductions below seizure score = 2 (average amplitude growth < 2 z-score from baseline). (D and E) Representative voltage traces (D, top), oscillation frequency (D, bottom), and (E) between-region oscillation frequency standard deviation (SD) for the full model (left) versus 6× reduced connectivity strength (right); *n* = 25 simulations from 5 mice pre-seizure initial conditions. (F and G) Experimental data as in D and E during seizures versus non-seizures (*n* = 25 seizures, *N* = 5 mice). (H and I) Example between-regional amplitude (H) and phase (I) coupling functions plotted on the torus showing phase and amplitude modulation along an experimentally observed seizure trajectory (gray dashed line). See Fig. S5D-G for other coupling functions. Red and purple dotted lines mark peak amplitude growth trajectories for SomL and MotL respectively. Note correspondence to upper and lower observed phase-difference bounds in Fig. 2M. (J) Analytically simplified phase coupling function showing phase modulation of band 2 by phase difference frequency modulation around respective intrinsic frequencies (ω, dashed lines); stability points marked at intersections (circles). Phase-lag stabilization (gray dashed line, J, right) occurs where the difference between curves cross from positive to negative between regions marked 2 and 3. Region marked 1 causes strong decrease in phase-lag, leading to wraparound to high phase-lags at point 2. Note similarity of stability point to mean phase-lag in experiment (Fig. 2N) and model (Fig. 3F). (K) Amplitude modulation showing peak growth for Som (red dashed line) and Mot (purple dashed line) band 2 as in H. Note similarity to phase-lags of the 95% interval experimentally in Fig. 2M and N. (L) Five example trajectories of band 2 amplitude and phase-difference colored by time. Dynamics cycle around phase stability point marked by gray dashed line. (M-O) Examination of band 2 trajectories along the chaotic attractor. (M) Dynamics as in L for long simulations. (N) Dynamics of SomL-SomR. (O) Dynamics of MotL-MotR. See Fig. S5H for MotR-SomR. 95% ranges from Fig. 2NPR indicated by red and purple lines and mean phase-lag by gray. Statistical analyses were performed by one-sample (C) or paired Student’s *t* test (E and G); *N* = 5 mice, *n* = 25 simulations or seizures per condition. Data represent mean ± SEM.

The sparse nature of our selected terms, obtained through model selection, yielded a mechanistically interpretable model. For each region and each band i=1,2,3, we denote the phase by θ_i_ and the amplitude by A_i_. Here autonomous dynamics (Fig. 4A) are defined via the dynamic equations dθ_i_/dt = ω_i_ and dA_i_/dt = α_i_A_i_^3^, with α_i_ <= 0. This describes an oscillator with intrinsic frequency ω_i_ (Library 1, Schematic: Fig. 4A, left), whose amplitude A_i_ decays to zero via a cubic nonlinearity (Library 1, Schematic: Fig. 4A, right). Since in the data-driven model, each individual band in each region is stable in isolation, the generation of large amplitude epileptic oscillations solely relies on between-band or between-region interactions. Each such interaction term involved phases and amplitudes of oscillator pairs or triplets. Functions included quadratic band 1 to 2 between-band, within-region terms (Library 2, Fig. 4B, left); within-region cubic band 1 and 2 to 3 terms (Library 3, Fig. 4B, middle); linear between-region, within-band terms (Library 7, Fig. 4B, right); and several nonlinear between-region, between-band interactions (Libraries 15– 21, Fig. S4E). These latter combined between-band, between-region functions contributed little to model accuracy (Fig. S4F). See Supplemental Methods for a mathematical description of all of these interaction terms.

In summary, from 27,216 potential parameters, our model selection identified an accurate and interpretable sparse dynamical model generating the dynamics of absence seizures, with only an average of 43 nonzero parameters per oscillator that are highly organized into interactions between bands and regions with different polynomial orders that can be combined analytically into a small number of interpretable functions.

### Strong between-region coupling and limited frequency dispersion are required for seizure generation

To identify which classes of function drive seizures, we iteratively set each class’s terms to zero in the learned model. Seizures could only be abolished by removal of between-region within-band coupling (Fig. 4B, right, Fig. 4C, Fig. S4G). This suggests that while other terms contribute to model accuracy, the *large-amplitude seizure oscillations are driven exclusively by between-region coupling*. We then swept the weights of between-region within-band coupling between 0 and 1 and found that a six-fold reduction is required to abolish seizures (Fig. S4H). Doubling all terms within each individual function class did not suppress seizures for any function class (Fig. S4G), suggesting that no individual function class strongly promotes damping of seizures.

The central importance of between-regional coupling suggests that synchronization between regions plays a fundamental role in seizure generation. To test this hypothesis, we increased the spread of intrinsic oscillator frequencies in the model by shifting each one independently (see Supplemental Methods). Such dispersal of intrinsic oscillator frequencies is known theoretically to suppresses synchronization.^27^ See example shifts for band 1 in Fig. S4I. Indeed, seizure scores decreased across all regions with increased frequency spread between regions (Fig. S4J). Thus, as predicted, suppressing synchronization by dispersing intrinsic frequencies between regions prevented seizures.

Also consistent with the hypothesis that synchronization plays a key role in seizure generation, we found that each region and band converged to a common band-specific oscillation frequency (Fig. 4D). Furthermore, reducing between-region connectivity strength sixfold eliminated both seizures (Fig. 4D, top-right) and frequency convergence (Fig. 4D, bottom-right), as quantified by frequency standard deviation between regions (Fig. 4E). Similarly, increasing intrinsic frequency spread (Fig. S4I) disrupted frequency convergence (Fig. S4K, L).

Thus overall, strong between-regional coupling and limited intrinsic frequency spread are two key features leading to seizure generation in the model; if either frequencies are dispersed or coupling is reduced, both seizures and synchronization are abolished, as is expected in theories of synchronization phase transitions in coupled oscillators.^27^

### Frequency locking, as predicted by the model, is confirmed in experimental Ecog data

Our model predicted that between-region frequency spread drops to near zero at seizure onset. We confirmed this prediction in experimental data in which each band transitions from dispersed frequencies across regions to a single common frequency across regions (Fig. 4F, left). In contrast, outside of seizures (Fig. 4F, right), oscillator frequencies varied widely across regions (Fig. 4G). The non-seizure frequency spread (Fig. 4G) resembled that of the reduced-connectivity model (Fig. 4E). In contrast, the degree of frequency dispersion required to block seizures in the model (Fig. S4J) far exceeds the experimental observations in the non-seizure state (>4-fold, Fig. 4L). This suggests that the transition from non-seizure to seizure states in the brain originates from an increase in effective between-regional connectivity strength, rather than a decrease in intrinsic oscillator frequency dispersion.

### Synchronization and phase-lag dependent amplitude growth drive seizure generation

We next tested whether specific bands or couplings were required for seizures. Robust seizures persisted when the model was reduced to only the intrinsic oscillatory terms - intrinsic frequency *ω*, cubic damping *α*, and the seizure-driving between-regional, within-band coupling (Fig. S5A). This shows that between-region coupling is not only necessary but also sufficient to drive seizures in the model. Further reduction of the between-region coupling to only Mot–Som or transcallosal connections reduced but did not fully eliminate seizures (Fig. S5A). In the full model, seizures persisted when any single band or all Mot– Som or transcallosal couplings were removed in isolation (Fig. S5B and C). Thus, no specific band or regional coupling is essential. In essence all within-band, between-regional couplings collectively contribute to seizure generation (Fig. 4C).

To gain insight into the mechanisms of between-regional seizure generation, we next examined the form of these crucial between-region, within-band couplings (Library 7, Fig. 4B, right). Given the strong between-region frequency locking of band 2 in both the model (Fig. 4E) and the data (Fig. 4G), as well as the strong association of band 2 with the SWD-spike (Fig 2C-D) and between-region SWD time lags (Fig. 2K-S), we focus here on between-region couplings within band 2. However, we also show several other between-region, within-band couplings in Fig. S5D-G, as seizure generation does not depend exclusively on band 2 coupling (Fig. S5B). These coupling functions, as above, could be converted to analytic expressions in the complex variable z=x+iy and converted to polar coordinates z=Ae^iθ^ resulting in functions of the following form given a pair of regions and a single band:

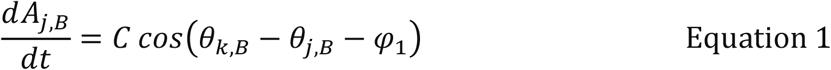

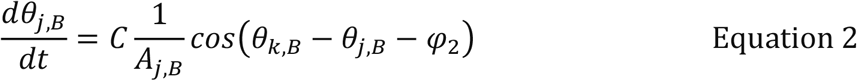

Here, A_j,B_ and θ_j,B_ denote the amplitude and phase respectively of target region j in band B and A_K,B_ and θ_k,B_ denote the amplitude and phase of a source region k (see Supplemental Methods). The constants φ_1_ and φ_2_ indicate preferred phase-lags in the specific source-to-target coupling. This form is followed by all between-region, within-band couplings, with transcallosal Mot couplings multiplied by an additional factor A_k,B_ in both equations. Remarkably, these couplings identified from only the real and imaginary components of experimental signals combined to resemble the paradigmatic Kuramoto model of synchronization of phase-coupled oscillators but with additional amplitude variation.^27^

Because seizures are defined by the growth of large amplitude oscillations from a low amplitude state (Fig. S1A-D), we first examined the form of the amplitude coupling functions which drive this amplitude growth. Equation 1 states that if the phase-lag *θ*_*k,B*_ − *θ*_*j,B*_ between the target region k and source region j is equal to the amplitude coupling’s preferred phase-lag φ_1_, then the target region amplitude A_j,B_ grows at a maximal rate. In contrast, if the phase-lag is the opposite at *φ*_1_ ± *π* then the amplitude decays at a maximal rate. Fig. 4H plots the band 2 amplitude coupling functions (Equation 1) from SomL to MotL (left) and MotL to SomL (right) as joint functions of the MotL-SomL phase-lag (i.e. on the phase torus). This revealed that the rate of amplitude growth occupies disjoint regions on the phase torus for MotL and SomL respectively. In particular, MotL amplitude growth occurs at a maximal rate when the MotL-SomL phase-lag is near zero (red region, purple trajectory in Fig. 4H, left). In contrast, in this same phase-lag region, SomL amplitude moderately decays (blue-green region in Fig. 4H, right). Conversely, when MotL-SomL phase-lag is slightly more negative than −π/2, SomL amplitude grows at a maximal rate (red region, red trajectory in Fig. 4H, right,), while MotL amplitude moderately decays (blue-green region of Fig. 4H, left). This result provides a hint that MotL and SomL amplitude growth will tend to be anticorrelated. We will see below that this anticorrelated growth corresponds to region specific bursting at a single neuron spiking level. Notably, these maximal growth trajectories correspond to the upper and lower bounds of the between region phase-lags (Fig. 2M and N), and thus between-region SWD-spike time-lags (Fig. 2K) observed experimentally.

Due to the dependence of amplitude growth on between-region phase-differences, we next examined the form of the between-region phase coupling functions. Fig. 4I shows band 2 phase coupling functions (Equation 2) from SomL to MotL (left) and MotL to SomL (right) as joint functions of the MotL-SomL phase lag. These coupling functions are shown at a fixed small amplitude of 200 μV for both regions, allowing us to understand phase modulation in the transition from the non-seizure to seizure state. In isolation, these functions do not explain how the MotL-SomL phase-lag stabilizes near a particular value in the seizure. To elucidate this, we plotted (Fig. 4J, left) the rate of phase advance (i.e. instantaneous oscillation frequency) arising from two sources: the region-specific autonomous intrinsic oscillation frequencies ω (dotted lines) and the sum of the intrinsic frequencies with the additional sinusoidal modulation arising from the between-region phase coupling (solid lines) between MotL and SomL band 2 (Equation 2). We examine the phase advance at fixed 200 μV amplitudes and discuss the amplitude variation effects below. The curves intersect (i.e. have equal oscillation frequency) at two points, with one notably near −π/4 rad (equivalently 3π/4, gray dotted line), corresponding to the mean phase-lag during seizures in experiment and model for MotL-SomL band 2 (Fig. 2M-N and Fig. 3F). The difference of the phase advance curves yields Fig. 4J, right, expressing the rate of change of the phase-lag between MotL and SomL as a function of the phase-lag itself. This plot reveals that this ~ −π/4 rad (= 3π/4) intersection is a stable fixed point at this fixed amplitude, with strongly decreasing phase-lag when at lower lags to the far left (arrow 1), leading to a wrap-around of phase-difference to higher lags to the right of the stable point (arrow 2), after which the phase-difference then continues to decrease towards the stable point. Just to the left of this stable point, the rate of phase-difference change switches directions, causing the phase-difference to now *increase* towards the stable point. Thus, the phase interactions between MotL and SomL (Fig. 4JK), at low fixed amplitudes, stabilizes their phase-lag at −π/4 rad, thereby explaining the mean phase lag observed in both model and experiment (Fig. 2N and Fig. 3F).

### Phase-amplitude coupling drives chaotic dynamics

The phase relationship at the above point would be a stable fixed point if amplitudes were clamped at 200 μV. However, the amplitude coupling functions show that at this phase-lag, there is a slight rate of amplitude growth in MotL *and* SomL band 2 (Fig. 4K, both orange and blue curves are positive at the fixed-point phase-lag –π/4 (= 3π/4) rad marked by the gray dotted line). Furthermore, due to the anti-correlated nature of MotL and SomL amplitude growth as a function of phase lag (Fig. 4H), we see in Fig. 4K that just to the right of the −π/4 rad phase lag (point 2), MotL amplitude grows faster and SomL amplitude moderately decays whereas to the left of this phase lag (point 3), MotL amplitude decays while SomL amplitude grows. Due to the 1/A_j_ scaling of the phase modulation curves, this causes the stable phase lag point to shift to the right towards 0 when SomL band 2 amplitude is larger and to shift to the left towards −π/2 rad when MotL amplitude is larger. Thus, in the combined MotL-SomL phase-amplitude feedback dynamics, the phase lag oscillates about −π/4 rad (gray dashed line) with the amplitudes growing and decaying (arrows 2 and 3) by oscillating between peak Mot (purple dashed line) and Som (red dashed line) growth at phase lags between 0 and − π/2 rad respectively while avoiding areas where both regions will decay (Fig. 4L). Overall, the reason the system grows to a net high amplitude (seizure) state is because in the narrow range over which the phase-lag oscillates (between −π/2 and 0 rad), the rate of change of amplitude is never strongly negative (Fig. 4K) and therefore cannot cause net decay to very low amplitudes. While SomL amplitude dynamics vary relatively smoothly as the phase lag oscillates about −π/4 rad indicated by arrows 2 and 3 (Fig. 4L, right), the MotL dynamics have additional variations in the high amplitude state (Fig. 4L, left), reflecting the additional transcallosal MotL-MotR coupling strength that grows multiplicatively with amplitude by A_k,B_.

In long simulations, the combined dynamics of phase-lag and SomL and MotL amplitude varied chaotically, with phase-lag oscillating, as expected, in the range of −π/2 and 0 rad (red and purple dashed lines respectively) about the mean overall phase-lag (~ −π/4 rad) (gray dashed line, Fig. 4M), as observed in *both* model *and* data (Fig. 2O, Fig. 3F). Brief destabilizations repeatedly returned to this basin. MotR–SomR showed similar behavior (Fig. S5H), while transcallosal Som coupling oscillated near 0 rad (gray dashed line) with repeated folding of the trajectory (Fig. 4N), reflecting the high coherence and 0 phase-lag observed in model and data (Fig. 2R, Fig. S3B). This is consistent with the structure of SomL-SomR couplings (Fig. S5F). Transcallosal Mot coupling varied in amplitude but was tightly locked near 0 rad (gray dashed line, Fig. 4O), again consistent with model and experiment (Fig. 4P, Fig. S3B) and the MotL-MotR coupling functions (Fig. S5G). Notably, the MotL-MotR couplings have phase and amplitude modulation growing with an additional factor of A_k_, explaining the tighter locking observed in model (Fig. S3B) and experiment (Fig. 2P). Altogether these results provide a comprehensive mechanistic understanding of the phase and amplitude relationships described in Fig. 2 and Fig. S2, which together define absence seizures as a chaotic dynamical attractor (Fig. 3).

Finally, while we have focused here on within-band between-region couplings, we analyze the complementary between-band, within-region couplings (see Supplemental Methods for these coupling functions). We show that these couplings only become strong at large amplitudes, thereby causing band 1-2 phase synchronization and SWD waveforms to appear in high amplitude seizure states (Fig. S5I and J), but not outside seizures as was observed in experiment (Fig. 2G-H. Fig. S2I). *Thus, SWD waveforms result from rather than cause the oscillation amplitude growth during seizures*.

Overall, these within-band, between-region coupling functions reveal a mechanism for seizure generation through synchronization-induced amplitude growth. Basically, each coupling function prefers a certain low-amplitude stable fixed point phase-lag between source and target oscillators. Arrival at this preferred phase-lag leads to amplitude growth. Then feedback coupling between amplitude dynamics and phase-lag dynamics induces a slow variation about the low amplitude stable fixed point phase-lag in the high amplitude state. Moreover, this variation appears to be chaotic with continued variations in amplitude and phase on the time scale of seconds, thereby suggesting a mechanistic origin for the overall chaos quantified in Fig. 3H. In summary, synchronization near the preferred phase-lag of pairs of oscillators induces amplitude growth in both oscillators. Below we will employ these model-derived explicit dynamical mechanisms for amplitude growth to uncover corresponding neuronal circuits and single neuron spiking patterns.

### Single neuron spiking in a Som-Mot-PO subnetwork frequency locks to the fundamental seizure frequency at seizure onset

We next examined whether single neuron spiking reflected Ecog features and model predictions. Using three simultaneous Neuropixels probes in awake mice, along with simultaneously recorded Ecog from surface and contralateral somatosensory electrodes (Fig. 5A), we mapped activity across brain regions during absence seizures. The Neuropixels probes were labelled with dyes DiI/DiD/DiO marking their position in the brain (Fig. 5A pink and yellow regions), and we identified single units through spike sorting (Fig. S6A-B). Together, this allowed alignment of spike rasters to Ecog SWDs in PreMotor and bilateral Som Face/Barrel cortex (Fig. 5B). Cortical units were classified as excitatory or inhibitory by waveform risetime (Fig. S6C). Mean firing rates showed only moderate region-specific changes (Fig. S6D) from non-seizure to seizure states. Notably, Som deep layers showed *decreased* overall firing (Fig. S6D, top-left), consistent with recent findings of no hyperexcitability in Som during or preceding absence seizures.^3–6^ These results suggest seizure organization depends more on spike timing and firing mode (burst vs. non-burst) than on overall firing rate.

**Figure 5.**
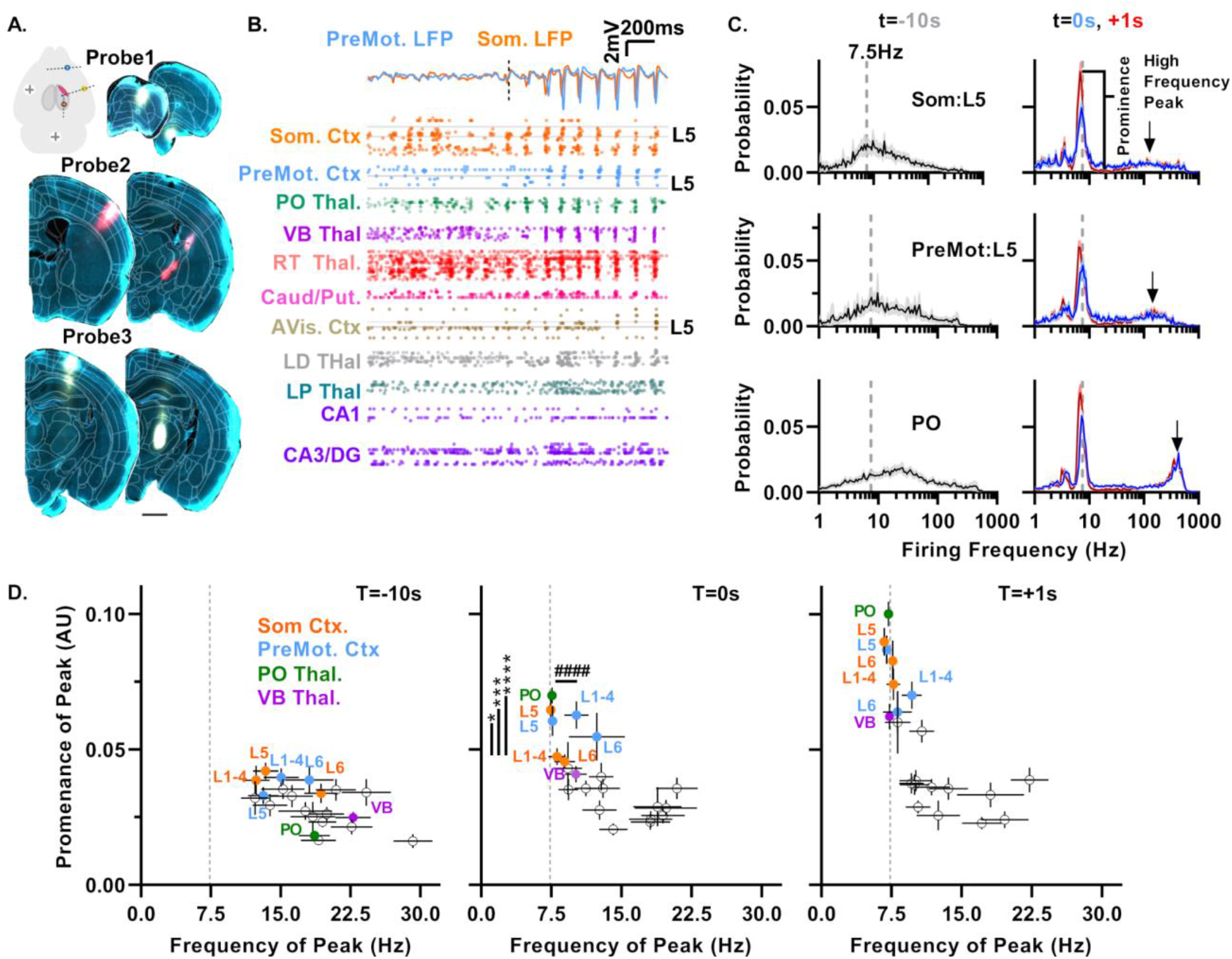
Neuronal population frequency locking identified Som:L5, PreMot:L5, and PO as an early seizure network. (A) Experimental design showing Neuropixels probe and SomL screw electrode placements for simultaneous Ecog and neuronal spiking recording. DiI/DiD-labeled probe tracts mark insertion points and maximum depths; white outlines denote region boundaries from SharpTrack alignment. Scale bar = 1 mm. (B) Simultaneously recorded spike rasters from single units across regions and Ecog in PreMot and Som Face regions during a single seizure. Black dashed line indicates seizure onset. Gray lines indicate L5 boundaries. Units marked by depth within region. (C) Instantaneous firing-rate histograms for Som:L5, PreMot:L5, and PO averaged across seizures (5 per recording, 6–15 recordings, from 4–8 mice), error bars indicate SEM. Time relative to seizure onset: −10, 0, +1 s. The window size of 2 s around these times. Prominence calculated as difference between peak and average left/right of peak 7 frequency bins away. (D) Scatterplots of frequency of peak vs. prominence. Dashed line marks the 7.5 Hz seizure fundamental frequency. At seizure onset (T=0), PO, Som:L5, and PreMot:L5 cluster at this frequency. Asterisks indicate significance for prominence for PO, Som:L5, PreMot:L5 vs Som:L1-4, the next closest to frequency locked population. Welch’s t-test. Hashtags indicate significance for frequency variance vs Som:L1-4 and PreMot:L1-4, both p<0.0001. At T=+1 s, PO, Som:L5, PreMot:L5 show highest peak prominence, with other regions converging to the fundamental.

Since our model predicted frequency locking, at the level of LFP across regions (Fig. 4DE) and we confirmed this prediction in Ecog data (Fig. 4EG), we next examined if frequency locking at the level of individual neurons would inform which networks drive this activity. We measured region-specific histograms of the reciprocal of interspike intervals (i.e. instantaneous spike frequencies) for all neurons in the region, obtained from 2 second time windows centered at different time points relative to seizure onset (nonseizure at −10s, onset at 0s, and +1s after onset corresponding to gray, blue, red histograms, respectively in Fig. 5C). We used peaks in the population’s spike interval distribution as indicators of frequency clustering, where concentrated spiking at a specific frequency produces an elevated count at that frequency and reduction at neighboring frequencies, quantified by peak prominence (Fig. 5C). We found no frequency locking in the nonseizure state as indicated by low prominence at the peak frequency (gray histograms in Fig. 5C, left, Fig. 5D left), but observed strong locking of firing frequencies to the fundamental seizure frequency near 7.5Hz at onset, which was sustained through the seizure in specific populations (blue and red histograms in Fig. 5C, right, Fig. 5D, middle, right). We also observed smaller isolated peaks at high frequencies > 40Hz (arrows Fig. 5C, right), indicative of bursting, which increased in both thalamus and cortex (Fig. S6F,G). The strongest spike frequency locking at seizure onset, (blue, Fig. 5C, right), occurred in somatosensory cortex layer 5 (Som:L5), premotor cortex L5 (PreMot:L5), and posterior thalamic nucleus (PO) (blue Fig. 5C, right, and Fig. 5D, middle). These regions all exhibited higher peak prominence than the next nearest population to the seizure frequency (Som:L1-4) (Fig. 5D). These populations also exhibited very low variance in the frequency of the peak at onset across seizures 0.34 (PO), 5.1 (SomL:5), 8.3 Hz (PreMot:L5) compared to next highest prominence region: (PreMotL:2-4, 77.2 Hz) and next population closest to the fundamental seizure frequency (SomL:L1-4, 32.9 Hz). This highly consistent frequency locking of PO, Som:L5, and PreMot:L5 indicates these regions are more reliably recruited to a common seizure frequency at onset than other regions (Fig. 5D). These regions also had the highest peak prominence after seizure onset (Fig. 5C, right, red, and Fig. 5D, right). Notably, other regions such as ventrobasal thalamus and the other Som and Mot cortical layers are strongly recruited to the locked state after onset (Fig. 5D, right).

These results support an *early seizure network linking Som:L5 and PreMot:L5 with PO*. Importantly, these three populations are highly interconnected and reflect the Mot and Som early seizure network indicated in Fig. 1C. *PO projects to the apical dendrites of layer 5 pyramidal neurons* in both of these regions and both L5 pyramidal populations are reciprocally connected.^28–30^ The recipient zone for connections from PO to these apical dendrites are located both in layer 1 and layer 5. Intriguingly, layer 1 apical dendrites are also the primary recipient of cortico-cortical feedback inputs from other cortical regions and prior work has found that PO thalamus boosts the effectiveness of input at these distal sites.^31, 32^ This anatomy then suggests that PO inputs could broadly modulate the strength of feedback coupling from Som:L5 to Mot layer 1 and Mot:L5 to Som layer 1 along with transcallosal connections in both regions during seizures.^33^ We next examine the relationship between LFP and single neuron spiking dynamics focusing on these three tightly locked interconnected regions.

### LFP amplitude growth predicted by model coupling functions is associated with single neuron bursting in the target region

We have seen above that LFP in band 2 strongly reflects between-region SWD-spike timing lags (Fig. 2K-R) and shows a strong increase in coherence from non-seizure to seizure states (Fig. 2NPR). Furthermore, we found that phase-lags between pairs of regions determine whether respective LFP amplitudes in each region grow or decay in the model (e.g. Fig. 4H for SomL and MotL, and S5D-G for other pairs of regions). However, single neuron spiking frequency locks near the fundamental band 1 frequency (7.5 Hz) (Fig. 5C), not the higher band 2 frequency (15 Hz). This raises a fundamental question of how band 2 phase-lags and thus ultimately the spike of the SWD relates to individual neuronal spike times underlying the seizure.

To address this, we examined the relationship between band 2 oscillations (Fig. 6A, top) in Som and Mot and the full spectrum unfiltered LFP containing the SWDs (Fig. 6A, middle) relative to single neuron spiking (Fig. 6A bottom). Recall that band 2 phase-lags between Mot and Som oscillate between 0 to −π/2 rad (Fig. 2N), which corresponds to a range of SWD-spike time-lags of ~ 0 to −15 ms (Fig. 2K). Moreover, note that a phase-lag of ~0 corresponds to peak amplitude growth in MotL (Fig. 4H, left) while a phase-lag of ~-π/2 rad corresponds to peak amplitude growth in SomL (Fig. 4H, right). We focused on Som:L5, Mot:L5, and PO neurons because these were the first to show strong single neuron frequency locking at seizure onset (Fig. 5D, middle). The temporal relationship of neuronal spikes to the SWD-spikes varied on a cycle-by-cycle basis (Fig.6A). The peristimulus time histogram (PSTH) of single-neuron spikes in Som:L5, PreMot:L5, and PO relative to the Som SWD-spike, revealed that the Som:L5 single unit spikes occurred preferentially −4 ms before the Som SWD-spike (Fig. 6B, blue) while PreMot:L5 single neurons preferentially fired +4 ms after (Fig. 6B, blue). Notably the time-lag between single neuron spikes closely resembled the time-lag of −8.2 ms between the SWD-spikes in these regions (Fig. 2K), reflecting time-lags found previously.^34^ PO single neuron spikes exhibited the highest peak, occurring at −6 ms before the Som SWD-spike (Fig. 6B, green). Notably, there was considerable overlap between these populations over a range of −15 to +15 ms, with PO exhibiting an earlier increase in spiking around −20 ms before the Som SWD-spike (Fig. 6B arrow 1) compared to Som:L5 or PreMot:L5 (both p<0.0001, Mann-Whitney U test), while Som:L5 and PreMot:L5 did not differ from each other at this timepoint. While the PreMot:L5 peak trailed the Som:L5 peak by 8 ms, there was a substantial shoulder (47% of the peak firing) of the PreMot:L5 distribution aligned with the Som:L5 peak (Fig. 6B arrow 2).

**Figure 6.**
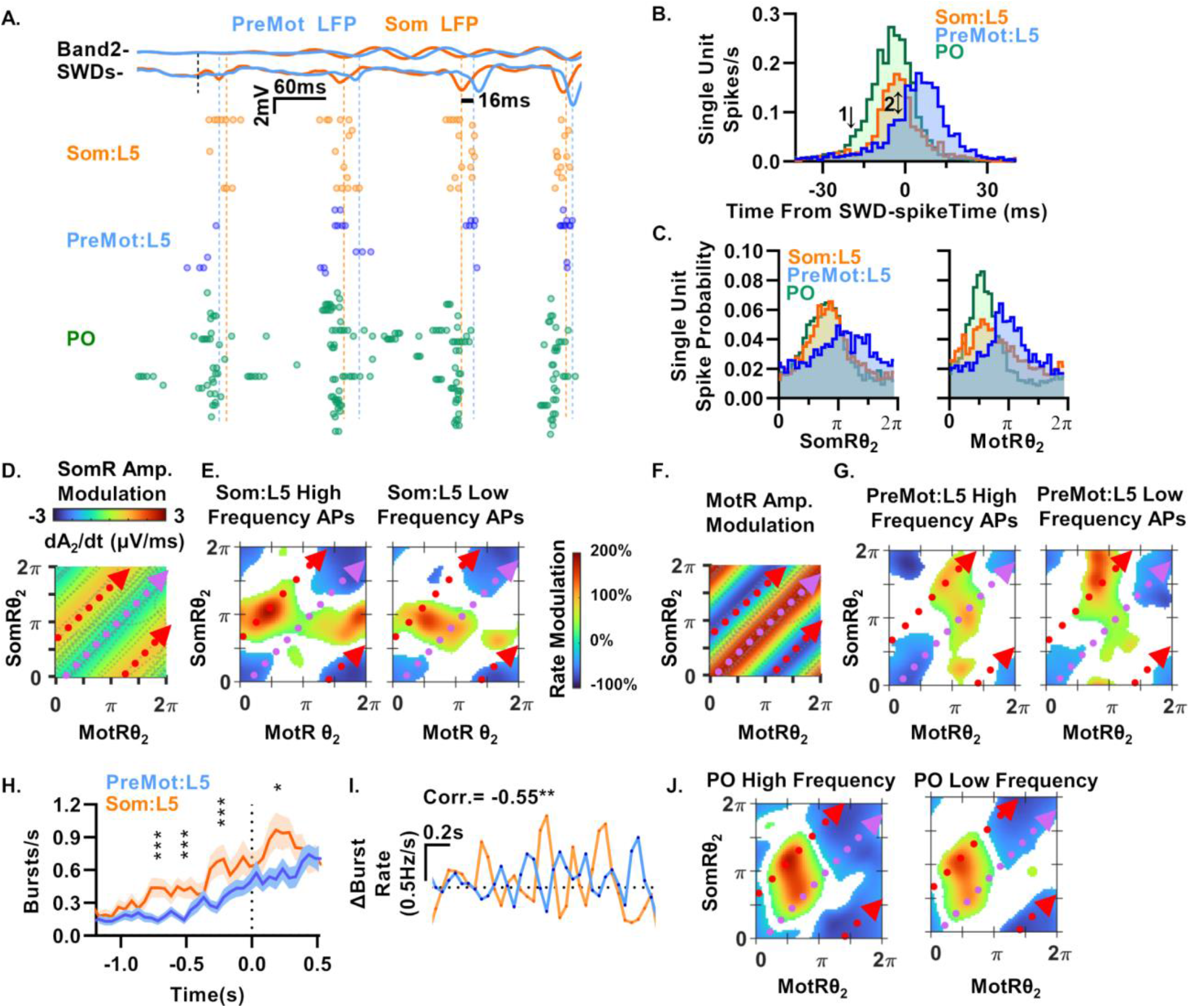
Amplitude coupling functions corresponded to region specific burst and tonic firing mode. (A) Expanded view of raster shown in Fig. 5B showing band 2 alignment (top) with full spectrum Ecog (middle) and unit firing in Som:L5, PreMot:L5, and PO. Orange and blue dashed lines mark trough of band 2, roughly aligning to the peak of the SWD-spike. Units sorted by unit ID. Note the variable separation of spiking clusters across regions on a cycle-by-cycle basis. (B and C) (B) Som SWD-spike triggered peristimulus time histogram. Arrow 1 indicates early elevation of PO firing above Som:L5 or PreMot:L5 (both p<0.0001, Mann-Whitney U test). Arrow 2 indicates PreMot:L5 shoulder overlap with Som:L5 firing peak. (C) Neuron-spike triggered phase histograms for band 2. (D) Amplitude coupling function of SomR, red dashed line marks peak SomR amplitude growth, purple dashed line marks peak MotL Amplitude growth, corresponding to SomR amplitude decay as Fig. 4H. (E) Som:L5 high-frequency firing modulation (left) and low-frequency firing (right) on the phase torus. Note peak amplitude growth trajectory (red line) traverses peak high-frequency firing (red region) followed by silencing (blue region) while amplitude suppressing trajectory (purple) traverses modest high-frequency firing (green region) (E, left) while low-frequency firing is elevated along both trajectories. (F) As in D for MotR modulation. Note opposing area of amplitude growth and decay from D as in Fig. 4H. (G) As in E for MotR high-frequency and low-frequency firing. Note MotR amplitude growth marked by purple trajectory traverses increase and decrease high-frequncy firing (left) while the amplitude suppression trajectory (red) traverses increased high-frequency and increased low-frequency firing without negative modulation. (H) Average number of detected bursts (a >40 Hz interval preceded by a <10Hz interval). Asterisks indicate significant differences between regions. (I) Correlation of change in burst rate from H showing changes in Som:L5 and PreMot:L5 burst rates are anticorrelated. (J) As in E and G for PO high-frequency (left) and low-frequency (right) modulation. Note the increase and decrease along both the Mot and Som amplitude growth trajectories.

The parallel between PreMot:L5-Som:L5 single neuron spike time lags (Fig. 6B) and Mot-Som SWD-spike time lags (Fig. 2K), together with the correspondence between SWD-spike time and band 2 troughs (Fig. 2N), suggest that the variable timing of single neuron spikes in these regions could be informed by band 2 phases. Single spikes in these regions were indeed modulated by Som:L5 and PreMot:L5 band 2 phases (Fig. 6C), but these phases in isolation did not clarify spiking relationships between regions. However, we unearthed a strong association between single neuronal spikes and LFP phases by examining when spiking occurs as a *joint* function of *both* Mot and Som band 2 phases, i.e. on the band 2 phase torus. This analysis is motivated by our model, since the joint phases of Mot and Som determine seizure amplitude growth in a manner not computable by knowing Mot or Som phases alone (Fig. 4H).

Fig. 6D shows the model SomR amplitude growth rate as a joint function of SomR and MotR phases. For reference, a hypothetical red constant phase-lag trajectory is shown to mark the region of maximal amplitude growth rate of Som as in Fig. 4H, indicated by the red regions at a MotRθ_2_-SomRθ_2_ phase lag of approximately −π/2 rad. Similarly, a hypothetical purple constant phase-lag trajectory is shown to mark the region of maximal Som band 2 amplitude decay rate, indicated by the light blue region at a phase-lag near 0 rad. As explained above (Fig. 4H-M) seizure trajectories (shown as dotted gray lines in Fig. 6D) wander between the hypothetical purple and red trajectories, leading to temporally varying Som and Mot amplitude growth and decay which leads to net amplitude growth in both regions.

We next plotted a 2-D histogram of spiking timing changes in SomR:L5 as a joint function of SomR and MotR phases. In particular for each successive interspike interval (ISI) we classified the ISI as either high-frequency (greater than 40Hz or ISI < 25 ms, Fig. 6E, left) or low-frequency isolated spikes (less than 40Hz or ISI > 25 ms, with neighboring spike frequency <= 7.5 Hz, Fig. 6E, right). We assigned each ISI a joint MotR-SomR phase of the recorded LFP at the time of the second spike of the ISI. We then constructed a normalized rate modulation index (see Supplemental Methods) of where in MotR-SomR phase plane high-frequency ISI’s occurred (Fig. 6E, left) and separately where low-frequency ISI’s occurred (Fig. 6E, right). Green to red regions indicate where the respective ISI’s occurred at a higher rate than chance, for example a +100% modulation being a 2-fold increase in spikes relative to expected. Blue regions indicate where the ISI’s occurred at a lower rate than chance, for example a −50% modulation being a 2-fold decrease relative to expected. White regions indicate no statistically significant elevated or reduced ISI rates relative to chance. Chance levels were determined by computing ISI distributions in shuffled data (see Supplemental Methods).

Fig. 6E, left (Som:L5 high-frequency firing) and Fig. 6E, right (Som:L5 low-frequency firing) thus separately correlate high- and low-frequency single neuron spike pairs in SomR:L5 with joint LFP phases in SomR and MotR. It revealed several regularities. First, the red phase-trajectory line which produces the peak amplitude growth in SomR (Fig. 6D), passes through a substantial red region indicating an excess of high-frequency ISIs (maximum 187% above expected, a 2.87-fold increase), followed by a blue region indicating a significant deficit of high-frequency ISIs (minimum: −66% below expected, a 2.94-fold reduction), relative to the expected values (Fig. 6E, left). This pattern of successive ISI’s is characteristic of a burst. Thus, bursting in SomR:L5 neurons is strongly associated with MotRθ_2_-SomRθ_2_ phase-lags that lead to seizure amplitude growth in SomR. In contrast, the purple trajectory travels through regions with silencing of high-frequency firing (blue regions, minimum 68% below expected, a 3.13-fold reduction) and no substantial increase (small green region, maximum 41% above expected, a 1.41-fold increase Fig. 6E, left). Instead, the purple trajectory experiences a transient increase (111% above expected, a 2.11-fold increase) in low-frequency ISI’s followed by a decrease (63% below expected, a 2.7-fold reduction) (Fig. 6E, right). The combined silencing of high-frequency ISIs but increase of low-frequency ISI’s along the purple trajectory indicates an overall increase then decrease in isolated spikes, and absence of bursts. Thus MotRθ_2_-SomRθ_2_ phase-lags that lead to seizure amplitude decay in SomR are primarily associated with single spikes, not bursts. In summary, SomR:L5 bursts preferentially occur in the MotR-SomR LFP phase plane region that leads to SomR amplitude growth (red trajectory), while isolated single spikes preferentially occur in disjoint regions of the MotR-SomR phase plane that lead to SomR amplitude decay (purple trajectory).

Fig. 6F-G analyzes the relationship between MotR LFP amplitude growth or decay and MotR:L5 isolated spikes or bursts in a manner parallel to Fig. 6D-E. Indeed, we find a completely parallel result: MotR:L5 bursts preferentially occur in the MotR-SomR LFP phase plane region that leads to MotR amplitude growth, while isolated single spikes preferentially occur in disjoint regions of the MotR-SomR phase plane that lead to MotR amplitude decay. We can see this as follows. The purple reference trajectory which leads to MotR amplitude growth near 0 rad phase-lag (red region in Fig. 6F) traverses significantly increased (111% above expected, a 2.11-fold increase) then decreased (68% below expected, a 3.13-fold reduction) high-frequency ISI rates (purple trajectory in Fig. 6G, left) with a modest increase (57% above expected, a 1.57-fold increase) in low-frequency ISIs along this trajectory (green region and purple trajectory, Fig. 6G, right). This pattern is again indicative of bursting. In contrast, the red reference trajectory which leads to MotR amplitude decay near −π/2 rad phase-lag (red trajectory, blue region in Fig. 6F) experiences increased rate of high-frequency ISIs (107% above expected, a 2.07-fold increase) that is *not* followed by a significant decrease in the rate of high-frequency ISIs (red trajectory in Fig. 6G), but is rather accompanied by significantly increased rate (144% above expected, a 2.44-fold increase) of low-frequency ISIs (red trajectory Fig. 6G, right). This pattern is indicative of an overall increase in the rate of isolated spikes without an increase in bursts.

Thus, in both SomR and MotR we have found a strong correlation between bursts, MotR-SomR phase-lags, and SomR/MotR LFP amplitude growth. In particular, in both brain regions, bursts primarily occur at LFP phase-lags that lead to LFP amplitude growth for that region, while isolated spikes primarily occur at LFP phase-lags that lead to LFP amplitude decay. This tight correspondence leads to a striking prediction regarding bursts as follows. First, comparing Fig. 6D and Fig. 6F, we note that MotR amplitude growth and decay regions are disjoint from SomR amplitude growth and decay regions in the MotR-SomR phase plane. Thus, as discussed for amplitude growth in the model (Fig. 4K-M), either SomR amplitude grows and MotR amplitude moderately decays (red trajectory in Fig. 6DF) or MotR grows and SomR moderately decays (purple trajectory in Fig. 6DF). Now given the strong association between bursts and amplitude growth in both regions, our analysis predicts that bursts in the two regions should be anticorrelated in time; in essence when MotR bursts more, SomR should burst less and vice-versa.

We tested this prediction directly at level of bursts without reference to LFP. We examined cortical bursting in SomR:L5 and PreMotR:L5 at seizure onset. We found that SomR:L5 bursting increased 4.4-fold relative to the non-seizure state (0.22 ± 0.07 vs 0.97 ± 0.16 bursts/s p<0.0001, Welch’s t-test) and PreMotR:L5 increased 6.2-fold (0.12 ± 0.04 vs 0.74 ± 0.11 bursts/s, p<0.0001, Welch’s t-test). This increased bursting preceded seizure onset (Fig. 6H) and, as predicted by our model, burst rates in SomR:L5 and MotR:L5 were indeed anti-correlated in time across these two regions (Fig. 6I).

Finally, since PO thalamus is the most strongly frequency locked region at the level of single neuron spikes (Fig. 5D, middle and right), and since PO projects to both Mot:L5 and Som:L5, we further examined the relationship between spikes in PO and the MotR-SomR LFP. In Fig. 6J we plot the rate of PO ISIs relative to chance, as a joint function of SomR and MotR phase, in direct analogy to Fig. 6E for SomR:L5 spikes and Fig. 6G for PreMotR:L5 spikes. Intriguingly, we find that the rate of high-frequency ISIs undergoes a significant increase then decrease (bursting) along both the red and purple reference trajectories (Fig. 6J, left) as well as a similar increase then decrease in low-frequency ISIs (Fig. 6J, right). This indicates that no matter whether SomR amplitude is growing and SomR:L5 is bursting (red trajectory), or MotR amplitude is growing and PreMotR:L5 is bursting (purple trajectory), PO is bursting in both cases.

The strong connection between bursting in a brain region, and LFP amplitude growth, also held in transcallosal coupling from SomL to SomR. For example, the band 2 coupling function from SomL to SomR indicated SomR amplitude growth at 0 rad phase-lag and SomR amplitude decay at π rad phase-lag (Fig. S7B). Correspondingly, the phase trajectories that were associated with SomR LFP amplitude growth from SomL drive, were also associated with increased bursting in both SomR:L5 neurons and PO neurons (Fig. S7C) and phase trajectories associated with SomR LFP amplitude decay, were associated with increased spikes and not bursts in SomR:L5 (Fig. S7C, left and middle). We also examined several other brain areas, cell type, and firing modes associated with the phase tori (Fig. S7D-M). Notably, cortical inhibition followed anticorrelated patterns between Som/Mot regions and opposed to those observed for bursting above. In the thalamic reticular nucleus (TRN, the major inhibitory thalamic nucleus) high-frequency firing exhibited a strong association with the Som amplitude growth trajectory (Fig. S7I). These results suggest the bursting above is further modulated by or dependent on coordination of inhibition in the cortex and thalamus. Non-seizure related regions lateral posterior thalamic nucleus (a higher-order visual system analogue of PO) and hippocampal CA3/DG neurons showed no association with the phase torus (Fig. S7L).

Overall, in pairs of source and target regions (e.g. Som-Mot, Mot-Som, or SomL-SomR) where our model predicts target region LFP amplitude growth as a joint function of source and target band 2 phase-lags, we find the target region exhibits more bursting. Conversely, when our model predicts LFP amplitude decay, the target region exhibits more isolated spikes and less bursts. Finally, when any of these target regions exhibit amplitude growth and bursting, PO also bursts.

### Posterior thalamic nucleus activation enhances premotor to somatosensory functional connectivity

Our model suggested that high between-region connectivity drives seizures (Fig. 4CD). Previous work has shown that PO activity can increase the effectiveness of inputs to Som:L5 and PreMot:L5 pyramidal neuron distal apical dendrites.^31^ These dendrites are precisely the site of PreMot:L5 to Som:L5 and Som:L5 to PreMot:L5 and transcallosal Som:L5 to Som:L5 and Mot:L5 to Mot:L5 feedback connectivity.^29, 31^ Moreover, Som:L5 and PreMot:L5 bursting followed their respective connection specific amplitude growth trajectories (Fig.6D-G), while PO bursting increased along both growth trajectories (Fig. 6J) as well as the SomL-SomR growth trajectory (Fig. S7B-C). This suggests PO’s role may be to increase overall cortico-cortical functional connectivity. We thus decided to directly test whether PO activity enhances functional connectivity between these areas in the context of absence seizures. In *Scn8a*^*+/−*^*/Thy1-ChR2* mice, we implanted an optical fiber in PreMot:L5, injected AAV–ChrimsonR in PO, and placed a second optical fiber above PO (Schematic in Fig. 7A). This setup allowed us to optogenetically stimulate PreMot:L5 pyramidal cells and PO thalamocortical cells independently with two different light wavelengths. Injections were largely confined to PO (Fig. 7B) with tdT labeled projections limited to Som:L1 and 5 (Fig. 7C and D). Som:L5 firing was recorded with Neuropixels during optogenetic stimulation of either PreMot:L5, PO, or both. When stimulating both, we stimulated PO and then PreMot:L5 with a delay of either 0 ms, 15 ms, or 60 ms corresponding to band 2 phase-lags of 0, −π/2, and 2π rad respectively. The choice of delay of 15 ms was motivated by the early increase in PO firing (Fig. 6B arrow 1) which occurred −14 ms before the peak in Som:L5 firing and was simultaneous with the PreMot:L5 shoulder (Fig. 6B arrow 2). These variable delays allowed us to determine how the temporal lag between PO and PreMot input to Som:L5 determines the effectiveness of PreMot input to drive Som:L5 spiking.

**Figure 7.**
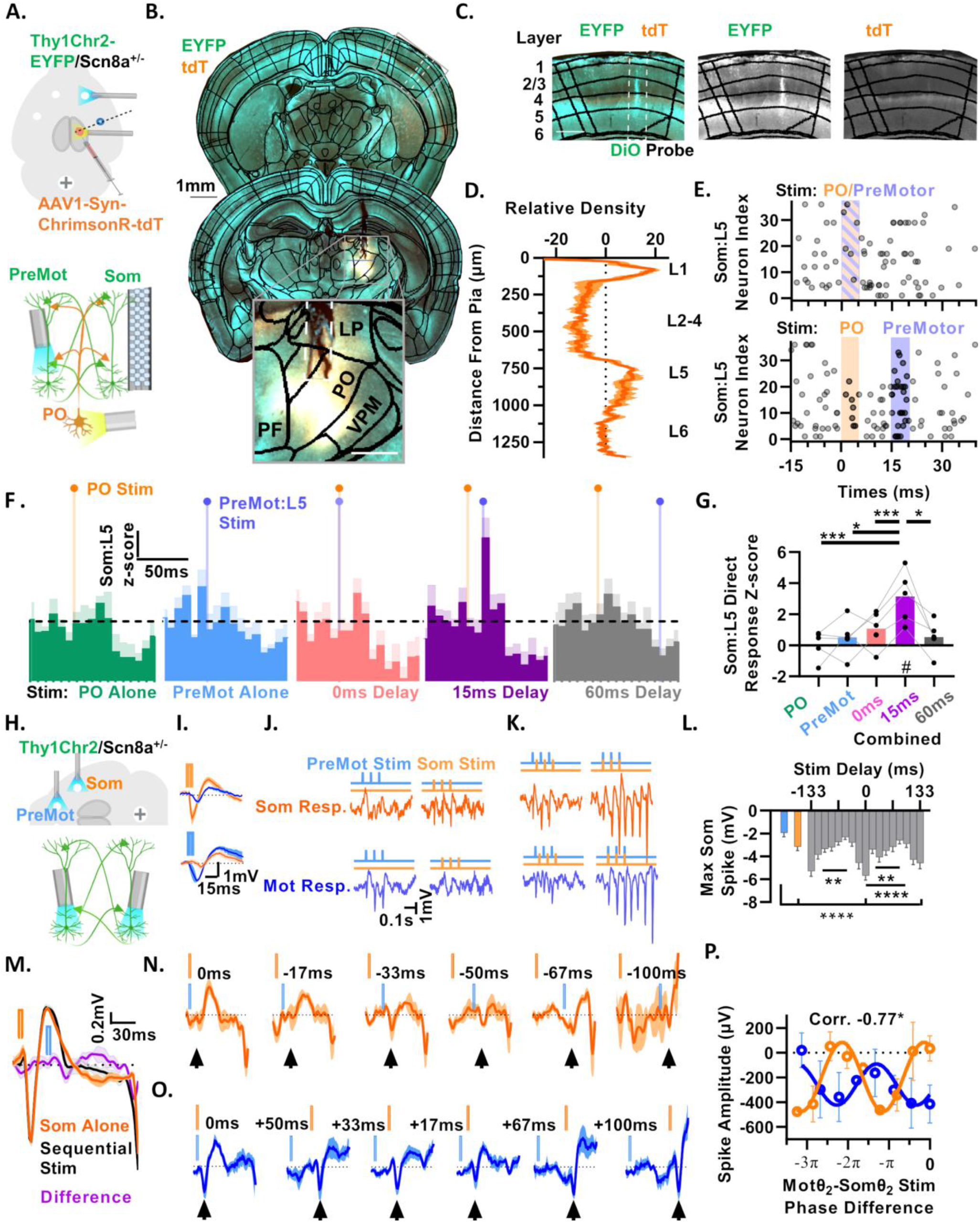
PO thalamus gates the PreMot:L5–Som:L5 pathway to promote absence seizures. (A) Experimental schematic: dual-fiber implants in *Scn8a+/−/Thy1-ChR2* mice for optogenetic stimulation of PreMot:L5 and PO while recording single-unit firing in Som:L5 Face/Barrel cortex. (B) Example histology of chrimsonR-tdT injection in PO and labeled fibers in L1 and L5 of Som cortex. Boundaries defined by SharpTrack. (C) (Left) Expanded view of recording site from (B) with DiO labeled probe (green vertical tract in white box), Thy1-GFP (green) expression in L1/L5, and tdT+ PO fibers (orange). (Middle) Green channel; (right) red channel. (D) Quantification of PO fiber density relative to mean along cortex depth at recording sites. (E) Som:L5 neuron spike raster during combined PO–PreMot stimulation at 0ms (top) or 15 ms (bottom) delay. Raster overlay from 10 trials. (F) Average of z-scored peristimulus time histograms (PSTHs) for PO, PreMot, or combined stimulation at 0, 15, or 60 ms delays for 5 recordings from 4 mice, error bars indicated SEM. Orange (PO) and Blue (PreMot) circles indicate time of stimulation. (G) Quantification of direct stimulus responses at 15-20 ms bin following initial stimulation (# indicate one-way t-test vs 0, asterisk indicate between group comparisons via paired t-test). (H) Experimental schematic: dual fiber implants in *Scn8a*^*+/−*^*/Thy1-ChR2* mice for optogenetic stimulation of PreMot:L5 and Som:L5 cortex. (I) Average with SEM effect of Som:L5 (top) and Mot:L5 (bottom) stimulation in Mot (blue) and Som (Orange) Ecog from 4 mice. (J an K) Stimulation patterns and resulting Ecog LFPs for PreMot and Som stimulation: single site (J) and combined antiphase (K left) or aligned (K right) stimulation. (L) Average minimum spike amplitude across 10 trials under each time-lag condition. Lines show pairwise t-tests. (M) Effect of Som:L5 stimulation alone (orange trace), the sequential stimulation of Som:L5 and then Mot:L5 33 ms later (black trace), and the difference between the two above traces to isolate the effect of Mot:L5 stimulation at a particular time-lag after Som:L5 stim (purple). (N) Average traces of the effect of Mot:L5 stimulation in Som Ecog at different time delays after Som:L5 stimulation after subtraction of Som:L5 stimulation alone as in M. Times indicate Mot-Som stimulation time-lag. Arrows indicate timing of spike. (O) As in N for the effect of Som:L5 stimulation in Mot Ecog. Times indicated Mot-Som stimulation time-lag. Order aligned to equivalent Mot-Som band 2 phase-lag of Som in O. (P) Circles indicate average spike amplitudes in N and O. Solid lines indicate sinewave fit. Asterisk indicates significant anticorrelated spike amplitudes at the given band 2 phase-lags.

We found that stimulating PO 15 ms (but not 0 ms or 60 ms) before stimulating PreMot:L5 induced robust Som:L5 firing during the PreMot:L5 stimulation window (Fig. 7EF). Notably, stimulating PO alone or PreMot:L5 alone induced no subsequent strong direct spiking increase in Som:L5 response (Fig. 7F, left 2 plots). Over longer time-scales after stimulation, a subset of stimulation cases exhibited Som:L5 reduced firing followed by a subsequent increased rebound, triggering an oscillation (Fig. S8B-D). Notably, stimulating PO then PreMot:L5 at 0 ms or 15 ms delay triggered a transient seizure like oscillation at the fundamental frequency of endogenous seizures (Fig. S8E). In contrast, stimulating PO alone or PreMot:L5 alone did not, whereas stimulating PO then PreMot:L5 at 60 ms delay triggered a transient oscillation at a much lower frequency than that of endogenous seizures (Fig. S8E).

In summary, only stimulation of PO then PreMot:L5 at a 15 ms time-lag induced *both* a direct response in Som:L5 *and* a transient seizure like oscillation. This causal effect is consistent with endogenous seizure dynamics wherein PO firing becomes substantial −15 ms before (Fig. 6B arrow 1) a shoulder of significant PreMot:L5 firing (Fig. 6B arrow 2), which is coincident with peak Som:L5 firing.

### Multisite optogenetic stimulation of Som:L5 and PreMotor:L5 cortex cooperated in phase lag-dependent seizure generation

The model predicted LFP seizure amplitudes grow due to phase-lag dependent coupling between regions in an anticorrelated manner. For example, at particular values of the band 2 Motθ_2_-Somθ_2_ phase lag, Som amplitude grows due to Mot input (red region and red trajectory, Fig.4H, right) but Mot amplitude does not grow due to Som input (blue region Fig. 4H, left). However, at opposite values of the band 2 Motθ_2_-Somθ_2_ phase-lag Mot amplitude grows due to Som input (red region and purple trajectory, Fig. 4H, left) but Som amplitude does not grow due to Mot input (light blue region, Fig. 4H, right). Moreover, the timing of the band 2 trough is strongly correlated with that of the SWD-spike (Fig. 2CD), and between-region time-lags of the band 2 trough reflect those of the SWD-spike (Fig. 2K, right and Fig. 2N). Therefore, we decided to test whether SWD-spike amplitude responses in Mot and Som were sensitive to, and indeed anticorrelated with respect to, temporal lags in the stimulation of PreMot:L5 and Som:L5, as suggested by our model. To do so, we implanted *Scn8a*^*+/−*^*/Thy1-ChR2* mice with optical fibers in PreMot:L5 and Som:L5, allowing us to independently stimulate PreMot:L5 and Som:L5 pyramidal cells while recording Ecog signals in both regions in layer 1 (Fig. 7H). We delivered 3 pulses to each region at the fundamental seizure frequency of 7.5 Hz, at a low enough intensity that a single pulse would not strongly propagate to the other regions. Pulses were delivered to PreMot alone, Som alone, or to both with delays between the first pulses spanning −133 ms to +133 ms.

On average, PreMot:L5 stimulation in isolation produced small local Mot SWD like Ecog responses (−1.1 ± 0.03 mV) peaking at 9.5 ms post stimulation onset, with a much smaller (−0.23 ± 0.16 mV) nonlocal Som Ecog response that peaked at 15.9 ms post stimulation onset (Fig. 7I, top). Som:L5 stimulation in isolation showed the reverse response pattern (Som peak = −1.5 ± 0.33 mV at 5.7 ms, Mot peak = −0.42 ± 0.13 mV at 14.3 ms, Fig. 7I, bottom). Fig. 7J shows single trial examples of the trial average in Fig. 7I, again showing that stimulation of one region alone at low intensity does not induce seizure like oscillations. However, large SWDs were only obtained when stimulating both PreMot:L5 and Som:L5, which depended strongly on the relative timing of stimulations. Recall the fundamental 7.5 Hz seizure period is 133 ms. Stimulating using half this relevant time-lag (−67 or +67 ms did not yield large SWD-spikes (Fig. 7K, left), but stimulating at 0 time-lag did (Fig. 7K, right). Indeed, the response to stimulation depends strongly on the stimulation time-lag from −133 ms to +133 ms as measured by the largest SWD-spike amplitude during stimulation (Fig. 7L and Fig. S8F) and by the seizure score in the post stimulation period (Fig. S8G). Remarkably, stimulation near −17 to 0 ms time-lags (the time-lags corresponding to phase-lags predicted to lead to amplitude growth for Som and Mot band 2 respectively) induced self-sustaining seizures, but stimulation near ± 67 ms did not (Fig. 7K and Fig. S8G).

To test our prediction of anticorrelated regional responses as a function of the time-lag in stimulation, we determined the specific contribution to the response in one region due to input from the other region at different time-lags relative to the first stimulation. To do this, we subtracted the response to stimulation in one region from the response to stimulation of both regions (e.g. response in Som due to both Som:L5 and PreMot:L5 stimulation minus response in Som due to Som:L5 stimulation alone (Fig. 7M), and similarly for responses in Mot). This differential response reflected specific contributions to the response in one region from the other region at defined time-lags and is plotted for several different stimulation time-lags (PreMot:L5 stimulation to Som contribution, Fig. 7N, and Som:L5 stimulation to Mot contribution, Fig. 7O). In both cases the between-region contribution, as measured by the SWD response peak after stimulation from the opposite region (Fig. 7NO, black arrows) displayed a strong dependence on stimulation time-lag. The variation in spike amplitude varied with a timing dependence resembling the frequency of band 2. Fitting a sinewave to these curves revealed a periodic effect with a frequency near band 2 (Mot: 13.4 ± 1.3 Hz, R^2^=0.74 and Som: 12.2 ± 0.6 Hz, R^2^=0.92) (Fig. 7P). The phase offset was ~opposite between regions (Mot phase=-0.50 ± 0.52 rad vs 2.04 ± 0.29 rad p=0.0078).

We thus plotted the evoked SWD-spike like amplitude response against the stimulation time-lag in units of the band 2 Mot-Som phase difference (Fig. 7P). This showed the maximal response in Som and Mot due to specific contributions from the other region were indeed anticorrelated: at band 2 phase-lags where Mot responds strongly due to Som input, Som responds weakly due to Mot input, and vice versa (Corr. = −0.77). This is reminiscent of the disjoint regions of amplitude growth in Mot and Som as a function of band 2 phase-lag between Mot and Som (Fig. 4HK).

Overall, these results directly demonstrate a multi-area seizure generation mechanism, in this case emerging from interactions between PreMot:L5 and Som:L5. Disjoint between-region phase-lags lead to either Mot SWD-spike growth or Som SWD-spike growth, specifically due to input from the other region at specific time-lags (Fig 7P), in direct analogy to how our model predicts that disjoint phase-lags lead to band 2 amplitude growth in Mot or Som (Fig. 4HK).

## Discussion

### Interpretable machine learning reveals an emergent mechanism for a low dimensional chaotic attractor state underlying LFP dynamics in absence seizures

Our brain-wide Ecog recordings (Fig. 1A) revealed a multiregional somatosensory and motor seizure network whose LFP amplitude rapidly and simultaneously elevates at seizure onset (Fig. 1C). While individual seizure trajectories rapidly and chaotically diverged in time from each other (Fig. 1DE), robust correlations between regions within a seizure were maintained (Fig. 1GH). To describe the underlying oscillatory seizure dynamics, we examined mean phase-lags, coherences, and amplitude trajectories between three harmonic bands across 4 representative regions (Fig. 2, Fig. S2HIL), resulting in 60 metric properties of the seizure that were impossible to understand in isolation. However, we used interpretable machine-learning to identify a dynamical model that simultaneously reproduced all 60 metric properties to within a precision matching inter-subject variability (Fig. 3, Fig. S3A-E, Fig. S4B). Moreover, our model revealed that the seizure state corresponds to a low 5.48-dimensional (Fig. 3CD) chaotic (Fig. 3H, Fig. S3FG) attractor state (Fig. 3EGH).

The model also revealed a simple between-regional dynamical mechanism for the origins of LFP synchronization, amplitude growth, and chaos. Our data-driven learned model remarkably converged to a coupled Kuramoto oscillator type interaction^27 35-37^ between brain regions with additional amplitude dynamics (Equation 1 and 2, Fig. 4BHIJK, Fig. S5D-J), that can lead to more complex phenomena.^38^ Both in the Kuramoto model and in our amplitude enhanced model, synchronization between regions occurs when between-regional coupling strengths overwhelm between-region heterogeneity in intrinsic oscillation frequencies, leading to multiple regions oscillating at a common locked frequency near seizure onset (Fig. 4DE), a model prediction which we successfully confirmed in the experimental data (Fig. 4FG). We previously showed reduced heterogeneity in voltage-gated sodium channels promotes synchronization and epilepsy by reducing intrinsic firing frequency heterogeneity, consistent with decreased biophysical heterogeneity in human epilepsy.^39–41^ However, here increased coupling strength better explained frequency locking in experimental seizures than did reduced intrinsic frequency spread (Fig. 4D-F, Fig. S4KL).

Near the onset of synchrony, regions start to oscillate at various relative phase-lags. Analysis of the between-region couplings of the model show that these couplings drive LFP amplitude growth in a phase-lag dependent manner (Equation 1). For example, if a pair of regions 1 and 2 oscillate at the coupling from region 2 to region 1’s preferred phase-lag, then the coupling will cause rapid amplitude growth in region 1. However, if they oscillate far from the preferred phase-lag, there will be reduced or no amplitude growth in region 1. Often times, the reverse coupling from region 1 to region 2 has a different preferred phase-lag. The combined amplitude and phase dynamics of the system would then cause the phase-lag between region 1 and region 2 to vary over time between these two phase-lags, leading to alternating periods of amplitude growth or no growth (or even moderate decay) in region 1 and region 2. This type of bidirectional coupling between-region pairs features prominently, for example in MotL-SomL (Fig. 4HK), and MotR-SomR (Fig. 6DF). This leads to a temporally alternating scenario where Mot amplitude grows due to Som input, but Som amplitude does not grow in response to Mot input, and then the opposite where Som amplitude grows due to Mot input, but Mot amplitude does not grow in response to Som input. Moreover, the alternation between these two scenarios is chaotic, because the combined feedback dynamics of amplitudes and phases amongst multiple regions itself is chaotic (Fig. 4M-O, S5H), as quantified by positive Lyapunov exponents (Fig. 3H). Importantly, the interpretable nature of our model allowed us to identify the cell types and firing properties involved: PO potentiation of between-region L5 cortical connectivity that drives phase-lag dependent burst firing.

Our model reveals an emergent mechanism for seizure genesis that arises from phase-lag dependent amplitude growth through many between-regional couplings resulting in a complex chaotic attractor state, rather than seizures originating in and then spreading from a single brain region. Discovering and understanding these between-region dynamic drives for seizure generation would not be feasible without the use of data-driven interpretable machine learning. The chaotic dynamics and non-reducibility of seizure drive to a single region, cell type, or constant firing pattern indicates fundamental constraints on how we can understand, predict, or control seizures. For example, because an emergent mechanism arises from many interacting processes, none of which causes the seizure on its own, reduction of seizure causes to single brain regions, cell types, or proteins in isolation will inevitably produce confounding results. Importantly, our results are at odds with a focal mechanism, cortical or otherwise, and provides an explicit mechanism for an emergent origin of absence seizures as indicated previously.^9, 42^

### The dynamic LFP model predicts the spatiotemporal organization of single neuron spiking

We next moved beyond LFP to a finer scale of analysis at the level of single neuron spiking across cortex, basal ganglia, hippocampus, and thalamus (Fig. 5AB). Remarkably, the dynamic behavior of our LFP model allowed us to reveal striking regularities in the spatiotemporal organization of individual spikes in a manner not possible without the model. First, we found individual neuron spiking moved from a heterogeneous distribution of frequencies in the non-seizure state to a locked common frequency in the seizure state (Fig. 5CD). This analysis was directly motivated by the analogous behavior of LFP, first predicted in the model (Fig. 4DE) and then confirmed in experiment (Fig. 4FG). Recent studies have reported heterogenous changes in the mean single neuron firing rates at absence seizure onset.^3, 4^ Our work explains such heterogenous changes as they are exactly what is required to achieve a common locked spike frequency across neurons in the seizure state: neurons that fire at higher (or lower) spike rates in the non-seizure state must lower (or raise) their firing rates to fire at a common rate in the seizure state. Importantly, analysis of the time dependence of this frequency locking (Fig. 5D) revealed an early seizure network consisting of single neurons in PO, Som:L5, and PreMot:L5, that were the earliest and most strongly frequency locked regions (Fig. 5C and Fig. 5D, middle, right). Consistent with this observation, these three populations have strong anatomical interconnectivity.^28–30^ We therefore focused our subsequent spiking analyses on these three areas, first focusing on the relationship between spiking and LFP.

Single neuron spikes occurred at a wide range of temporal lags with respect to LFP variables like the SWD-spike (Fig. 6AB) and SomR or MotR band 2 phases (Fig. 6AC). On the surface, these observations in isolation did not clarify a correspondence between spike patterning and LFP. However, guided by our model, we found a tight correspondence between band 2 between-region phase-lags, alternating amplitude growth in Som and Mot, and alternating tonic versus burst firing in single neuron spiking. Indeed, a prominent feature of spiking in the seizure state is the prevalence of bursts in addition to tonic single spike firing (Fig. 5C and Fig. S6FG). We analyzed when tonic spiking versus bursting occurred in both Som:L5 and PreMot:L5 as a function of the band 2 phases Somθ_2_ and Motθ_2_ and we found a striking correspondence: when the model predicts LFP amplitude growth in a given region (Fig. 6DF, Fig. S7A), that region exhibits more bursting (Fig. 6EG, left, Fig. S7C left); conversely when the model predicts amplitude decay (Fig. 6DF, Fig. S7B), that region exhibits more tonic firing (Fig. 6EG, right, Fig. S7C, middle). Given that amplitude growth in Mot and Som is anticorrelated over time due to chaotic variation in the Motθ_2_-Somθ_2_ phase-lag (either Mot amplitude grows and Som does not, or vice versa), our analysis then predicts that bursting in Som:L5 and PreMot:L5 should be anticorrelated in time (either Som:L5 bursts and PreMot:L5 does not, or vice versa). We directly confirmed this striking prediction in the data (Fig. 6HI).

This analysis reveals a privileged role for band 2 in relating LFP to SWD-spikes. Indeed, at the level of LFP alone, the trough of band 2 is strongly correlated with the timing of the SWD-spike (Fig. 2CD, Fig. S2A) and between-region band 2 phase-lags captured the corresponding between-region SWD-spike time-lags (Fig. 2K-R, Fig. S2JK). Now connecting LFP to single neurons, the SWD-spike occurred with an approximately 4 ms delay relative to peak single neuron spiking (Fig. 6B). Also given that band 2 phase-lags determine both amplitude growth (Fig. 6DF) and burst versus tonic firing (Fig. 6EG), a natural hypothesis is that band 2 LFP at 15 Hz predominantly reflects the time course of the calcium plateau and repolarization associated with cortical burst firing, while the slower 7.5 Hz interval of single neuron spiking (Fig. 5CD) reflects an inability of neurons to fire again until the second cycle of band 2, mediated for example by inactivation of sodium and calcium channels.^32^ These results are consistent with previous observations that absence seizures appear to arise from the progressive amplification of every second trough of a spindle-like oscillation with a frequency of twice the SWD-spike interval frequency.^12^ Fittingly, the first line anti-absence seizure medication ethosuximide blocks T-type calcium channels which promote cortical and thalamic burst firing, which by our analysis above is associated with LFP amplitude growth.^4, 43^ This suggests disruption of the underlying drive towards the absence seizure attractor is the therapeutic target of the leading anti-absence treatment.

### Higher order thalamus amplifies cortico-cortical functional connection strength during absence seizures

In our LFP model, increased coupling strength between-regions drives seizures (Fig. 4C, Fig. S5G). Also, our analysis of the higher-order thalamic nucleus PO, part of the early seizure network with Som:L5 and PreMot:L5 (Fig. 5D) revealed that on each seizure cycle, PO spiking increases before Som:L5 and PreMot:L5 (Fig. 6B) and PO bursts with both of them as a function of the band 2 Motθ_2_-Somθ_2_ phase-lag (Fig. 6EGJ). PO has been implicated in absence epilepsy previously with some of the earliest pre-seizure activity changes.^44^ Moreover, previous work has found that PO projects to both cortical layers 1 and 5 of Som and PreMot, and potentiates feedback coupling between layer 5 pyramidal neurons in these two regions by depolarizing their apical dendrites, lowering the threshold for layer 1 inputs to trigger somatic spikes.^31^ Somatic inputs, when paired with a strong coincident or slightly prior distal input, elicit amplified dendritic depolarization and voltage gated sodium and calcium channel dependent burst firing.^45–47^ This distal input could in principle be provided by PO inputs early in the SWD-spike oscillation cycle (Fig. 6B) which would then prime PreMot:L5 and Som:L5 apical dendrites to promote burst firing when long range input from other regions arrive, either from the PreMot:L5/Som:L5 connections or from transcallosal inputs to these regions.

We tested this hypothesis experimentally with multisite optogenetics, observing that stimulation of PO 15 ms before stimulating PreMot:L5 causes a large increase in Som:L5 firing when PreMot:L5 is stimulated (Fig. 7E-G) and triggers oscillations at the fundamental seizure frequency (Fig. S8E). In contrast, stimulation of PreMot:L5 or PO in isolation does not. Thus, we conclude the strong coupling between regions in the model that drives seizures is mediated by PO-induced amplification of cortico-cortical L5 functional connectivity across the motor and somatosensory systems. Additionally, these same mice show post-seizure onset myelination increases resulting from myelin plasticity, suggesting a contribution of a structural connectivity increase.^48^

Notably, neurons in PreMot and Som not only receive projections from PO but also send projections back to PO. This suggests future investigation of the effect of top-down cortical control of PO could further refine this mechanism. This top-down control could be mediated through direct cortico-PO projections, or indirectly via projections to the basal ganglia or thalamic inhibitory nucleus RT.^49^ Correspondingly, burst frequency corticothalamic fiber stimulation drives RT dependent post-inhibitory rebound bursting and oscillation frequency modulation of thalamocortical neurons.^7^ Our prior work showed *Scn8a* knockdown in RT can cause absence seizures and disrupts RT-RT inhibition.^24^ Furthermore, RT-RT inhibition powerfully suppresses thalamic synchrony.^42^ This suggests absence seizure oscillations emerge as PO thalamus amplifies between-regional feedback connectivity, thereby generating alternating somatosensory and motor system burst firing within a chaotic attractor, in a manner that RT is unable to shut off when *Scn8a* is knocked down. Future work should examine further cell-type specificity and explore additional regulators of this process such as dendritic inhibition and intra-thalamic circuits.^11, 50^

### Multisite optogenetic stimulation reveals a direct model prediction of phase-lag-dependent seizure growth

At the level of LFP amplitudes, our model revealed a simple mechanism for seizure growth: at certain phase-lags between SomL and MotL, input from SomL to MotL causes amplitude growth in MotL, while input from MotL to SomL does not cause amplitude growth in SomL. At other phase-lags, the opposite happens (no amplitude growth in MotL but amplitude growth in SomL) (Fig. 4HJK). We tested this prediction at the level of driving source spiking with optogenetic stimulation of Som:L5 and PreMot:L5 while recording Ecog in both regions (Fig. 7H). Remarkably, as predicted by our model, we found a strong dependence of the measured LFP amplitude in both regions on the time-lag between stimulation of each region (Fig. 7KL and Fig. S8H). Furthermore, when isolating the specific contribution of input from one region to the amplitude of the first SWD-spike in the other region (Fig. 7M), we found a strong anticorrelated response as a function of the stimulation phase-lag (Fig. 7N), as predicted by our model. At certain stimulation phase-lags between Som:L5 and PreMot:L5, input from Som:L5 to PreMot:L5 leads to a maximal response in Mot, while input from PreMot:L5 to Som:L5 leads to a minimal response in Som (Fig. 7P). At other phase-lags, the opposite happens (minimal response in Mot and maximal response in Som).

In summary, our model-guided experiments revealed three related anticorrelated properties of Mot and Som as a function of (stimulation) phase-lags between them: model LFP amplitude growth versus no growth (Fig. 4HK), bursting versus tonic firing (Fig. 6I), and strong versus weak functional connectivity (Fig. 7P). These three properties covary together in one region and are opposite in the other regions. For example, when one region will have LFP amplitude growth, high burst rates, and strong functionally enhanced input connectivity, the other will have the opposite in all 3.

### A common circuit mechanism with general anesthesia

A defining clinical feature of absence seizures is a sudden loss of consciousness that lasts for a typical duration of tens of seconds, followed by an equally sudden restoration of consciousness. Remarkably, there are no overt effects on cognition or awareness just before or just after the seizure. This rapid, specific, and direct transition from consciousness to unconsciousness and back at the beginning and end of the seizure, suggests that the absence seizure network may be a central substrate of consciousness.

Our results indicate that during seizures, this network undergoes disrupted long-range cortico-cortical and cortico-thalamic communication. Specifically, we identify the posterior thalamic nucleus (PO) as modulating feedback connectivity strength between motor and somatosensory cortices. Notably, corticothalamic loops of the kind we identified, particularly those involving higher-order processing rather than first-order sensory processing, have been proposed as substrates of consciousness.^51–54^ This proposal aligns with recent findings that PO regulates cortico-cortical communication by depolarizing apical dendrites of cortical L5 pyramidal neurons and that this process is disrupted under anesthesia.^31^ Such disruption would lead to attenuated signal flow between Mot and Som systems. The overlap between absence seizure networks and anesthesia sensitive circuits suggests a shared mechanism of loss of consciousness: disrupted between-region L5 pyramidal coupling linked to PO dysregulation of feedback connectivity.

Our results may also connect to one framework for thinking about consciousness: the Global Neuronal Workspace (GNW) Theory.^55^ This framework emphasizes the integration of local information processing within a shared larger-scale network through a global broadcasting principle that could include wide-spread communication via distributed networks like motor and somatosensory areas. Our results show that seizures reflect PO-induced hyper-functional connectivity between motor and somatosensory areas, which in turn drives brain dynamics to a pathological low-dimensional chaotic attractor state that would impede global broadcast signals necessary for consciousness in the GNW theory. Similarly, in anesthesia, attenuated cortico-cortical functional connectivity could also impede such global broadcast signaling. Overall, highly regulated cortico-cortical functional connectivity that is neither too large, as in seizures, nor too small, as in anesthesia, may be essential for both global broadcasting and consciousness, as well as possible effective transitioning between brain states underlying consciousness.^56^

### Limitations

Our model, like all models, is an abstraction of more complex dynamics and advances understanding at a specific level of abstraction. We separated signals into discrete bands and the identified model used nonlinear cross-frequency coupling to handle interactions between frequency bands, a result similar to those from nonlinear optics.^57^ Other embeddings like PCA, UMAP, or autoencoders also decompose signals but without constraints on interpretability and are less readily compatible with symbolic equation discovery.^58^ But still, obtaining biological interpretations of individual bands should also be done cautiously. For this reason, we often returned to the SWD-spike as a focal point of analysis. Our model, trained on LFPs, reflects a combination of spiking, synaptic activity, and subthreshold voltage changes.^59^ Also, our model focused strictly on identifying the causes of absence seizure generation. Importantly, it does not model the termination of seizures. We propose the model coefficients represent a snapshot of the more slowly varying parameters that support seizure generation. These slow variations include arousal, attention, sensory input, and neuromodulation, all of which have been shown to impact seizure prevalence.^6, 60–62^ However, each of these processes are highly complex and beyond the scope of this study. Thus, our model is a simplification of underlying influences affecting neuronal dynamics. Nevertheless, by focusing on learning the model from data specifically around the time of seizure onset, we obtained a highly accurate model that captured 60 seizure metrics 3 s into the future seizure from a non-seizure initial state, and yielded many novel experimentally confirmed predictions about the dynamical causes of absence seizures.

## Supplemental Figures

**Figure S1.**
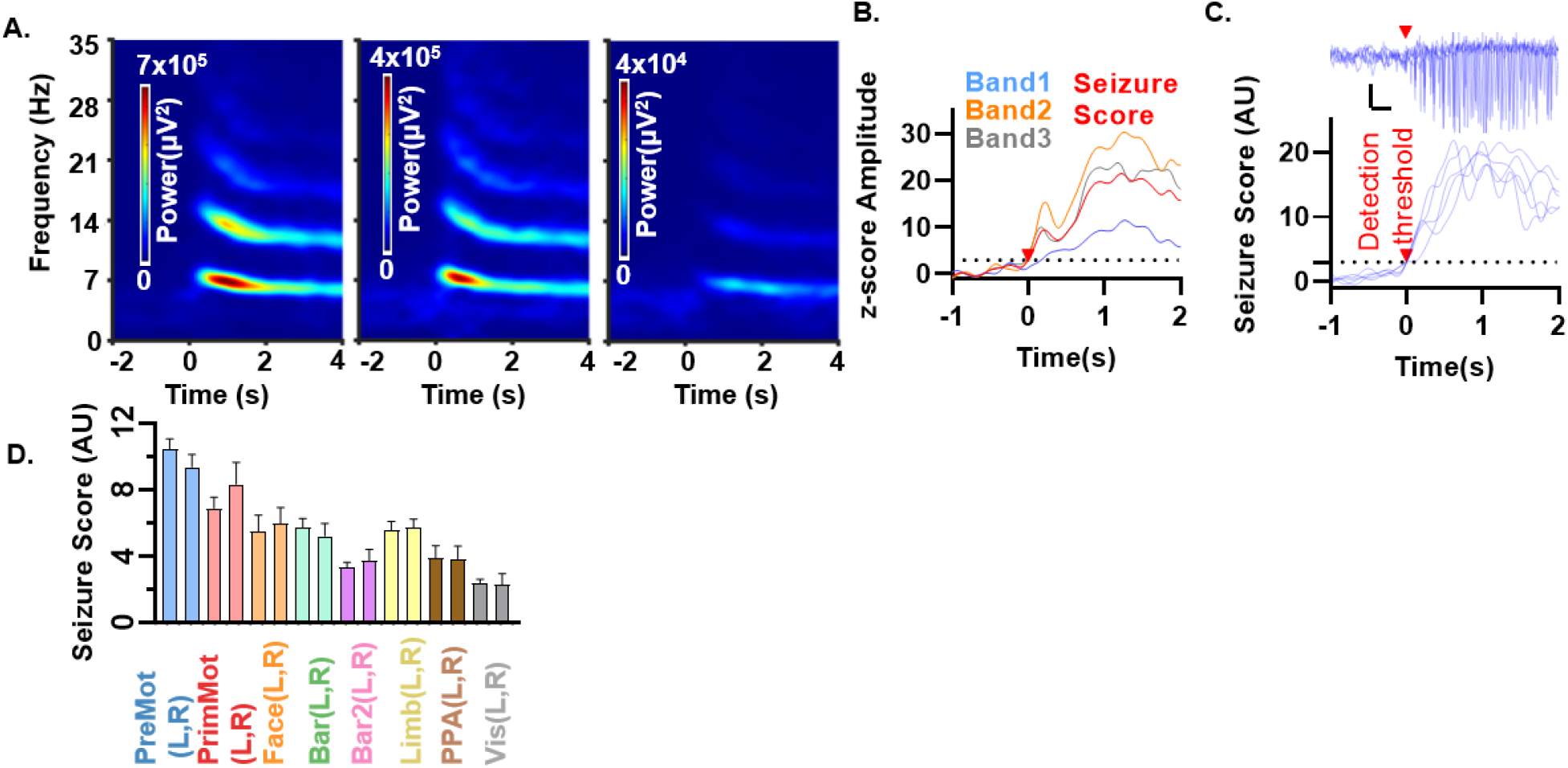
Seizure amplitude dynamics quantified absence seizure power. (A) Time-frequency spectrograms of Ecog signals for different brain regions in Premotor Left (PreMotL), Face Left (FaceL), and Visual Left (VisL) showing increases in power in bands that are harmonics of the fundamental frequency. (B) z-score amplitude for an example PreMot band 1, 2, and 3 and the seizure score (the average of the z-score of the amplitude envelope of each of the 3 bands). Note band 1 exhibits the smallest change in amplitude. (C) Five PreMotL seizures (top) aligned to crossing seizure score of 3 (red arrow), the detection threshold of seizures. Scale bar 200 ms and 2 mV. Onset times defined as last time point of crossing seizure score of 1 prior to passing the detection threshold. (D) Average and SEM seizure score for 3 s as in C after onset at the leading site across all recorded regions.

**Figure S2.**
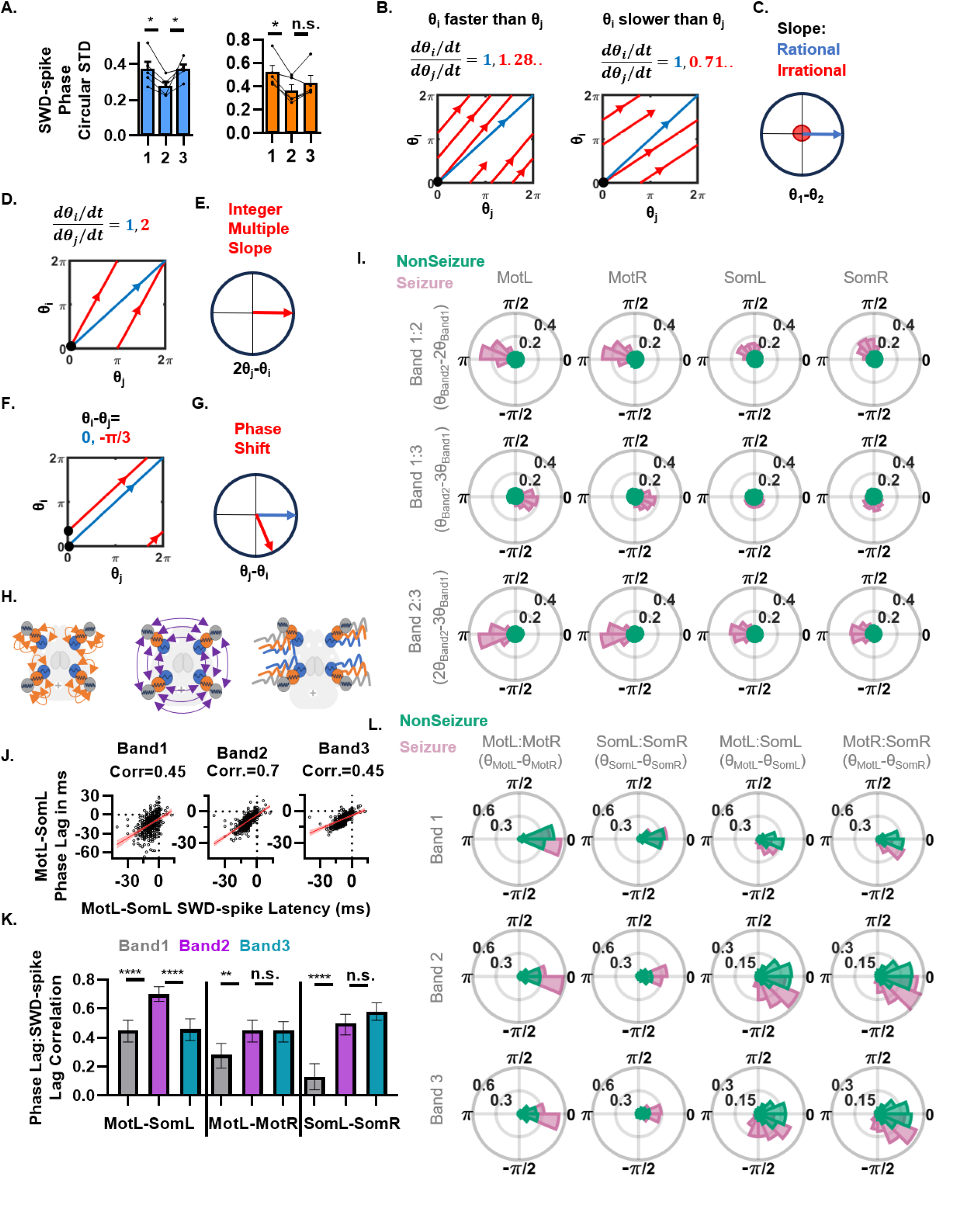
(A) Circular standard deviation of the phase of the SWD-spike showing SWD-spikes are more precisely locked to band 2 than band 1 in Mot (blue) and Som (orange). Paired t-test. (B-F) Schematics showing several theoretical phase relationships on toroidal manifolds. (B) Effects of different phase velocity ratios on theoretical trajectories. 1:1 ratio phase velocities (blue) return to same trajectory while irrational frequencies with θ_i_ > θ_j_ (red trajectory B, left) and θ_i_ < θ_j_ (red trajectory B, right) drift filling the torus. (C) Phase difference histogram quantifies mean phase-lag (m) and phase coherence (r) of the corresponding relationships as in B with no mean phase-lag and no phase coherence when slopes are irrational (red) and strong coherence (r) and phase-lag (m) when differences are locked at a ratio of 1 with zero phase-lag (blue). (D) Rational slopes on the torus (here 2, red) also return to the same trajectory similar to slope 1 trajectories (blue). (E) Phase difference histogram quantifies phase-lag and coherence when the slower phase is multiplied by the ratio of the two velocities. (F) Oscillation phase-lags on the torus of integer slope trajectories alter the intercept (−π/3 red vs 0 rad blue) but not slope of the trajectory. (G) Phase histogram quantifies change in phase-lag as a shift in the angle on the histogram. (H) Schematic of the between-band (left) and between-region (middle) relationships and amplitude trajectories (right) used in defining the state space of absence seizures. (I) Phase-difference histograms across all between-band relationships in (H). Each of the 12 histograms has a seizure specific coherence and phase-lag that varies substantially across bands and regions. (J) Scatter plots of individual SWD-spike time-lags between MotL and SomL and the observed phase-lag converted to ms as in main text. Note high correlation with between band 2 phase-lags and SWD-spike lags. (K) Correlations as in J across different regions. Note band 2 is higher than band 1 across all connections and higher than band 3 in the MotL-SomL. (L) Phase-difference histograms across all between-regional relationships in (H). Each of the 12 histograms has a seizure specific coherence and phase-lag that varies substantially across bands and regions.

**Figure S3.**
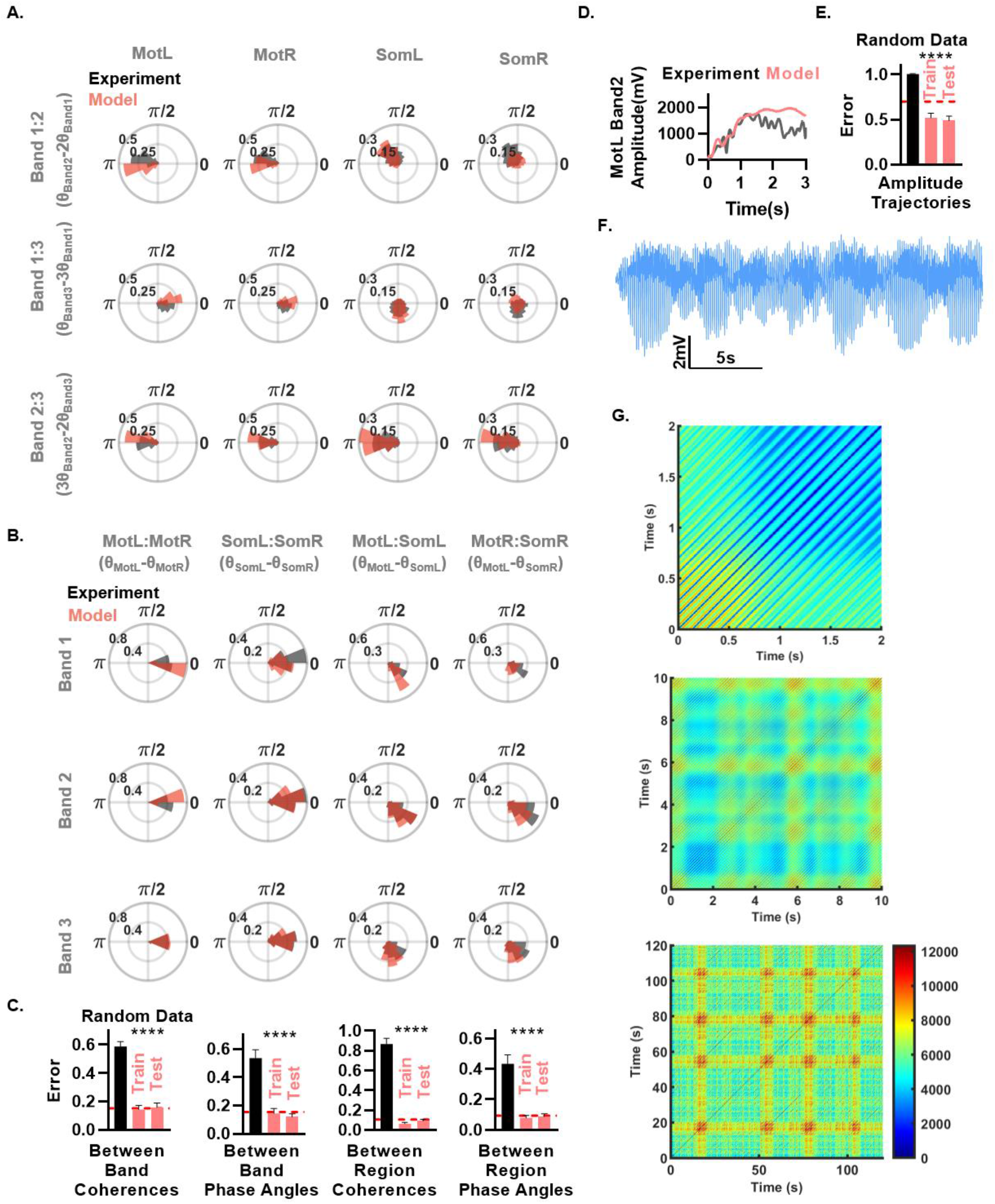
The model recapitulated evolution to the seizure state space. (A and B) Phase-histograms of experiment and model from the same initial conditions recapitulate phase-lags and coherence across bands (A) and regions (B). (C) Model error on coherence and phase-lag attractor space metrics. Red dashed line indicates the error metric computed between different experimental mice as if they were the results of simulation. Asterisks indicate significance relative to time shuffled data, all Train and Test p<0.0001. Train data corresponds to error of simulated data when the 400 ms before seizure onset to 800 ms after seizure onset were included in regression steps and used for performance on state-space error metrics used in library selection steps. Test data indicate novel seizures not included in regression or in library selection. (D) Example MotL band 2 amplitude trajectory showing initial tracking of the trajectory over ~1s. (E) Model error in tracking amplitude trajectories. Asterisks indicate significance relative to time shuffled data. Test and train as above, both p<0.0001 vs time shuffled data. Red dashed line as above. (F) MotL voltage trace over first 25 s of a 1200 s long simulation. Note variation in amplitude over seconds long time scales. (G) Recurrence plots of a simulation during the last 120 s of the above simulation. Color indicates Euclidian distance between states of the entire system at indicated time-lags. Scale bar in μV. Note the diagonal unity line at 0 time-lag indicates the 0 μV distance from a state to itself. Note the blue parallel diagonal lines at temporal lags occurring at multiples of the SWD interval (~133 ms), indicate that successive SWDs are similar, but not identical as amplitudes and phases subtly and chaotically vary from cycle to cycle across bands and regions. (Middle) Distances between state pairs over a longer range of temporal lags indicates aperiodic dynamics, with the system often coming near to previous states, but never identically back to a previous state. Note the strong blue diagonals at short temporal lags, even during periods of low similarity (yellow) indicating SWDs occurring in these states. (Bottom) Longer view of the last 120 s shows strong and transient aperiodic departures from previous states (red regions).

**Figure S4.**
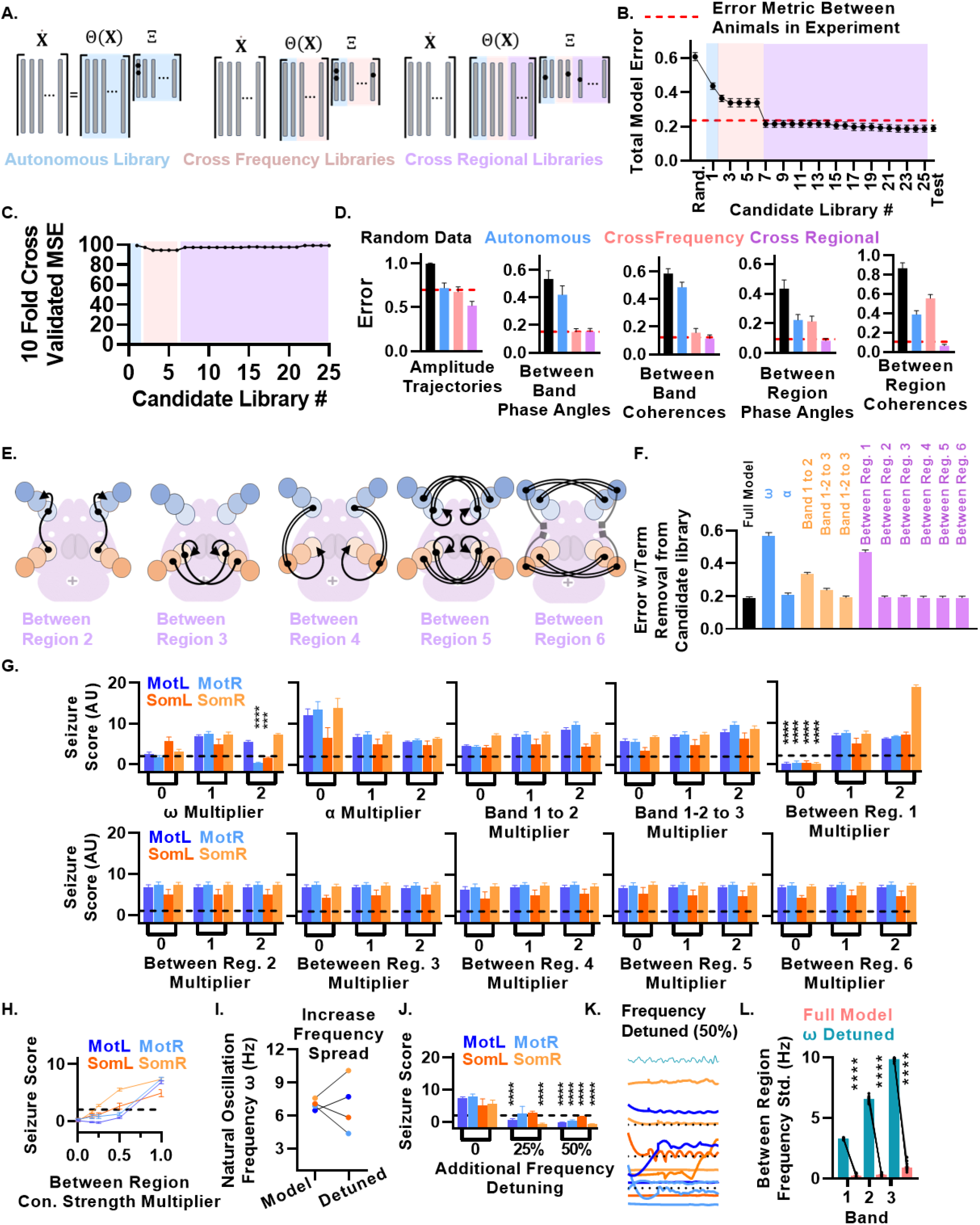
The model selection approach enables interpretable model performance metrics and model function. (A) Schematic representation of the modified SINDy (Sparse Identification of Nonlinear Dynamics) approach for identifying dynamical models from experimental data using candidate function library (θ**(X)**) built from various expansion steps to identify term coefficients (**Ξ**) that best describe the system’s dynamics 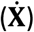. Broad classes of θ**(X)** at the level of autonomous functions, between-band functions, and between-regional functions shown in respective colors highlighting the appending of more complex library terms to those selected at lower complexity levels. (B) Total error metric (Supplementary Equation 9) for each tested library as defined in table S1. Red dashed line indicates the same error metric computed for experimental data between mice. Test indicates performance on held-out seizure data not included in regression or library selection steps. (C) Cross-validated mean square error for single step regression across libraries showing no basis for model selection between libraries which show improvement in (B). (D) Model error for amplitude trajectory, and mean phase angles and consistency between-band, and between-regions at different broad levels of candidate library model complexity showing improvement in specific metrics corresponding to library type (Autonomous libraries, between-band libraries, and between-region libraries). Red dashed line indicates same metrics computed between animals. (E) Schematics of coupling functions not shown in Fig. 4B. (F) The effect of removal of each function class on total model error when that function class is removed from the final candidate library. Note that model error increases most with removal of ω (from library 1), between-band 1-2 coupling (library 2), and between-region (library) coupling. Note the negligible effect of between-region function classes 2-6 (libraries 15, 17, 20-22), function schematics in (E). (G) Effect of multiplying each function class term coefficients by 0 (0), doubling them (2), and the learned model (1). Note only removal of between-regional coupling abolishes seizures. Also note that modification of other between-region function classes has a negligible impact on seizure score. Dashed line indicates seizure score of 2, i.e. amplitude growth below 2 z-score from starting condition. (H) Sweep weights of between-region within band coupling reveal total abolition of seizures at a 6-fold reduction. (I) Intrinsic frequencies ω in model for band 1 and 50% maximal detuning i.e. with oscillators detuned 50%, 25%, −25%, and −50% of the learned ω. Bands 2 and 3 set at corresponding multiples of their band 1 frequency. (J) Seizure score vs maximum frequency detuning. Asterisks indicate significant reduction below seizure score of 2, i.e. amplitude growth below 2 z-score from starting condition. Comparison via a 1 sample t-test. ****p<0.0001. (K) Representative voltage traces and oscillation frequency in each region for model with detuning as in I. (L) Standard deviation of frequency between regions as in Fig. 4E and G. n=25 simulations with initial conditions from 5 mice. Asterisks indicate significance via two tailed t-test. ****p<0.0001

**Figure S5:**
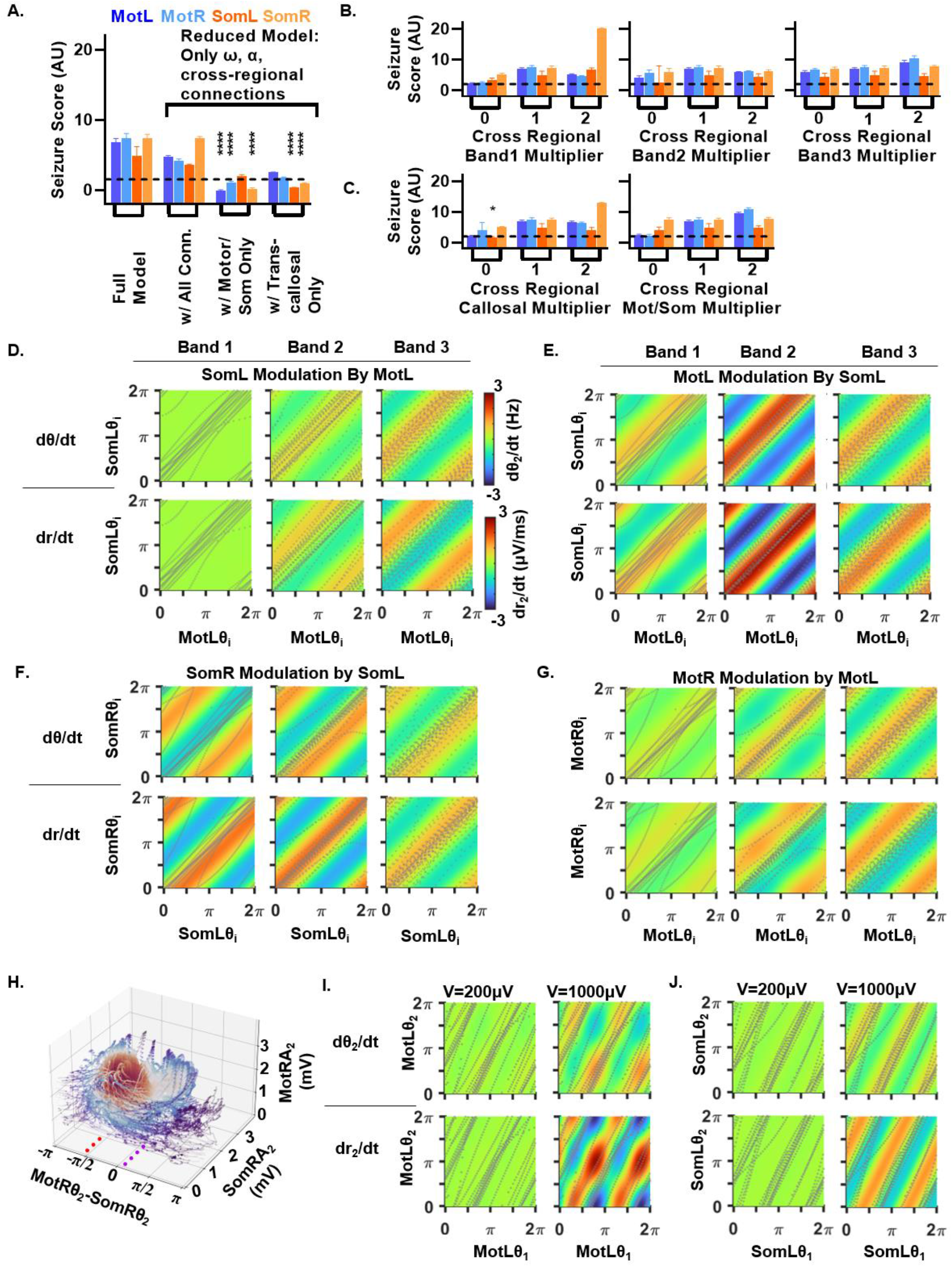
Between-region coupling functions enabled interpretation of drives behind dynamics. (A) Effect of reducing the model to include only the autonomous and stable oscillator terms (intrinsic frequencies ω and limit cycle α) and the seizure driving between-region coupling shown schematically in Fig. 4B. Note robust seizure generation comparable to full model. Further reducing the model by removing either Mot to Som and Som to Mot or by removing all transcallosal coupling reduces but does not abolish seizures. Asterisks indicate significant reduction below seizure score of 2, i.e. amplitude growth below 2 z-score from starting condition. Comparison via a 1 sample t-test. ****p<0.0001 (B) In the full model, removing or doubling the seizure-driving coupling terms in each band individually does not abolish seizures in isolation. (C) In the full model, removing or doubling either Mot to Som and Som to Mot or by removing all transcallosal coupling impairs but does not abolish seizures as in (A). (D) Phase and amplitude coupling functions for band 1, 2, and 3 for SomL modulation by MotL. Note the lack of Mot to Som coupling in band 1 while band 2 and 3 are stronger and both exhibit a relative phase offset in amplitude modulation toward −π/2 rad. Gray dots indicate a typical seizure trajectory. (E) Phase and amplitude coupling functions for band 1, 2, and 3 for MotL modulation by SomL. Note the strong coupling in band 1 in contrast to lack of coupling in band 1 in D. Also note that all coupling functions in E follow a similar pattern and amplitude modulation is nearly symmetrical through 0 phase offset across all bands. (F) Phase and amplitude coupling functions for band 1, 2, and 3 for SomR modulation by SomL. Note the strong coupling in band 1 in contrast to D. Also note that all coupling functions in F follow a similar pattern and amplitude modulation is nearly symmetrical through 0 phase offset across all bands. (G) Phase and amplitude coupling functions for band 1, 2, and 3 for MotR modulation by MotL. (H) Examination of band 2 trajectories along the chaotic attractor during long simulations. Dynamics of MotR-SomR coupling. Compare to similar coupling in Fig. 4M. (I and J) Coupling functions for band 1 to 2 in MotL (I) and SomL (J). Note the amplitude dependence of the coupling functions lead to negligible modulation at 200 μV but modulation comparable to between-regional connections at 1000 μV, indicating why band 1-2 phase relationships are only observed during seizure.

**Figure S6.**
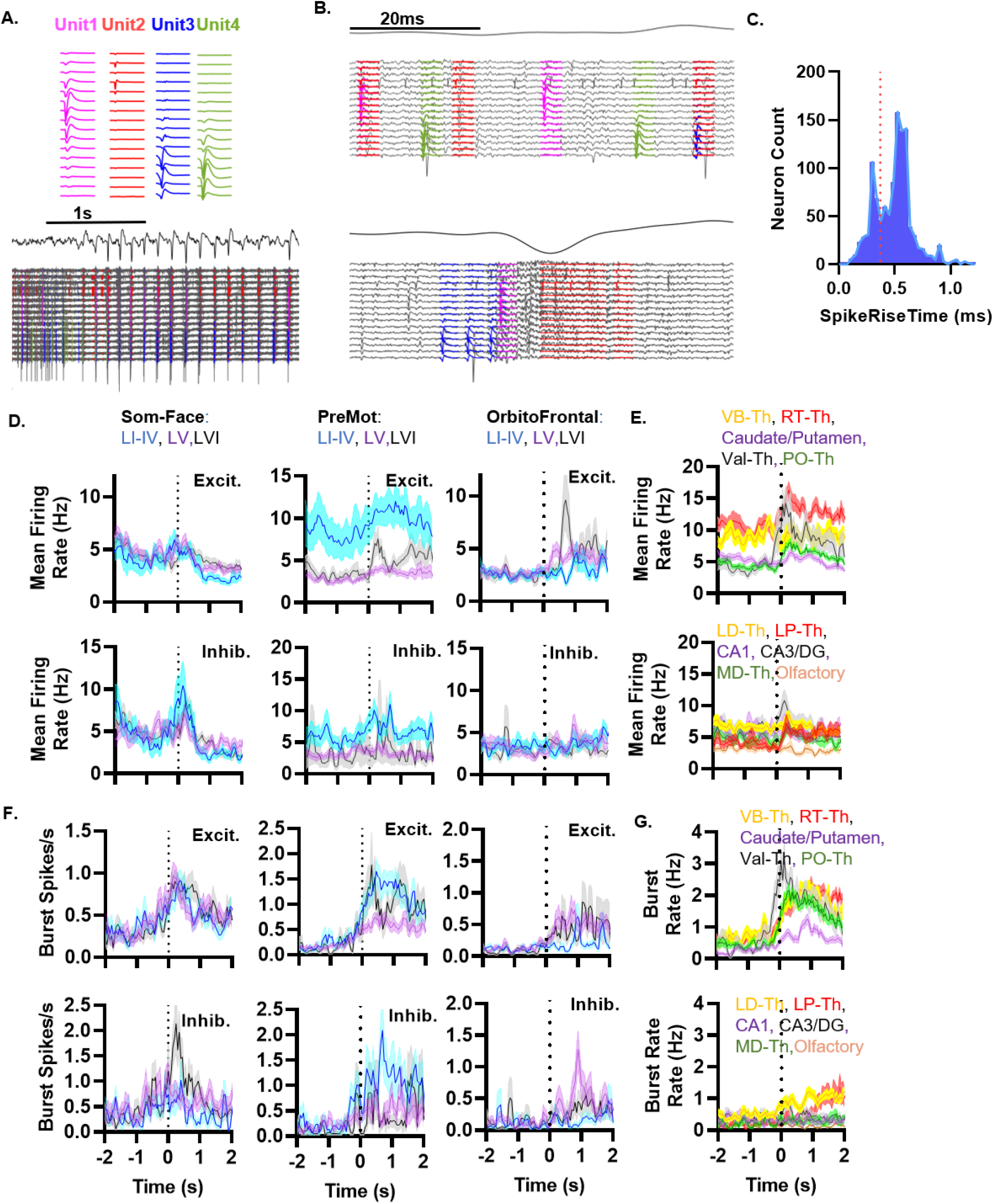
Spike Sorting and basic spike rate parameters across recorded populations. (A) Example average waveforms of 4 identified single units in PO on the same recording channels (top) and example seizure Ecog trace (middle) and example recording channels with units marked by color. Note dispersed firing just before seizure and spikes aligned to SWD-spike with near total silence between SWDs. (B) Expanded view of spiking outside of seizure in A. Expanded view of spiking during the SWD-spike and relative lack of spiking outside the SWD-spike (bottom). Also note while there is period of no detected units spanning 3.7 ms during the spike, there is prominent detection of multiple high-frequency spikes (i.e. bursting) of stable waveform during the SWD. (C) Histogram of single unit spike widths in PreMot and Som cortex used to differentiate excitatory (right of the dashed line) from inhibitory (left of the dashed line) units. (D and E) Mean firing rates across populations leading up to and during absence seizures. Note the lack of large changes in mean firing within any population with the largest percent change being a 2.8 fold increase in Val thalamus (E). Note the *decreased* firing in Som:L5 and L6 during seizures. (F and G) Count of bursts/s across regions. Bursts defined as >40Hz interval preceded by <10Hz interval. Note the >4-fold increases in many regions. Note also the delayed onset or absence of burst firing in orbitofrontal cortex and LD, LP, MD thalamus as well as CA1, CA3/DG and olfactory areas.

**Figure S7.**
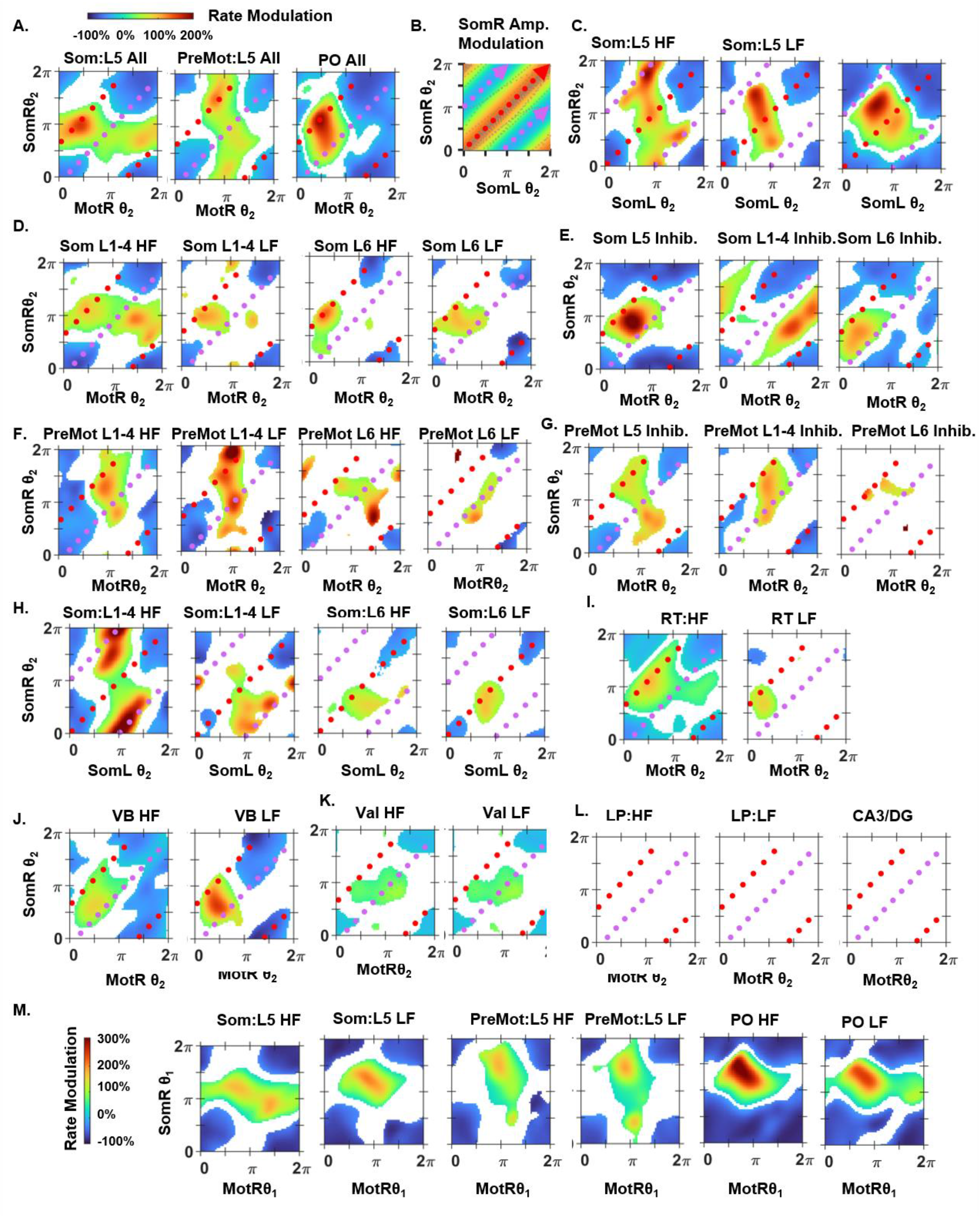
Model coupling functions inform firing pattern across different populations. (A) Rate modulation of all spiking (burst and tonic firing) in populations from Fig. 6. (B) Coupling function for SomL modulation of amplitude in SomR band 2. Peak amplitude growth marked by red and decay by purple trajectories. (C) Firing along trajectories in (B). Som:L5 high-frequency (HF) firing (left) follows a similar pattern similar pattern as shown in Fig. 6 with increases (red region) and decreases (blue region) along the amplitude growth trajectory (red trajectory) for the SomL to SomR connection in the model and an increase in HF firing without accompanying silencing along the amplitude decay trajectory (purple trajectory). Low-frequency (LF) firing (middle) increased and decreased along the growth trajectory and was not modulated along the decay trajectory (purple trajectory). PO HF firing (right) showed a similar pattern as PO in Fig. 6 with rate modulation increasing (red region) and then decreasing (blue region) along the growth trajectory while the decay trajectory (purple trajectory) was not strongly modulated. (D) Som:L1-4 and Som:L6 followed similar but weaker pattern of modulation as Som:L5 observed in Fig. 6E. (E) Som Inhibitory units followed distinct patterns from excitatory neurons across layers. Som:L5 inhibition (left) was strongly positively modulated (red region), occupying the region between trajectories and showed silencing (blue regions) along all trajectories. Som:L6 (right) followed a weaker but similar pattern to Som:L5. Som:L1-4 (middle) inhibitory firing followed a distinct pattern from both Som:L1-4 excitatory firing (D, left) and inhibitory firing in Som:L5 and L6. Note the Som:L1-4 inhibitory unit silencing along both trajectories and strong activation outside of the seizure state space. (F) PreMot:L1-4 and L6 followed more distinct patterns from PreMot:L5 observed in Fig. 6G. Notably, L1-4 exhibited burst behavior along both trajectories while L6 was weakly modulated. (G) PreMot:L5 inhibition (left) peaked along Som growth trajectories and was silenced during Mot Growth trajectories while PreMot:L1-4 firing increased and then was silenced along both trajectories. PreMot:L6 was weakly modulated. (H) Som:L1-4 HF firing was weakly activated (green region) and then silenced (blue region) along the SomL-SomR (coupling function shown in B) growth trajectory (red trajectory) and LF firing was strongly activated along the amplitude decay trajectory (purple trajectory). Som:L6 HF and LF was moderately activated and silenced along the growth trajectory. (I) The thalamic reticular nucleus (RT) exhibited increased HF firing followed by silencing along the Som growth trajectory (red trajectory) and modest HF firing along the Mot growth trajectory (purple trajectory. Note however RT contains neurons associated with diverse brain systems including motor, sensory and limbic systems complicating interpretation. (F and K) Ventrobasal thalamus (VB) showed burst firing between trajectories, similar to Som:L5 and Som:L6 inhibition suggesting a role in feedforward inhibition. Note overall weaker association with the torus than PO (Fig. 6J). (K) Ventral anterolateral thalamus (Val) showed bursting along the Mot growth trajectory. (L) There are regions with firing uninformed by the phase torus. Lateral posterior (LP) thalamic nucleus, a higher order vision related nucleus, and CA3/DG exhibited no modulation on the band 2 Mot-Som phase torus. (M) Band 1 Mot-Som phase torus exhibits modulation of Som:L5, Mot:L5 and PO firing. Modulation of Som:L5 indicates it predominantly represents coupling with band 2 due to the presence of multiple peaks, as predicted by the lack of band 1 coupling from Mot to Som (Fig. S5D).

**Figure S8.**
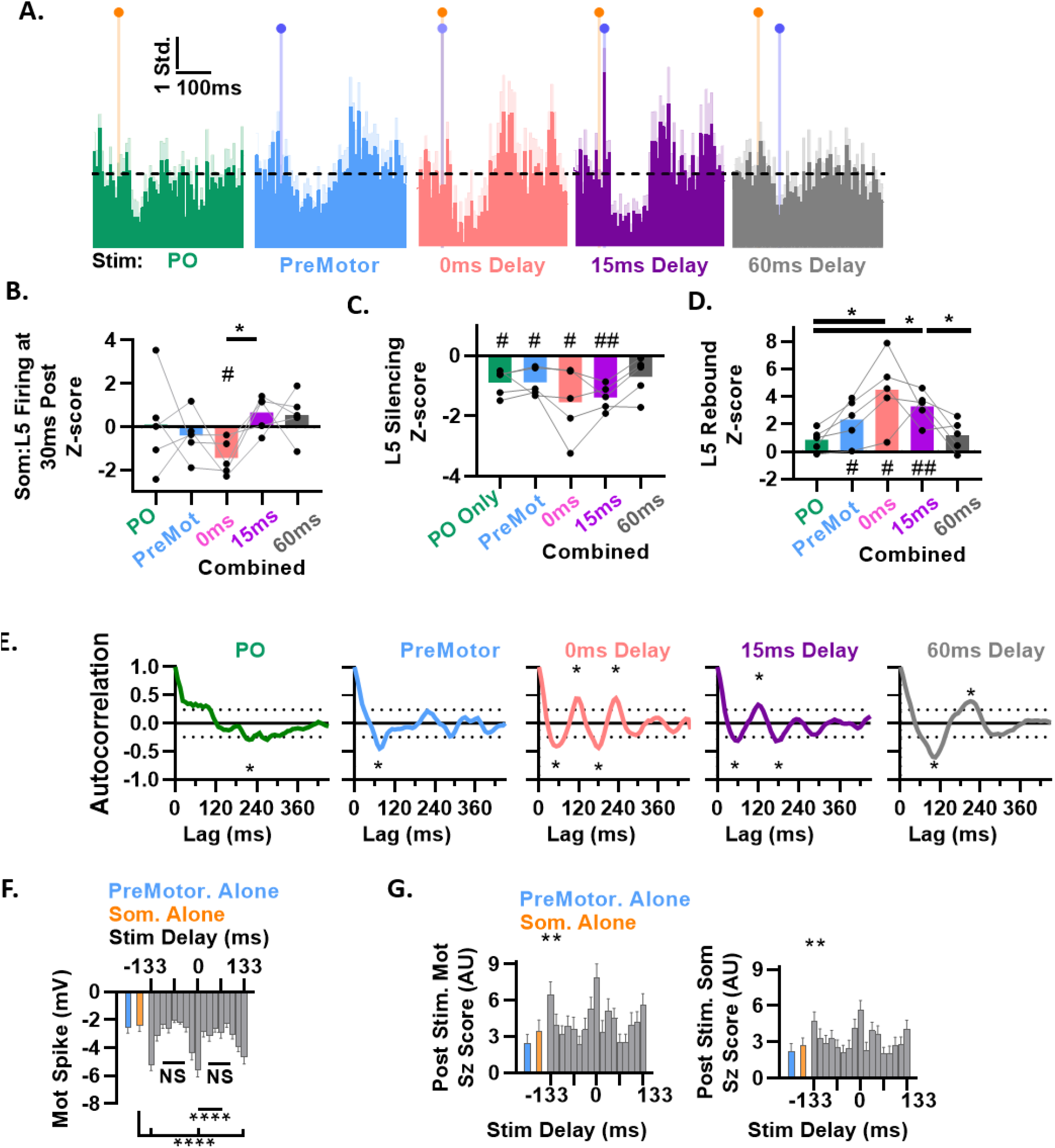
Combined optogenetic stimulations induce seizure rhythms. (A) Extended time view of PSTH plots from Fig. 7F. (B) Firing of Som:L5 neurons at 30 ms after initial stimulation onset. Note robust silencing with the 0 ms delay stim, consistent with 0 ms delays promoting spiking suppression after the (nonsignificant) peak occurring at 15 ms in Fig. 7G. In contrast, the 15 ms delay, which caused a large increase in firing at the 15 ms delay, is significantly higher than the 0 ms delay at 30 ms post initial stim of PO. PO alone, PreMot alone, or 60 ms delay do not cause silencing at this early timepoint. # indicate one-way t-test vs 0, asterisk indicate between group comparisons via paired t-test). (C) All stimulations except 60 ms delay produce Som:L5 silencing (average at 45:90 ms post initial stimulation). # indicate one-way t-test vs 0, asterisk indicate between group comparisons via paired t-test). (D) PreMot alone, 0 ms delay, and 15 ms delay produced a rebound increase in Som:L5 firing. 0 ms and 15 ms were elevated relative to PO or 60 ms stimulation while 0 ms was additionally elevated relative to PreMot. # indicate one-way t-test vs 0, asterisk indicate between group comparisons via paired t-test). (E) Autocorrelation of the average PSTH following the silencing period. 95% confidence bounds for the were computed using Bartlett’s formula under a white-noise null. Only the 0 and 15 ms delay produced seizure interval oscillations (~120 ms). The 60 ms delay produced a slow oscillation near 2x the seizure interval. Asterisk indicate >two consecutive points above the 95% confidence bounds. (F) Average minimum spike amplitude across 10 trials under each phase-lag condition for Mot as in Fig. 7L. Lines show pairwise t-tests. (G) Seizure score in the 1 second after stimulation end. Asterisks indicate significance by ANOVA. **p<0.01.

## Supplemental Methods

### Animals

Heterozygous loss of function mutation in *Scn8a* (*Scn8a*+/−) mice were purchased from Jackson laboratory (C3HeB/FeJ-*Scn8a*med/J, Kohrman et al., 1996, Stock#: 003798) and maintained on the C3HeB/FeJ background. For optogenetic experiments, homozygous *Thy1-mhChr2-YFP* (B6.Cg-Tg(Thy1-COP4/EYFP)18Gfng/J, Stock# 007612) were purchased from Jackson laboratory and crossed with *Scn8a*+/− mice to generate *Scn8a*+/−; *Thy1-mhChR2-YFP*+/− mice. Mice were genotyped at postnatal day 21 through the Transnetyx corporation. Mice were maintained on a 12 hr light/dark cycle and male and female mice were used in all experiments. Experiments were approved by the Stanford Administrative Panel on Laboratory Animal Care (APLAC, Protocol #12363) and in accordance with the National Institute of Health guidelines.

### Stereotaxic surgeries

For all surgeries mice were p56-90. Mice were anesthetized with an isoflurane oxygen mixture at 4% for induction of anesthesia and 1-2% for maintenance. Subcutaneous Rimadyl (5mg/kg) was administered after induction and for three days following surgery. Mice in Fig. 1 were implanted with 16 gold electrodes (1.27mm, Mill-Max Manufacturing Corp) in contact with the dura in the locations shown in Fig 1A. Coordinates relative to Bregma for electrodes following the numbering in Fig. 1A in mm are; (1,2): 2.68 AP, ±1 ML, (3,4): 1.68 AP, ±2 ML, (5,6): 0.3 AP, ±3.5 ML, (7,8): −0.82 AP, ±3.5 ML, (9,10): −1.82 AP, ±3.5 ML, (11,12): −0.82 AP, ±1.5 ML, (13,14): −1.82 AP, ±1.5ML, (15,16): −2.8 AP, ±2.5 ML. All Ecog signals throughout the manuscript were referenced to a stainless steel Ecog screw (J.I Morris Company FF00CE125) implanted in the skull above the cerebellum. Mice were also implanted with a custom designed head bar (eMachineShop). Headcaps were constructed using dental cement (Metabond).

For Ecog and Neuropixels recordings with two site optogenetic stimulation of premotor cortex and PO with combined Neuropixels recording from somatosensory cortex *Scn8a+/−; Thy1-mhChR2-YFP*+/− mice were used. The red shifted opsin ChrimsonR (pAAV-Syn-ChrimsonR-tdT (AAV1), 1×10^13^ vg/ml) was delivered to PO as described previously at coordinates (−2.06AP, +1.25ML,-3.2DV).^63^ During optogenetic experiments an optic fiber was connected to the optic fiber implanted on the mouse. Blue and yellow laser light were delivered by OEM Laster Systems 473 nm and Cobolt Mambo 594 nm lasers. Pulses were 5 ms wide and laser power titrated to generate small local LFP responses in the target region. Pilot experiments found ChrimsonR virus generated seizures even at very low light stimulation intensity, so we diluted 10x in saline. Injection volume was 400 nL delivered at a rate of 50 nL/min. An optical fiber (200 μm, 0.48-NA, 1.25 mm-diameter stainless steel ferrule, Doric Lenses) was then implanted 500 μm above the injection site and a separate fiber in L5 of premotor cortex (coordinates +2.68AP,+1.5ML,-0.5DV). Gold electrodes as above were placed adjacent to the optical fiber in contact with the dura at (2.68AP, +1.0 ML) and two additional electrodes were placed on the contralateral side (coordinates: +2.68AP, −1ML) and (coordinates −0.0 AP, −4ML). Future Neuropixels recording site was marked (coordinates: 0 AP, 4ML). Mice were allowed to express the virus for a minimum of 3 weeks before recording. For Ecog and dual site optogenetics stimulation of premotor and somatosensory cortex *Scn8a+/−; Thy1-mhChR2-YFP*+/− were used. Two optical fibers as above were implanted at coordinates (+2.68, 1.5 and 0, 3.5) with adjacent gold pins at (+2.68AP,1ML and 0AP, 4.0ML). Optogenetic stimulations were excluded from analysis if preceded by a spontaneous seizure.

For acute Neuropixels recordings, an initial surgery implanting a headbar and reference screw were placed as in the Ecog surgeries. The surface of the skull was marked for future craniotomy sites at sites 1: 2.68 AP, 1.5ML; 2: −0.78AP, 4ML; and 3-2.06AP,1.5ML and small ridge were constructed to create a well for recording. A stainless-steel screw for Ecog recording and additional headcap stability was implanted in the contralateral somatosensory cortex was implanted at −0.78A/D, −4ML. Following one week of recovery, mice were trained for wheel running as in Ecog mice. On the day of recordings, mice were briefly anesthetized, and 3 craniotomies were performed at the marked locations, creating ~0.5 mm diameter craniotomies. Craniotomies were filled with DOWSIL silicon gel (DOW) and covered by KWIK-SIL silicone elastomer (World Precision Instruments).

### Electrocorticogram and Neuropixels recordings

For 16 channel Ecog recording, animals were allowed to recover for 1 week and then were acclimated to head fixed wheel running over the course of 1 week. Ecog recordings were performed with head fixed freely running mice in a dark and sound insulated chamber for 1 hour of Ecog recording. For mice with viral injections, three weeks were allowed for virus expression prior to recording. The Ecog electrodes and ground/reference were connected to an Intan headstage (RHD 2132) and OpenEphys acquisition board and sampled at 30 kHz. Data were recorded at 30 kHz using the OpenEphys software and recording system and down sampled to 500 Hz for analysis.^64^

Neuropixels 1.0 probes were coated in DiD, DiI, or DiO dyes prior to insertion. 12-16 hours after craniotomy surgeries, mice were head fixed and the silicon elastomer was removed. Craniotomy sites were washed with saline and surface moisture was wicked away. 3 probes were inserted at an angle of 18 degrees (site 1), 45 degrees (site2), and 0 degrees (site 3) from vertical. Craniotomies were then filled with silicone oil. Probes were intermittently advanced using Sutter MP-285 manipulators at a rate of ~10 μm per second with over the course of 5 minutes. Insertion continued until 384 μm was traversed along the insertion axes. Zero was marked as the measure of the surface of the dura where the 384^th^ electrode contacted conductive tissue at the surface of the brain. Probe ground and reference were shorted and referenced to the cerebellar Ecog screw. Acquisition utilized a National Instruments PXIe-1071 Chassis and PXIe_1000 base station card and recordings were performed with OpenEphys software. Voltages were sampled at 30 kHz and filtered between 300 Hz to 10 kHz for spike band and voltages were sampled at 2500 Hz for LFP band information.

Probes were allowed to rest for 15 minutes prior to the start of acquisition and 3 bouts of recordings of 15 minutes were acquired. Seizures were selected as above for Ecog recordings from the final 15 minute recording bout and additional seizures were selected from the previous bout if there were insufficient number of seizures in the third recording bout. Following recording, probes were removed and craniotomies sealed with Kwik-Sil or probes were repositioned for a second bout of recording in different probe placement locations. A subset of mice were then recorded again with probe repositioning or 24 hr later with an alternate probe dye-coloring at each location. Mice were used in a maximum of two recordings, whether sequential on the same day or after 24 hours.

### Neuropixels probe tract reconstruction

Mice were euthanized with intraperitoneal injection of Fatal+ (Pentobarbital, Vortech Pharmaceutical). We then intracardially perfused 10 ml PBS followed by 10 ml PBS with 4% paraformaldehyde (PFA). Brains were stored overnight in PBS with 4% PFA, washed and stored in PBS until slicing. Serial coronal sections (50 μm) were made on a vibratome (Leica VT1000s). All sections containing probe tracks were mounted in DAPI-Fluoromount-G (Southern Biotech 0100-20). Whole section images were taken spanning all probe tracks on a Zeiss Axio Imager M2 microscope. SHARP-Track (Allen mouse brain common coordinate framework) software was used to reconstruct the 3 dimensional probe locations from the surface of the brain to the tip of each electrode. Regions traversed were assigned to the Allen mouse brain common coordinate framework. Histological boundaries and physiological markers were then used to assign channels to different brain regions. Several functionally similar or difficult to distinguish divisions based on physiological markers and histology are grouped as common structures. The number of animals, distinct probe insertions into a region (Hits). Regions with fewer than 10 neurons were excluded as a hit.

Traversed regions included: (ventrobasal thalamus VB = 4 mice, 8 Hits, 147 neurons); (Reticular thalamic nucleus RT = 4 mice, 7 hits, 389 neurons); (posterior thalamic nucleus PO = 4 mice, 6 hits, 291 neurons), (Mediodorsal thalamus MD = 3 mice, 3 hits, 162 neurons), (ventral lateral thalamus Val = 4 mice, 5 hits, 178 neurons); (lateral dorsal thalamus LD = 4 mice, 6 hits, 339 neurons); (lateral posterior thalamus LP = 3 mice, 4 hits, 90 neurons); (Basal ganglia = 7 mice, 10 hits, 243 neurons); (Som:L1-4 = 7 mice, 10 hits, 121 neurons); (Som:L5 = 7 mice 11 hits, 154 neurons), (Som:L6 = 8 mice, 11 hits, 154 neurons), (CA1 = 5 mice, 7 hits, 68 neurons); (CA3/DG = 4 mice, 7 hits, 60 neurons); (PreMot:L1-4 = 7 mice, 10 hits, 115 neurons); (PreMot:L5 = 7 mice, 9 hits, 115 neurons); (PreMot:L6 = 4 mice 5 hits, 162 neurons); (Orbitofrontal cortex:L1:4 = 5 mice, 7 hits, 352 neurons); (Orbitofrontal cortex:L5 = 5 mice, 7 hits, 278 neurons); (L6 Orbitofrontal cortex:L6 = 3 mice, 4 hits, 277 neurons); (Olfactory areas = 4 mice, 6 hits, 221 neurons).

### Ecog spectrogram and filter construction

We computed the spectrogram using Matlab’s spectrogram function with an interval of 500 samples, 95% overlap, and an FFT length of 10*500 samples. We constructed ideal passband FIR filters with Matlab’s firls function with bands centered at 7 Hz, 14Hz, and 21 Hz with a spread of ± 4 Hz, a transition width of 2 Hz, and order 500 samples resulting in filter amplitudes shown in Fig. 2A. Filters were applied using Matlab’s filtfilt function.

### Seizure score and Cox proportional hazard model

Seizure score was calculated using custom MATLAB software. We first computed the amplitude for each band and region using the Hilbert transform. A baseline period outside of seizures was used to compute the standard deviation of amplitude for each of the 3 bands. A moving gaussian filter of width 250 ms was applied to remove sharp transients and then seizure score was defined as the average of the 3 band amplitude z-scores. Seizures were then automatically detected as an increase of seizure score over 3 for a period longer than 3 s. Seizure onset time for each electrode was defined as elevation above band averaged z-score of 1 with a subsequent cross of 3 for at least 3 seconds without dropping back below 1 prior to crossing 3. Seizure offset time was defined as the last electrode to fall below a seizure score of 2. Cox proportional hazard model was performed in MATLAB using the coxphfit function on onset times from 30 seizures per mouse from 5 mice.

### Measures of Ecog phase and frequency

Phase-coherence was calculated as

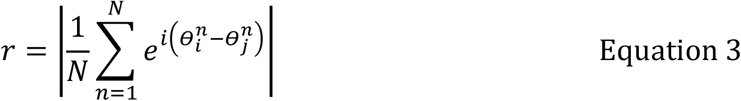

Where 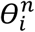 and 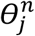 denote the phase in regions i and j at time point n. For calculation of instantaneous oscillation frequency, the phase angles computed from the Hilbert transform were unwrapped and a 4^th^ order Savitzky-Golay filter was applied with a window size of 1.33, 0.66, and 0.45 s for bands 1, 2, and 3 respectively to cover ~10 oscillation cycles and the rate of change was calculated via the finite difference.^65^

### Conversion calculations between SWD-spike time-lags and oscillation phase-lags

To compare the relationship between SWD-spike time-lags and the band specific phase-lags we calculated the mean lag and the 95% range (corresponding to the range of 2 standard deviations). We began by first converting time values of SWD-spike time-lags between regions to the corresponding predicted phase-lags as follows. The mean MotL-SomL SWD-spike time-lag was −8.2 ± 0.9 ms, indicating SomL spikes typically occur before MotL spikes (example traces Fig. 2K, left, gray circles Fig. 2K, right) with 95% of lags between −19.6 and +3.3 ms (Fig. 2K purple and red circles respectively). We compared this lag to the relative phase-lags of MotL and SomL band 2 (MotLθ_2_-SomLθ_2_, schematic and example traces, Fig. 2L). Since band 2’s frequency is 15Hz (a period of 67 ms), SWD-spike lags of −8.2 ms divided by the period of 67 ms times 2π rad, should correspond to a phase-lag of −0.77 ± 0.1 rad or approximately −π/4 rad (about 1/8th the band 2 period). Similarly, the 95% range of SWD-spike temporal lags from −19.6 ms to +3.3 ms, corresponds to band 2 phase-lags ranging from −1.84 to +0.31 rad, or equivalently from slightly less than −π/2 rad to slightly more than 0 rad. As predicted, the directly measured MotLθ_2_-SomLθ_2_ phase-lag exhibited a mean of about −π/4 rad (−0.80 ± 0.10 rad) during seizures, with 95% interval between −1.86 ± 0.29 rad and 0.27 ± 0.15 rad (gray, red, and purple trajectories and corresponding circles in Fig. 2M and N). The ~ 0 rad phase-lag in band 2 for MotLθ_2_-MotRθ_2_ (−0.07 ± 0.05 rad, Fig. 2P schematic and histogram) and SomLθ_2_-SomRθ_2_ (0.07 ± 0.11 rad, Fig. 2R schematic and histogram). The MotL-MotR SWD-spike time-lag 95% range (from −7.0 to +5.6 ms purple and red circles, Fig. 2P) predicted a MotLθ_2_-MotRθ_2_ range of −0.66 to 0.53 rad, paralleling the observed phase-lag (−0.42 ± 0.11 and +0.27 ± 0.05 rad, Fig. 2Q purple and red circles). The 95% range for SWD-spike time-lags in SomL-SomR was −9.3 and +10.1 ms (Fig. 2R, purple and red circles), predicting a phase-lag range of −0.87 and 0.95 rad, paralleling the observed SomLθ_2_-SomRθ_2_ range (−0.74 ± 0.13 to 0.89 ± 0.23 rad, Fig. 2S, purple and red circles).

### Spike sorting and Neuropixels data quantification

Spike sorting was performed as outlined previously.^63^ Individual units were sorted using Kilosort2 (https://github.com/MouseLand/Kilosort). Unit clusters were manually curated using Phy2 (https://github.com/cortex-lab/phy). Single units with interspike interval violations >5% percent were excluded from the analysis. Clusters with >20% contamination percentage with neighbor clusters were excluded. Mean firing rate was computed in one second long bins in a 250 ms long sliding window as the number of spikes in a population divided by the number of neurons. Inter-spike intervals distributions were computed for each neuron in a 2 second long window. The distribution of ISIs in the population was then computed for each seizure at times T=-10 s before seizure, T=0, and T=1s after seizure onset. ISIs were converted to frequency as 1/ISI. Probability-normalized histograms were computed using MATLAB’s histcounts function. Bin edges spanned from 1 to 1054 Hz in natural log spaced bins of widths of 0.06 (117 bins). The resulting histograms represent the probability distribution of instantaneous firing frequencies of individual neurons in the above time windows. Frequency clustering was computed by finding the peak of the distribution and computing the peak prominence as the difference between the probability peak and 7 frequency bins away (average of each side of the peak). Distributions for individual seizures with less than 20 occupied bins were excluded. To compute firing on phase tori, for each seizure, spike times in the peri-onset window (−0.5 to +0.5 s relative to seizure onset) and phases were extracted from simultaneously recorded local field potentials in the different brain regions (Somatosensory cortex, PreMotor cortex, and screw electrode). We binned the phase tori in a 60 by 60 grid and computed a 2D histogram of spikes on this grid. To control that some phase combinations occur more rarely along the torus but may highly promote spiking within individual regions we computed the distribution of oscillation phases sampled at all time points within the same analysis window (−0.5 to +0.5 s relative to seizure onset), providing an occupancy reference that controls phase availability independent of spike timing. For each region and phase band, phase distributions were concatenated across seizures and neurons, and normalized probability histograms were constructed.

The expected spike counts per bin were computed by scaling the occupancy map to the total number of observed spikes. Both the observed and expected maps were circularly smoothed (2D Gaussian; σ = 4 bins; wrap-around on both axes). We then calculated a per-bin modulation index (observed number of spikes - expected number of spikes) /expected number of spikes). Statistical significance was assessed by a multinomial shuffle test (1,000 surrogates of spike counts drawn from the occupancy probabilities), yielding two-sided p-values and 95% null confidence intervals; multiple comparisons were controlled with Benjamini–Hochberg FDR at q=0.05. The heat map displays the modulation index.

### SINDy method

We sought to identify models in the form:

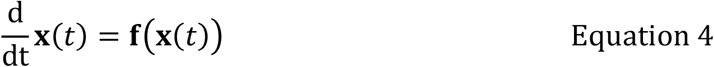

The vector **x**(*t*) ∈ ℝ^*n*^ denotes the system state at time t which consist of the real (x) and imaginary (y) components of the analytic signals z_R,B_=x_R,B_+iy_R,B_ as defined in Fig. 2 and where R=1,2,3,4 represents regions MotL, MotR, SomL, and SomR respectively and B=1,2,3 represents frequency bands 1,2, and 3 respectively. Given 4 regions times 3 bands times 2 components (real and imaginary), the dimensionality n of our dynamical system is n=4*3*2=24.The data are sampled at m time points t_1_,t_2_,…,t_m_ and represented by an m by n matrix **X** with each row **x**^T^ being an n dimensional system state at a given time:

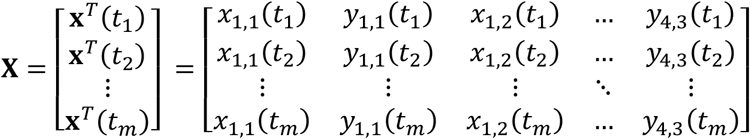

Derivatives were approximated by the central difference method as (**x**^T^(*t*_i+1_)-**x**^T^(*t*_i-1_))/(2Δt) and represented by the matrix

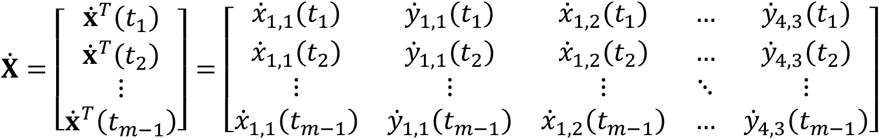

Next we constructed a candidate function library Θ(**X**) consisting of functions of the columns of **X**.

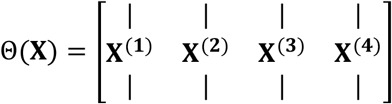

Where **X**^(1)^ denotes linear terms of x_R,B_ and y_R,B_,

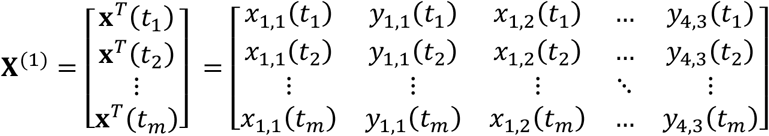

**X**^(2)^ denotes products of x_R,B_ and y_R,B_ in each possible combination in second order.

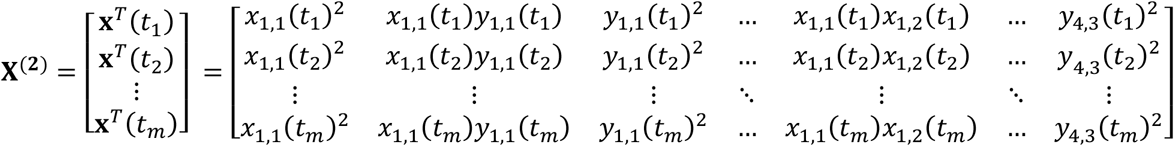

And **X**^(3)^ and **X**^(4)^ denote similar matrices in 3^rd^ and 4^th^ order respectively.

We then set up a sparse regression problem to determine sparse vectors of coefficients *Ξ* = [ξ_1_ ξ_2_ … ξ_n_] that determines active terms in the model:

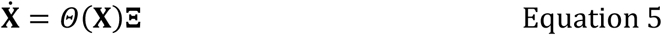

Each column of *Ξ* denoted by ξ_k_ is then a vector of coefficients determining the active terms in 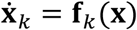. Governing equations are then represented as

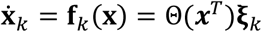

Where Θ(**x**^*T*^) is a vector of symbolic functions of **x**.

### Regression and model error

For each library an independent elastic net regression was performed to solve for **ξ**_k_ for each real and imaginary component of each region and band separately. Regressions used MATLAB’s LASSO function with a λ of 0.04 and an α of 0.9, determined in exploratory steps for parameter ranges which identified stable models. Regression used 10 fold cross-validation. Regressions were performed on the concatenated data of 25 seizures per mouse from 5 mice. Seizure data were from 400 ms before seizure onset to 800 ms after seizure onset (600 samples) for each seizure. 25 simulations (with initial conditions from 5 per mouse) were performed from experimental starting conditions 400 ms before seizure onset. Simulations used MATLAB’s ode45 function.

Five measures of model accuracy were used in SINDy library optimization. Total model error was calculated as the average of 5 error measures computed from 25 simulations.

Amplitude correlation error *AC*_*error*_ was computed as

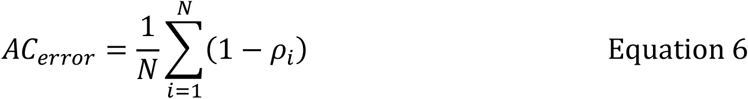

where ρ_i_ is the Pearson correlation coefficient between the experimental and model oscillation amplitude time series for oscillator *i*, and the sum runs over all N = 12 oscillators (4 regions × 3 frequency bands, schematic Fig. S2H, right).

For mean phase-lag between model and experimental distributions we first computed the phase differences *φ* = (*aθ*_*i*_ − *bθ*_*j*_), where *i* and *j* are oscillator indices and *a* and *b* are band ratios between angles *θ*_*i*_ and *θ*_*j*_ as in Fig. 2 and Fig. S3AB. These were computed for experiment (*φ*_Exp_) and model (*φ*_Model_), with the distributions for all compared relationships shown in Fig S3AB. Mean phase-lag angle error Lag_*error*_ was then computed separately for between-band Lag_*error*(*between*−*band*)_ and between-region Lag_*error*(*between*−*region*)_ in each case as

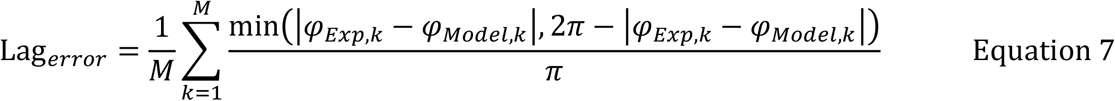

where k indexes each of the M phase-lag relationships for the between-band (Fig. S3A, schematic Fig. S2H, left) or between-region (Fig. S3B, schematic Fig. S2H, middle) relationships. Each term is normalized to π, since the absolute circular difference spans half the circle, giving an error between 0 and 1.

Phase coherence error *Coh*_*error*_ was then similarly computed separately for between-band *Coh*_*error* (*between*−*band*)_ and between region as *Coh*_*error* (*between*−*region*_)

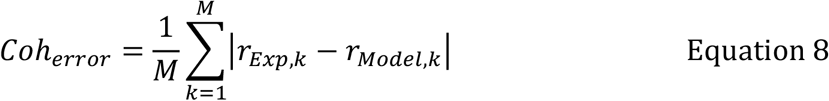

where r is the coherence computed using equation 3 and k indexes each of the M coherence relationships for the between-band (Fig. S3A, schematic Fig. S2H, left) or between-region (Fig. S3B, schematic Fig. S2H, middle) relationships.

Total error was then defined as the average of between these errors as

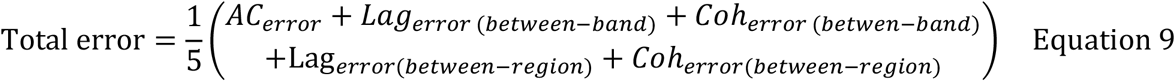

For comparisons to between-mouse variability, the errors computed between mice were computed as above replacing model for traces from each alternate mouse.

### Lyapunov spectrum

The Lyapunov spectrum was computed using a tangent-space method with repeated QR re-orthonormalization, following Benettin et al. (1980) and Shimada & Nagashima (1979).^66, 67^ The state equations were integrated jointly with the variational equation for the tangent propagator **Φ**(*t*) ∈ ℝ^*n*×*n*^,

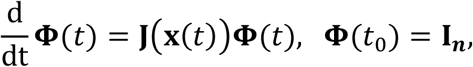

where **J** = ∂**f**/ ∂**x** is the Jacobian of the vector field and **I**_***n***_ is the identity matrix. Both **f** and **J** were derived symbolically (MATLAB Symbolic Math Toolbox).

The augmented (*n* + *n*^2^)-dimensional system was integrated with MATLAB’s ode45 with a relative tolerance of 10^−9^ and absolute tolerance of 10^−12^. To prevent the tangent vectors from collapsing onto the direction of maximal expansion, the integration was performed in chunks of length *τ* = 0.4 ms. At the end of each chunk the tangent matrix was re-orthonormalized via QR decomposition, **Φ**(*t*) = **QR**, with the columns of **Q** carried forward as the new orthonormal tangent basis. The signs of the diagonal of **R** were fixed to be positive before taking logarithms, so that sign flips introduced by the QR factorization do not contaminate the accumulated sums.

The Lyapunov exponents were then obtained as the time averages of the logarithmic expansion rates,

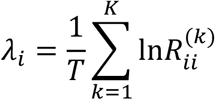

where 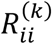 is the *i*-th diagonal element of the upper-triangular factor at the *k*-th renormalization step and *T* is the total accumulation time (2000 s). An initial transient of 8 s was discarded before accumulation began, so that the trajectory had relaxed onto the attractor. The initial conditions were drawn independently for each real and imaginary component from a uniform distribution between −150 to 150 μV.

The sum of all exponents (−26.427 ± 1.412 /s, mean standard deviation) was used as a check for dissipative dynamics, as negative sums signify dissipative systems indicative of an attractor. There were two positive exponents (0.255 /s ± 0.010 and 0.073 /s ± 0.004, mean and standard deviation respectively) indicative of chaos, and the third exponent was close to 0 (0.001 /s ± 0.002, mean and standard deviation respectively) indicative of the neutral flow direction. To estimate the attractor’s fractal dimension, we computed the Kaplan–Yorke dimension *D*_*KY*_. The exponents were sorted in descending order, and their cumulative sum was calculated (5.480 ± 0.117, mean and standard deviation respectively). *D*_*KY*_ was computed as

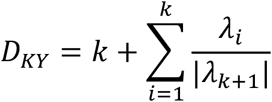

Where *k* is the largest integer where the sum of the *k* largest exponents are >= 0.

### SINDy library optimization

While Θ(**X**) represents a broad set of possible terms and interactions, the vast majority of the coefficients are expected to be 0. In the SINDy method, the candidate function library must be specified a priori, with a limitation that term selection may become poorly conditioned when the candidate library becomes too large or when the redundancy is too high amongst predictors. In the original formulation of SINDy, it was found that thresholding least squares regression enables pruning of the candidate library in an iterative fashion to promote sparse model identification.^26^ This was later found to be related to utilizing L0 norm regularization for term removal.^68^ This pruning procedure is based on the value of a thresholding parameter to promote term removal and this was found to result in improved model selection relative to promoting sparsity through using the least absolute shrinkage and selection operator (LASSO) regression.

In our study, we observed that terms varied over several orders of magnitude in their coefficient amplitudes, such that constant thresholding resulted in removal of terms which, though of small magnitude, greatly improved simulated model performance. Levels of thresholding that preserved these terms thus also resulted in other classes of candidate terms receiving no pruning during the application of the iterated thresholding least squares procedure. While normalizing these terms can mitigate this problem, there is little basis for identifying how these thresholds would then be set between different variables. We thus sought to provide a basis for expanding and removing terms from the candidate function library to identify subsets of the library containing terms which are beneficial for model accuracy. To promote parsimony, we sought a nested level of complexity where at each step, increasingly complex subsets of Θ(**X**) were included. Library expansions which did not contain accuracy increasing terms (defined as reducing error by >0.01) were removed. Accuracy for library selection was measured at the level of autonomous only function libraries simply as *AC*_*error*_, at the level of between-band within-region functions as *AC*_*error*_ + *Lag*_*error* (*between*−*band*)_ + *Coh*_*error* (*betwen*−*band*)_, and at the level of between-band and between-region as *AC*_*error*_ + *Lag*_*error* (*between*−*band*)_ + *Coh*_*error* (*betwen*−*band*)_ + Lag_*error*(*between*−*region*)_ + *Coh*_*error*(*between*−*region*)_.

Here we briefly describe the ordering of candidate libraries with formal definitions below. We define autonomous terms as including only products consisting of the real and imaginary components of the target signal without including between-band or between-region terms in the library, between-band terms defined as those which include terms of the same region but of other frequency bands, and finally the highest complexity being terms which include between-region terms which are then ordered by complexity, with within-band terms being less complex than between-band terms. Note, these broad classes of libraries were further subdivided during library expansion with increasing complexity of increasing product order containing terms from a single signal followed by product orders consisting of products of two signals (mixed products). The candidate library performance for all library additions is shown in Fig. S4B and the definitions of each addition, their order, number of terms selected/total terms possible to be selected are provided in Table S1. Additionally, models identified with simpler libraries may also contain terms which approximate the roles of more accurate but more complex functions arising in later library expansions, which after library expansion then renders the simpler function obsolete or detrimental to model performance. This suggested that iterative pruning of these functions may also serve to constrain the candidate library size during expansion and to directly compare model performance between competing predictors. Thus, at each stage upon selection of terms from an expanded library, each identified class of previously selected terms were tested for removal, keeping only terms which degraded model performance by >0.01 with removal.

### SINDy library nomenclature

We begin by outlining a notation with which to describe classes of candidate function libraries, which are combined as outlined below to expand the candidate function library in levels of increasing complexity. Each library is defined by the subset of columns (candidate functions) of *Θ*(**X**) that it admits. A general candidate function is a monomial 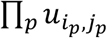 of degree 1–4, where each factor 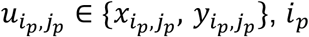 indexes region and *j*_*p*_ indexes band. Throughout, (*R, B*) denotes the region R and band B of the *target* signal, i.e., the pair for which the regression 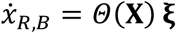 (and likewise for 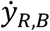) is being solved, and (*i, j*), (*k, l*) denote the region and band indices of the candidate factors.

Band library subset *Θ*(**X**_*B*_) candidate functions all of whose factors come from band *B*,

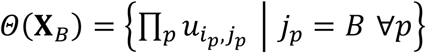

**Region library subset** *Θ*(**X**_*R*_): candidate functions all of whose factors come from region

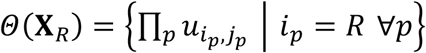

**Linear term library subset** *Θ*(**X**_Lin_): candidate functions of degree 1 (the columns of **X**^(1)^),

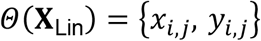

**Quadratic term library subset** *Θ*(**X**_Quad_): candidate functions of degree 2 (the columns of **X**^(2)^),

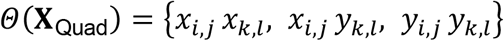

**Cubic term library subset** *Θ*(**X**_Cub_): candidate functions of degree 3 (the columns of **X**^(3)^), e.g., 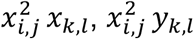, etc.

**Quartic term library subset** *Θ*(**X**_Quart_): candidate functions of degree 4 (the columns of **X**^(4)^), e.g., 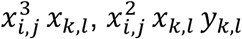, etc.

**Autonomous function library subset** *Θ*(**X**_Aut_): candidate functions all of whose factors come from the target signal itself,

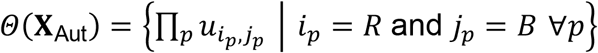

**Cross-band Intra-region function library subset** *Θ*(**X**_CrossBIntraR_): candidate functions whose factors come from the target region but products from a single band other than the target band,

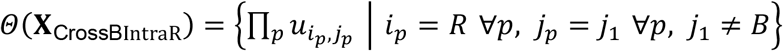

**Mixed-product cross-band intra-region library subset** *Θ*(**X**_MixedProdCrossBIntraR_): products within the target region combining at least one factor from the target band with at least one factor from a different band,

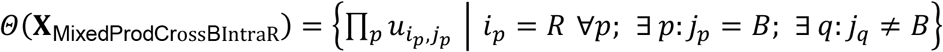

**Mixed-product cross-region library subset** *Θ*(**X**_MixedProdCrossR_): products combining factors from at least two different regions,

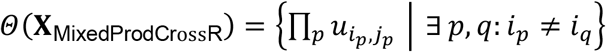

**Cross-regional intra-band library subset** *Θ*(**X**_CrossRIntraB_): candidate functions whose factors come from regions other than the target region but from the target band,

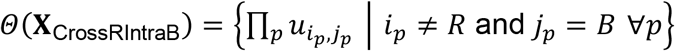

**Cross-regional cross-band library subset** *Θ*(**X**_CrossRCrossB_): candidate functions whose factors come from regions other than the target region and from bands other than the target band,

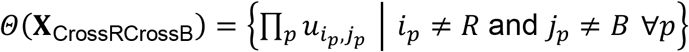

**Band-pair restriction subset** *Θ*(**X**_*k*/*l*_): mixed products whose factors come from the specific pair of bands {*k, l*}, *k* ≠ *l*,

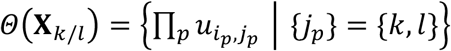

**Region-pair restriction subset** 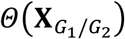: analogously, mixed products whose factors come from the specific pair of region groupings {*G*_1_, *G*_2_} (e.g., Mot/Som or Left/Right, where Transcallosal Mot = {1,2}, Transcallosal Som = {3,4}, Mot/Som Left = {1,3}, and Mot/Som Right = {2,4}.

With these sets we can then express various combinations using the following notation: *A* ∪ *B* represents the candidate functions in *A*, in *B*, or in both; *A* ∩ *B* represents the candidate functions in both *A* and *B*; *A*\*B* represents the candidate functions in *A* but not in *B*; and *A* indicates the candidate functions of *Θ*(**X**) not in *A*.

### Iterative candidate library expansion and pruning procedure

Next we performed an iterative expansion and pruning procedure where subsets of the above **X** matrices were included in the candidate library as described below where the restricted library is referred to as *Θ*(**X**′). Upon addition of a subset of functions **A** to the current *Θ*(**X**^′^) matrix, (*Θ*(**X**^′^) ∪ *Θ*(**A**)), a regression was performed and the resultant model was used in simulation and performance was calculated as above. An iterative pruning procedure was then applied to **A** to identify a subset (*Θ*(**A**_Subset_)) which maintained or improved the performance relative to *Θ*(**X**^′^) ∪ *Θ*(**A**). If after pruning, model performance was improved relative to *Θ*(**X**^′^), *Θ*(**X**′) was redefined as *Θ*(**X**^′^) ∪ *Θ*(**A**_Subset_). The pruning procedure for **A** started with product order (**X**_Lin_, **X**_Quad_, **X**_Cub_, and **X**_Quart_), then frequency band of the source of term **X**_R,B_ where B=1,2,or 3 and then region source **X**_R,B_ R=MotL, MotR, SomL, SomR, and then finally the frequency band and region of the target. Additionally, at the level of each expansion, each addition was tested for amplitude normalized variations of the functions between 0-4^th^ order and the best performing normalization was selected. We performed iterative expansions in the order shown in Table S1.

**Supp. Table 1.**
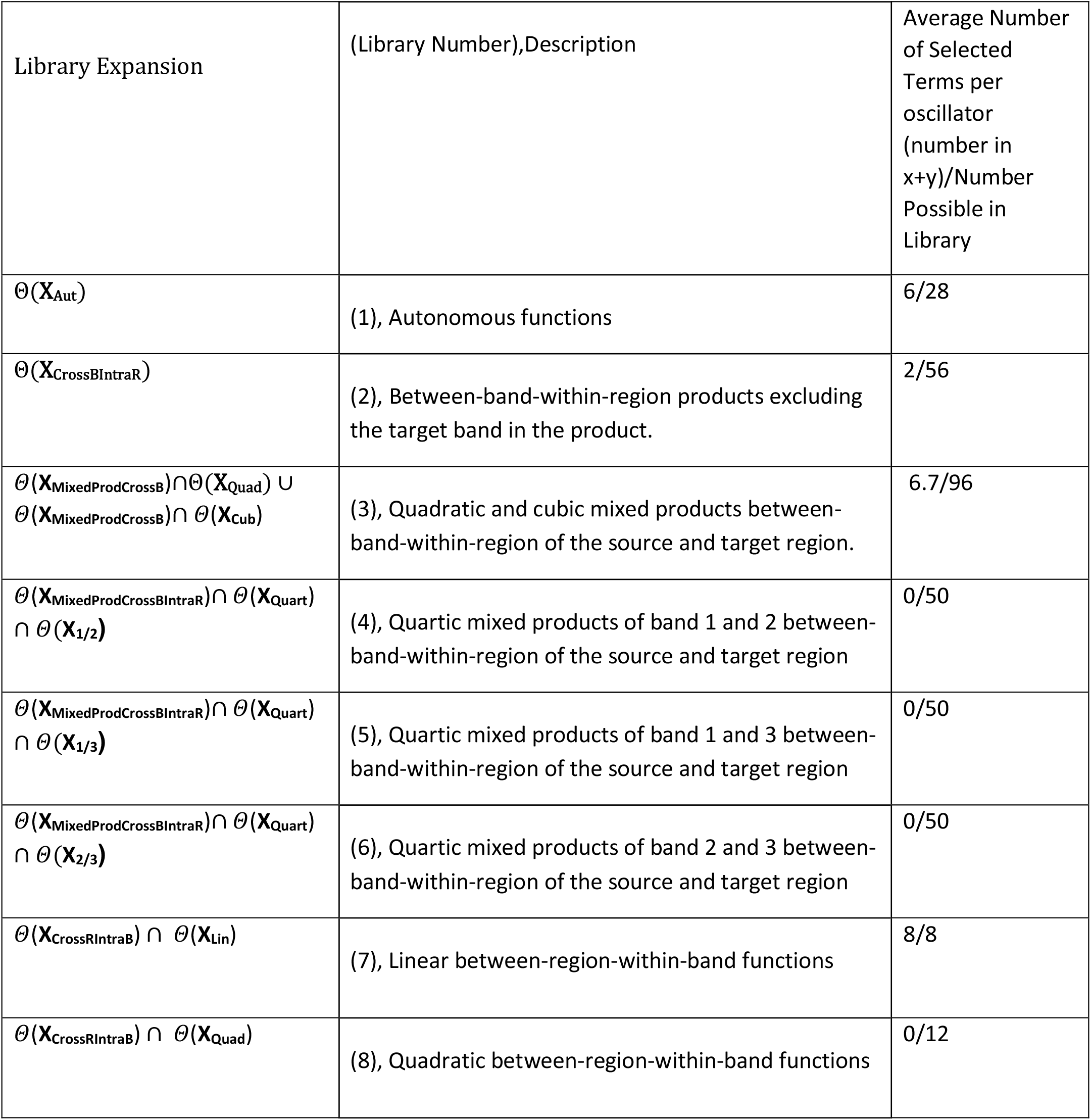
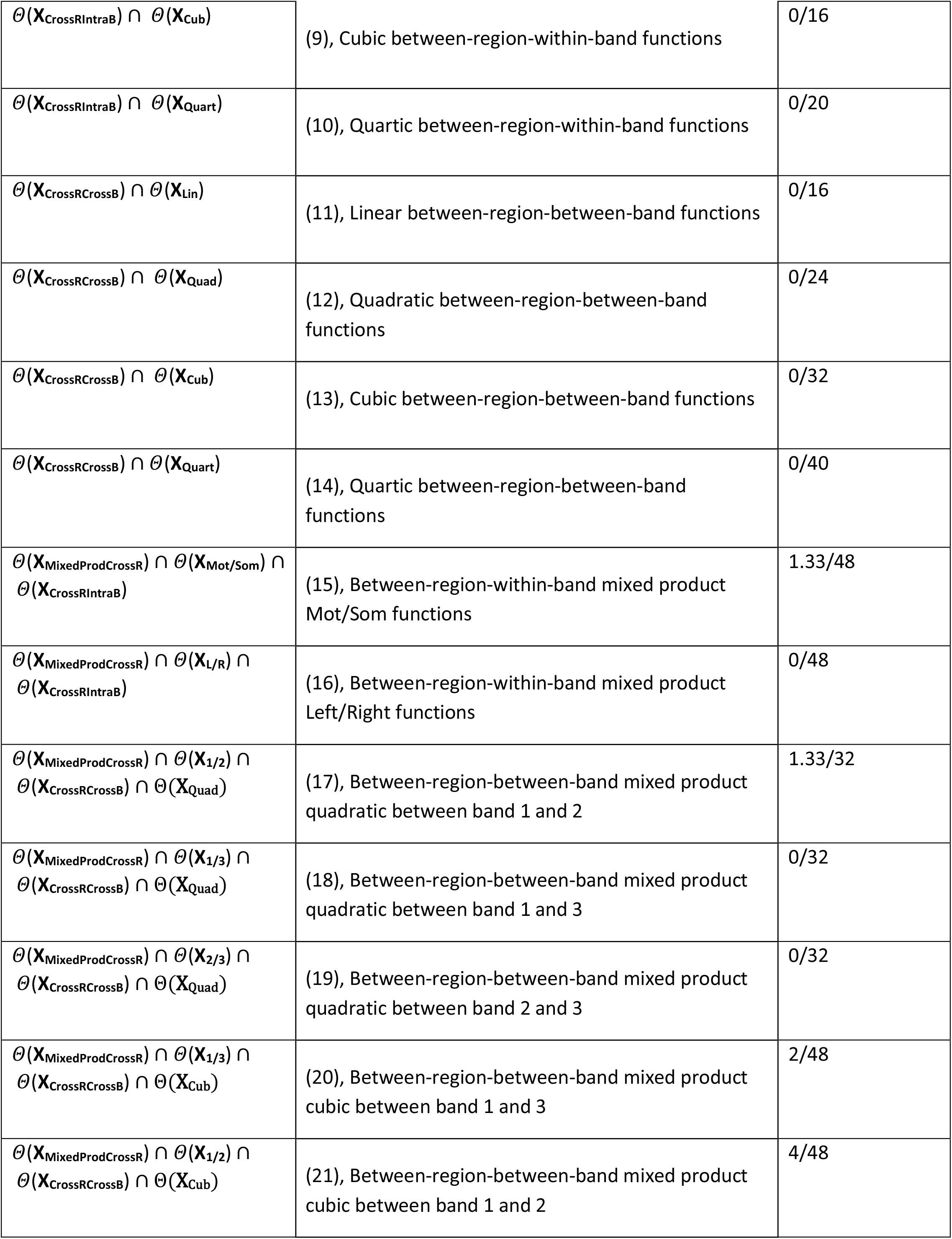
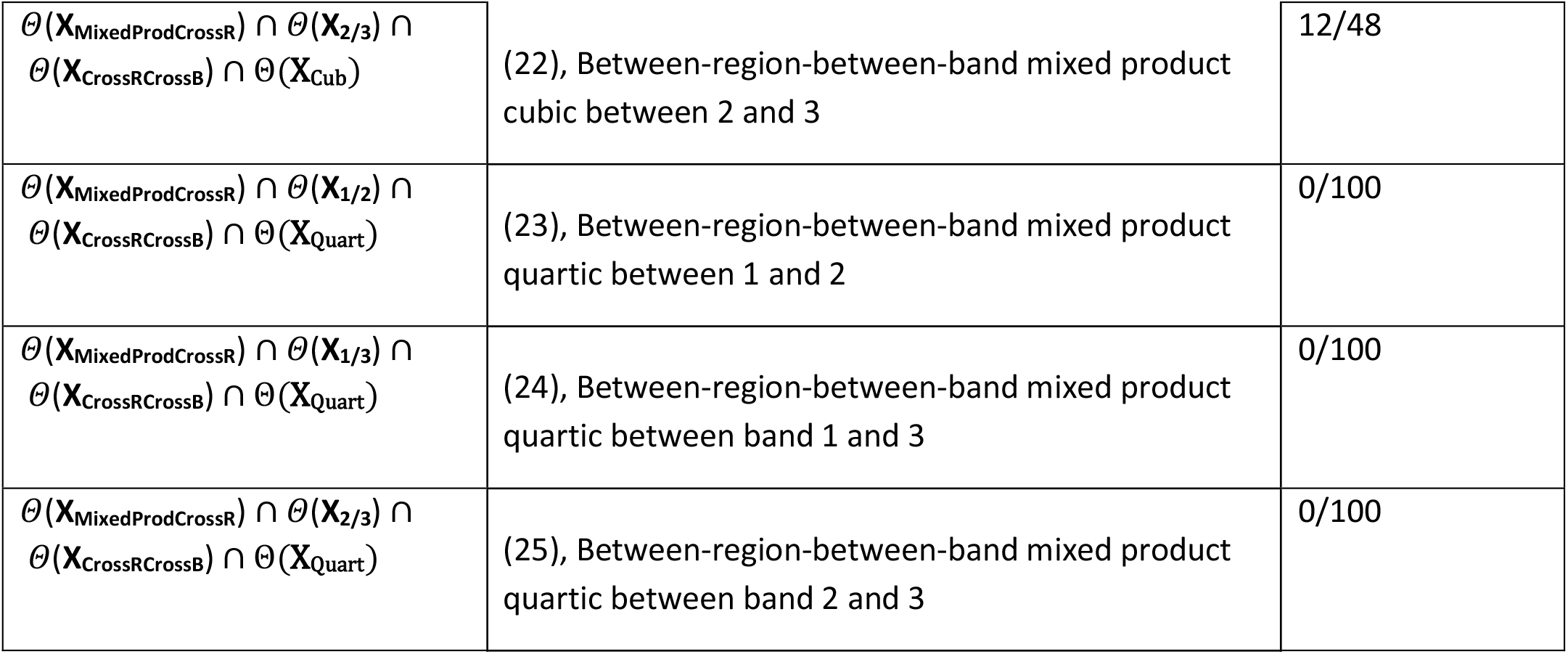
SINDy library expansion order.

| Library Expansion | (Library Number),Description | Average Number of Selected Terms per oscillator (number in x+y)/Number Possible in Library |
| --- | --- | --- |
| $\theta(\mathbf{X}_{\text{Aut}})$ | (1), Autonomous functions | 6/28 |
| $\theta(\mathbf{X}_{\text{CrossBintraR}})$ | (2), Between-band-within-region products excluding the target band in the product. | 2/56 |
| $\theta(\mathbf{X}_{\text{MixedProdCrossB}}) \cap \theta(\mathbf{X}_{\text{Quad}}) \cup \theta(\mathbf{X}_{\text{MixedProdCrossB}}) \cap \theta(\mathbf{X}_{\text{Cub}})$ | (3), Quadratic and cubic mixed products between-band-within-region of the source and target region. | 6.7/96 |
| $\theta(\mathbf{X}_{\text{MixedProdCrossBintraR}}) \cap \theta(\mathbf{X}_{\text{Quart}}) \cap \theta(\mathbf{X}_{1/2})$ | (4), Quartic mixed products of band 1 and 2 between-band-within-region of the source and target region | 0/50 |
| $\theta(\mathbf{X}_{\text{MixedProdCrossBintraR}}) \cap \theta(\mathbf{X}_{\text{Quart}}) \cap \theta(\mathbf{X}_{1/3})$ | (5), Quartic mixed products of band 1 and 3 between-band-within-region of the source and target region | 0/50 |
| $\theta(\mathbf{X}_{\text{MixedProdCrossBintraR}}) \cap \theta(\mathbf{X}_{\text{Quart}}) \cap \theta(\mathbf{X}_{2/3})$ | (6), Quartic mixed products of band 2 and 3 between-band-within-region of the source and target region | 0/50 |
| $\theta(\mathbf{X}_{\text{CrossRIntraB}}) \cap \theta(\mathbf{X}_{\text{Lin}})$ | (7), Linear between-region-within-band functions | 8/8 |
| $\theta(\mathbf{X}_{\text{CrossRIntraB}}) \cap \theta(\mathbf{X}_{\text{Quad}})$ | (8), Quadratic between-region-within-band functions | 0/12 |
| $\theta(\mathbf{X}_{\text{CrossRIntraB}}) \cap \theta(\mathbf{X}_{\text{Cub}})$ | (9), Cubic between-region-within-band functions | 0/16 |
| $\theta(\mathbf{X}_{\text{CrossRIntraB}}) \cap \theta(\mathbf{X}_{\text{Quart}})$ | (10), Quartic between-region-within-band functions | 0/20 |
| $\theta(\mathbf{X}_{\text{CrossRCrossB}}) \cap \theta(\mathbf{X}_{\text{Lin}})$ | (11), Linear between-region-between-band functions | 0/16 |
| $\theta(\mathbf{X}_{\text{CrossRCrossB}}) \cap \theta(\mathbf{X}_{\text{Quad}})$ | (12), Quadratic between-region-between-band functions | 0/24 |
| $\theta(\mathbf{X}_{\text{CrossRCrossB}}) \cap \theta(\mathbf{X}_{\text{Cub}})$ | (13), Cubic between-region-between-band functions | 0/32 |
| $\theta(\mathbf{X}_{\text{CrossRCrossB}}) \cap \theta(\mathbf{X}_{\text{Quart}})$ | (14), Quartic between-region-between-band functions | 0/40 |
| $\theta(\mathbf{X}_{\text{MixedProdCrossR}}) \cap \theta(\mathbf{X}_{\text{Mot/Som}}) \cap \theta(\mathbf{X}_{\text{CrossRIntraB}})$ | (15), Between-region-within-band mixed product Mot/Som functions | 1.33/48 |
| $\theta(\mathbf{X}_{\text{MixedProdCrossR}}) \cap \theta(\mathbf{X}_{\text{L/R}}) \cap \theta(\mathbf{X}_{\text{CrossRIntraB}})$ | (16), Between-region-within-band mixed product Left/Right functions | 0/48 |
| $\theta(\mathbf{X}_{\text{MixedProdCrossR}}) \cap \theta(\mathbf{X}_{1/2}) \cap \theta(\mathbf{X}_{\text{CrossRCrossB}}) \cap \theta(\mathbf{X}_{\text{Quad}})$ | (17), Between-region-between-band mixed product quadratic between band 1 and 2 | 1.33/32 |
| $\theta(\mathbf{X}_{\text{MixedProdCrossR}}) \cap \theta(\mathbf{X}_{1/3}) \cap \theta(\mathbf{X}_{\text{CrossRCrossB}}) \cap \theta(\mathbf{X}_{\text{Quad}})$ | (18), Between-region-between-band mixed product quadratic between band 1 and 3 | 0/32 |
| $\theta(\mathbf{X}_{\text{MixedProdCrossR}}) \cap \theta(\mathbf{X}_{2/3}) \cap \theta(\mathbf{X}_{\text{CrossRCrossB}}) \cap \theta(\mathbf{X}_{\text{Quad}})$ | (19), Between-region-between-band mixed product quadratic between band 2 and 3 | 0/32 |
| $\theta(\mathbf{X}_{\text{MixedProdCrossR}}) \cap \theta(\mathbf{X}_{1/3}) \cap \theta(\mathbf{X}_{\text{CrossRCrossB}}) \cap \theta(\mathbf{X}_{\text{Cub}})$ | (20), Between-region-between-band mixed product cubic between band 1 and 3 | 2/48 |
| $\theta(\mathbf{X}_{\text{MixedProdCrossR}}) \cap \theta(\mathbf{X}_{1/2}) \cap \theta(\mathbf{X}_{\text{CrossRCrossB}}) \cap \theta(\mathbf{X}_{\text{Cub}})$ | (21), Between-region-between-band mixed product cubic between band 1 and 2 | 4/48 |
| $\theta(\mathbf{X}_{\text{MixedProdCrossR}}) \cap \theta(\mathbf{X}_{2/3}) \cap \theta(\mathbf{X}_{\text{CrossRCrossB}}) \cap \theta(\mathbf{X}_{\text{Cub}})$ | (22), Between-region-between-band mixed product cubic between 2 and 3 | 12/48 |
| $\theta(\mathbf{X}_{\text{MixedProdCrossR}}) \cap \theta(\mathbf{X}_{1/2}) \cap \theta(\mathbf{X}_{\text{CrossRCrossB}}) \cap \theta(\mathbf{X}_{\text{Quart}})$ | (23), Between-region-between-band mixed product quartic between 1 and 2 | 0/100 |
| $\theta(\mathbf{X}_{\text{MixedProdCrossR}}) \cap \theta(\mathbf{X}_{1/3}) \cap \theta(\mathbf{X}_{\text{CrossRCrossB}}) \cap \theta(\mathbf{X}_{\text{Quart}})$ | (24), Between-region-between-band mixed product quartic between band 1 and 3 | 0/100 |
| $\theta(\mathbf{X}_{\text{MixedProdCrossR}}) \cap \theta(\mathbf{X}_{2/3}) \cap \theta(\mathbf{X}_{\text{CrossRCrossB}}) \cap \theta(\mathbf{X}_{\text{Quart}})$ | (25), Between-region-between-band mixed product quartic between band 2 and 3 | 0/100 |

### SINDy: Autonomous library

Below we show in detail an example of several steps of the above procedure. As an initial library, we first included only autonomous terms, *Θ*(**X**′) = *Θ*(**X**_Aut_) and performed the iterative pruning procedure.

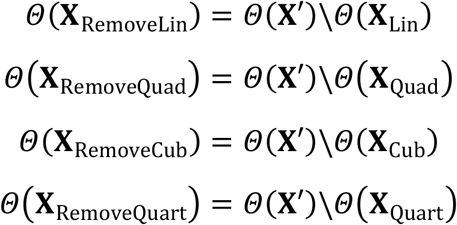

Of these subsets, *Θ*(**X**_RemoveLin_) and *Θ*(**X**_RemoveCub_) degraded model performance relative to *Θ*(**X**′), while *Θ*(**X**_RemoveQuad_) and *Θ*(**X**_RemoveQuart_) did not, so *Θ*(**X**_Combined_) was defined as

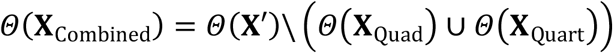

We then iteratively removed subsets of the remaining terms (*Θ*(**X**_Lin_) and *Θ*(**X**_Cub_)) by band:

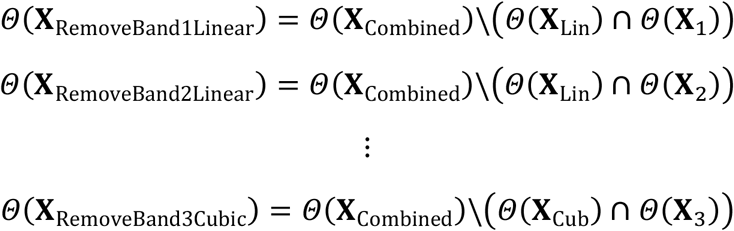

This resulted in the preservation of all sub-band terms for these product orders, leading to a redefinition of

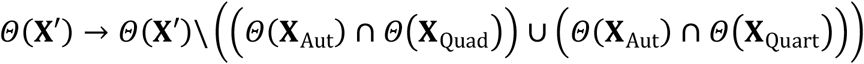

Each x and y each had 14 autonomous terms possible for inclusion in the model and 6 were selected for each x and y.

We noted at this stage that there are regularities in the magnitudes of coefficients which suggest they may be combined to form common coefficients. Below we show this procedure for the selected linear and cubic terms in the model at this step. The selected terms followed the following pattern:

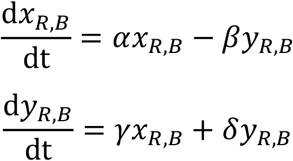

Where α, β, γ, and δ are the corresponding coefficients in ξ_k_. Coefficients α and δ were found to be near identical and β and γ were found to be near identical. We rename these common coefficients μ (α and δ) and ω (β and γ) due to their dynamical roles in intrinsic growth (μ) and intrinsic oscillation frequency (ω):

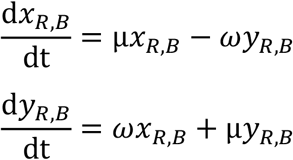

This pattern of coefficients allows us to combine terms into an equation in the complex variable z:

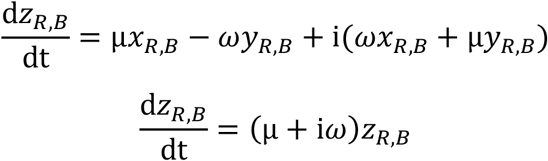

We then changed coordinates from *z* to amplitude A and phase θ:

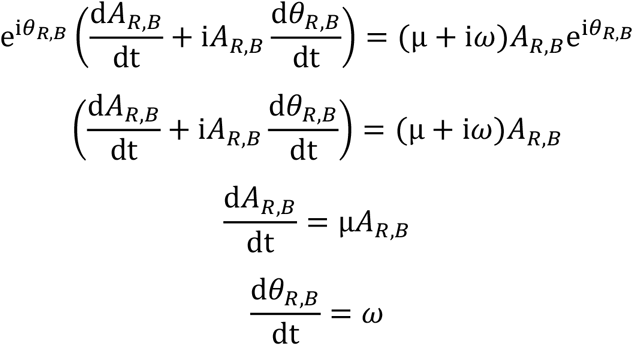

**Supp. Table 2.** Linear autonomous coefficients in the model. Algebraic simplification constructs intrinsic frequency (ωi)z. Bold indicates the simplified function in terms of z.

| Region | $dx_{(R,B)}/dt = \omega y_{(R,B)}$ | $dy_{(R,B)}/dt = \omega x_{(R,B)}$ | **Simplified Function and Remainder** | Intrinsic Frequency |
| --- | --- | --- | --- | --- |

**Supp. Table 2.** Linear autonomous coefficients in the model. Algebraic simplification constructs intrinsic frequency ( $\omega$ )z. Bold indicates the simplified function in terms of z.
| | dx(R,B) | $\gamma^* y_{R,B}$ | $\beta^* x_{R,B}$ | dz(R,B)/dt= | |
| --- | --- | --- | --- | --- | --- |
| MotL Band 1 | (1,1) | -18.33 | 18.35 | **(18.33 i)** $z_{1,1} + (0.013i)\text{Re}(z_{1,1})$ | 6.3 Hz |
| MotR Band 1 | (2,1) | -19.38 | 19.38 | **(19.38 i)** $z_{2,1} + (0.01i)\text{Re}(z_{2,1})$ | 6.7 Hz |
| SomL Band1 | (3,1) | -20.56 | 20.52 | **(20.56 i)** $z_{3,1} - (0.04i)\text{Re}(z_{3,1})$ | 7.1 Hz |
| SomR Band 1 | (4,1) | -21.75 | 21.75 | **(21.8 i)** $z_{4,1} + (0.00i)\text{Re}(z_{4,1})$ | 7.54 Hz |
| MotL Band 2 | (1,2) | -35.49 | 35.89 | **(35.49 i)** $z_{1,2} + (0.39i)\text{Re}(z_{1,2})$ | 12.3 Hz |
| MotR Band 2 | (2,2) | -37.44 | 38.08 | **(37.74 i)** $z_{2,2} + (0.63i)\text{Re}(z_{2,2})$ | 13.1 Hz |
| SomL Band 2 | (3,2) | -40.37 | 40.36 | **(40.6 i)** $z_{3,2} - (0.00i)\text{Re}(z_{3,2})$ | 14.1 Hz |
| SomR Band 2 | (4,2) | -40.83 | 40.85 | **(40.83 i)** $z_{4,2} + (0.02i)\text{Re}(z_{4,2})$ | 14.1 Hz |
| MotL Band 3 | (1,3) | -51.02 | 50.99 | **(51.02 i)** $z_{1,3} - (0.03i)\text{Re}(z_{1,3})$ | 17.7 Hz |
| MotR Band 3 | (2,3) | -54.12 | 54.10 | **(54.12 i)** $z_{2,3} - (0.02i)\text{Re}(z_{2,3})$ | 18.7 Hz |
| SomL Band 3 | (3,3) | -56.29 | 56.35 | **(56.29 i)** $z_{3,3} + (0.07i)\text{Re}(z_{3,3})$ | 19.5 Hz |
| SomR Band 3 | (4,3) | -56.45 | 56.47 | **(56.45 i)** $z_{4,3} + (0.02i)\text{Re}(z_{4,3})$ | 19.5 Hz |

Notably, while the μ terms are selected at the introduction of the library, they were later pruned after the introduction of cross regional coupling library 7 (between-region coupling within band). We show the final values of ω in the final model and the remainder after simplification. The following coefficients are divided by a normalizing scale factor of 230 to put them on a regular scale across terms. Truncation of the remainders did not affect total error (0.19 vs. 0.19).

**Supp. Table 3.**
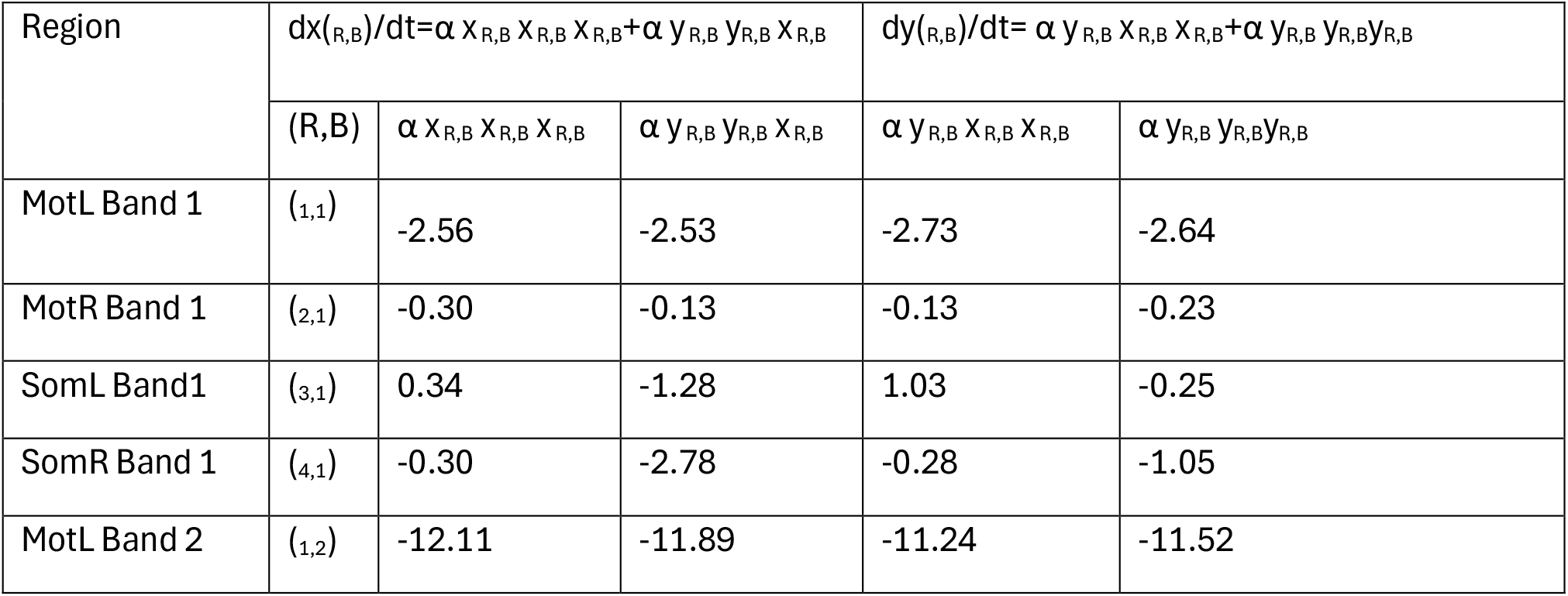
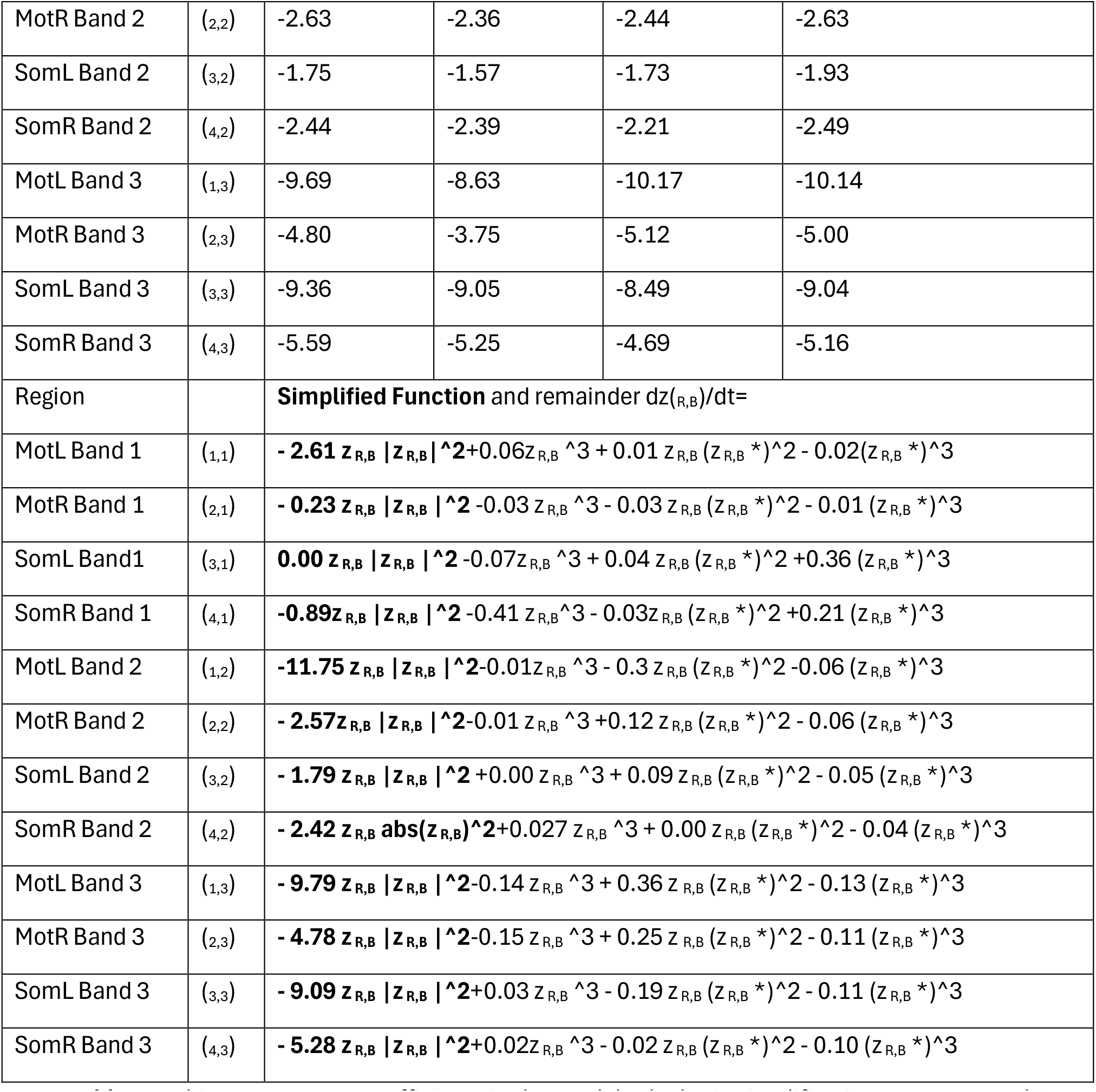
7Cubic autonomous coefficients in the model. Algebraic simplification constructs cubic limit cycle term α z _R,B_ |z _R,B_ |^2. Bold indicates the simplified function in z.

We also observed for cubic autonomous terms several common coefficient amplitude terms. Here we combine directly to α and β, named for their roles in cubic nonlinear oscillator behavior.

Cubic terms:

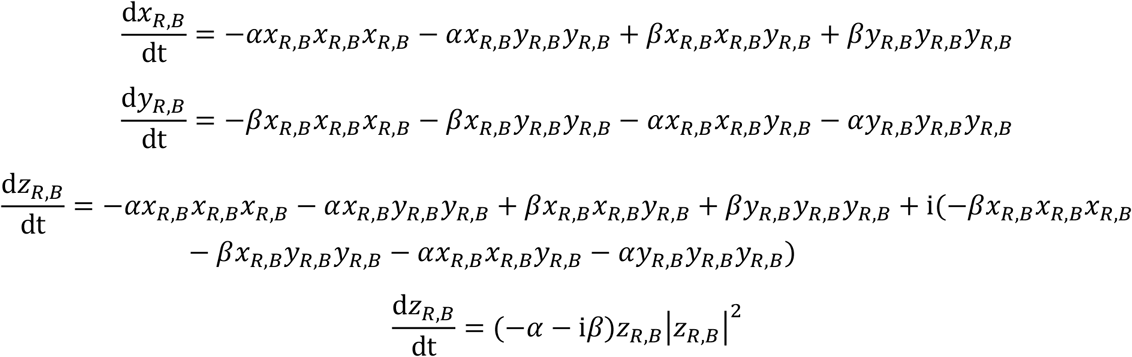

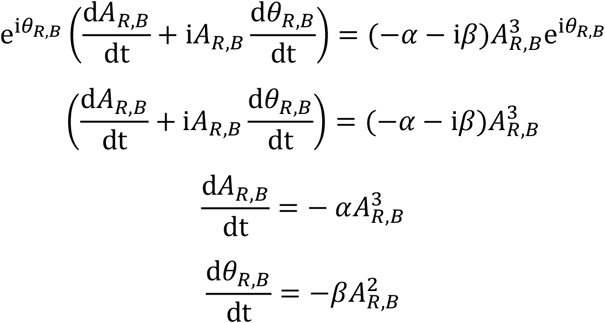

Notably at this stage, the combination of the linear and cubic functions reveals a common structure where the normal form for the Hopf bifurcation is identified with bifurcation parameter μ, limit cycle term *α*, natural oscillation frequency term *ω*, and an amplitude dependent frequency term *β*. Of the 28 possible coefficients for each oscillator (14 for each x and y) there were 12 terms selected at this stage (6 for each x and y), which when common amplitude coefficients are combined reduce to a 4-parameter model for each oscillator:

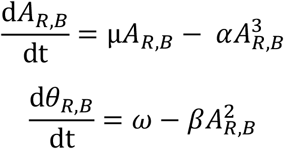

While a 4 term model was identified at this stage, μ and *β* were later culled upon addition of linear cross regional coupling and mixed products cross frequency libraries respectively reducing the number to 6 terms selected per oscillator (3 for each x and y) which reduce to the two autonomous parameters shown in Fig. 4A. Additionally, while the coefficients for α are conserved robustly up to the level of within-band cross-regional coupling, they exhibit a decrease in amplitude in the final model and exhibit higher remainders after factoring out the common terms for α. The components going into the final model are as follows. The following coefficients are divided by a normalizing scale factor of 7543457397 to put them on a regular scale across terms. Truncation of the remainders did not affect total error (0.19 vs. 0.19).

**Supp. Table 4.**
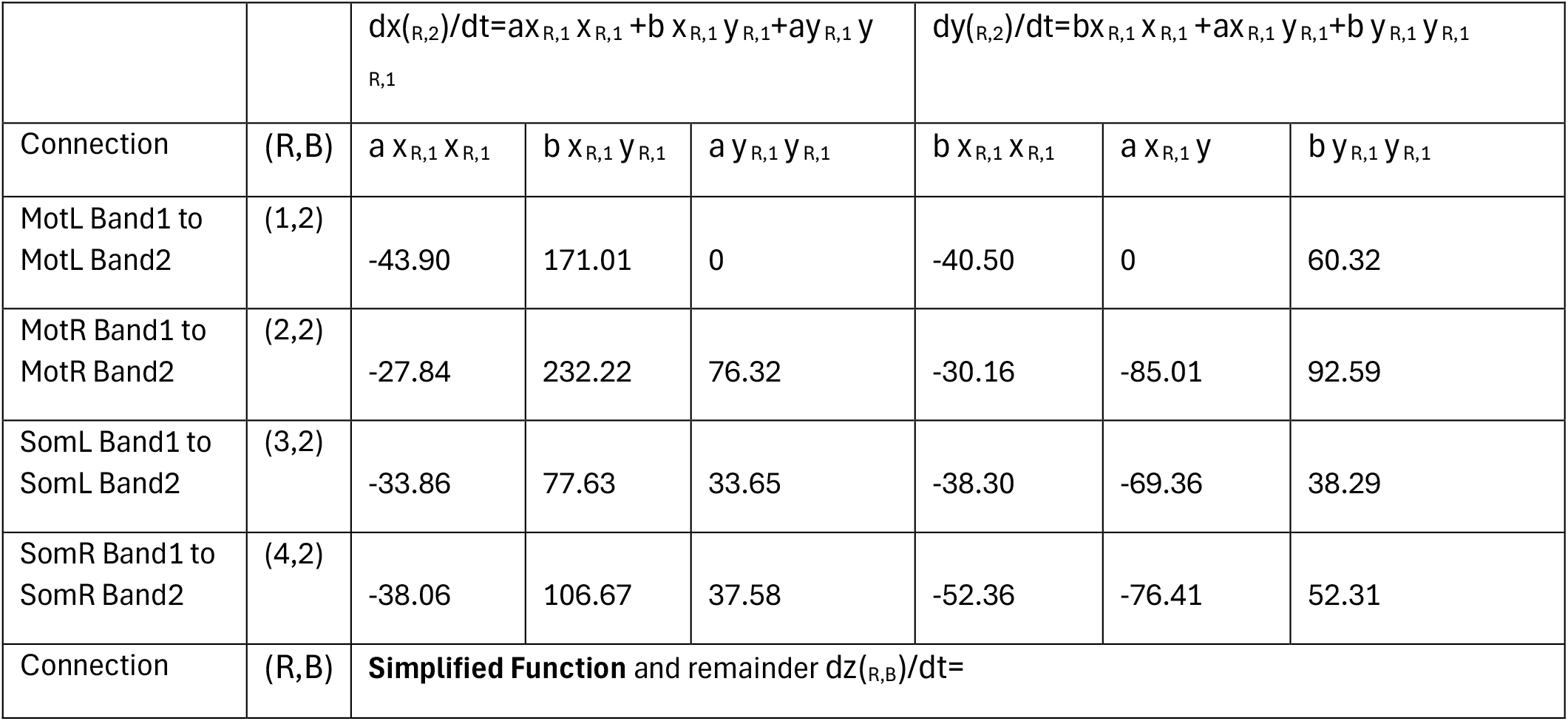
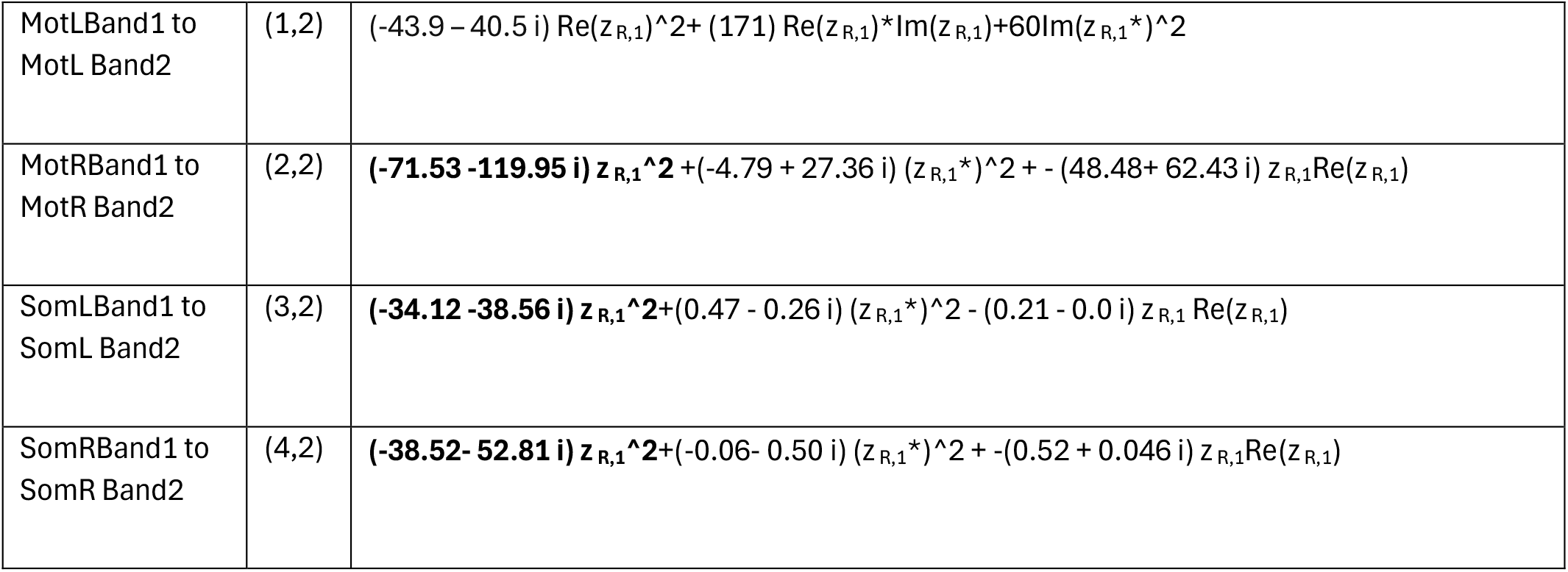
Band 1 to 2 coupling functions in the model. Bold indicates the simplified function in z.

### SINDy: Between-band coupling libraries

The next expansion was cross frequency coupling terms to the candidate function library (*Θ*(**X**^′^) ∪ *Θ*(**X**_*CrossBIntraR*_)). This expansion includes 28 functions for each x and y.

We then sought sub-regions within *Θ*(**X**_*CrossBIntraR*_) which contributed to the improved model performance of *Θ*(**X**^′^) ∪ *Θ*(**X**_*CrossBIntraR*_) relative to *Θ*(**X**^′^).

We defined subregions first by the product order within the expansion for terms which are linear, quadratic, cubic, or quartic and cross frequency:

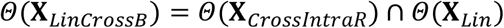

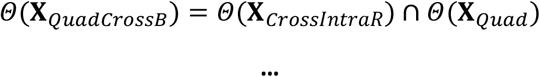

We then test the removal of each of the above subsets from *Θ*(**X**′) ∪ *Θ*(**X**_*CrossB*_):

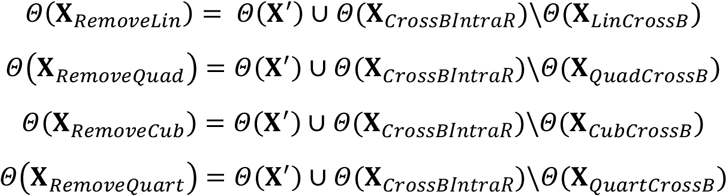

Of the subsets, *Θ*(**X**_*RemoveQuad*_) and *Θ*(**X**_*RemoveCub*_) degraded model performance relative to *Θ*(**X**^′^) while *Θ*(**X**_*RemoveLin*_) *and Θ*(**X**_*RemoveQuart*_) did not so *Θ*(**X**_*CombinedOrder*_) was defined:

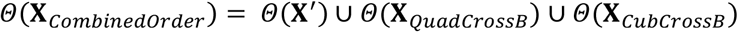

We then removed subsets of *Θ*(**X**_*QuadCrossB*_) *and Θ*(**X**_*CubCrossB*_) by band

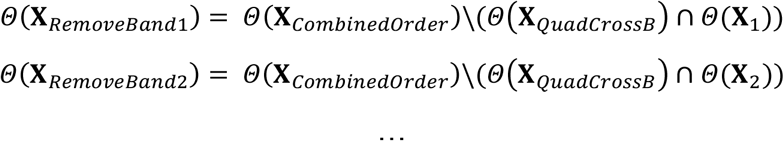

Which resulted in the sub regions of **X**_*CrossBIntraR*_ included in the candidate function being:

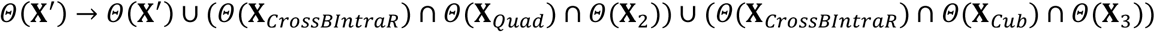

Examination of the functions built for cross frequency interactions at this step identified a common structure which is limited to quadratic cross frequency functions in band 2 terms and cubic cross frequency functions in band 3 terms while all linear and quartic functions were removed. The selected functions exhibited a high degree of regularity as observed above with the Autonomous library, resulting in common coefficients. For band 1 coupling to band 2, we identify equations of the following form:

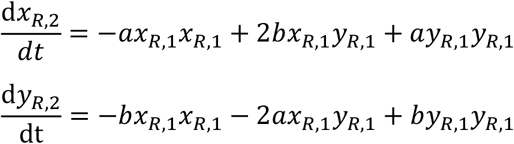

Where a and b are coefficients in ξ_k_. We then combined common terms as above and converted to polar coordinates, yielding cross frequency coupling functions which relate the product order (quadratic) to the frequency ratio between the source and target.

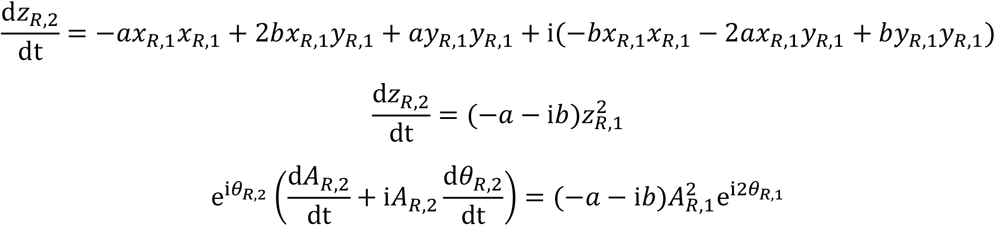

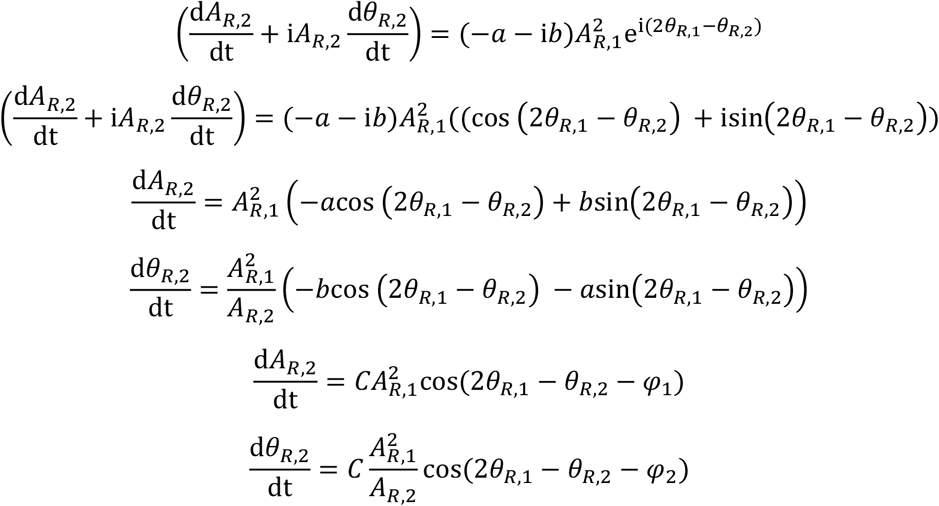

Where 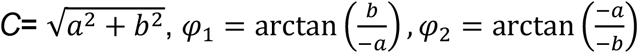

Here, coupling strength grows with *A*^*2*^_*R,1*_ for amplitude and *A*^*2*^_*R,1*_*/A*_*R,2*_ for phase coupling. This results in negligible modulation at 200 μV but grows to levels comparable to between-regional coupling at 1000 μV (Fig. S5I and J). Thus amplitude increases band 1-to-2 coupling strength, explaining the absence of these phase relationships outside seizures in experimental data (Fig. 2F, Fig. S2I).

In the final model, the Mot region deviated from the above relatively simple pattern though plotting the function on the torus gave an overall very similar form as the Som regions but with an additional distortion along diagonal trajectories (Fig. S5IJ). The following coefficients are divided by a normalizing scale factor of 19763881.45 to put them on a regular scale across terms.

We next expanded to include mixed products consisting of mixed quadratic functions and mixed cubic functions. Several functions were selected at this stage, many of which were pruned in the later models when cross regional functions were included in the library. The functions that remain in the final model at this stage are those shown graphically in Fig. 4B, middle, see accompanying MATLAB code for model expansion and sparsification steps. These functions contributed relatively little to overall model performance (Fig. S4F). See accompanying model spreadsheet for the exact coefficients.

### SINDy: Between-region coupling libraries

We next added linear cross regional coupling within band functions to the candidate function library. We observe that two different normalized variants are selected. Only the MotL/R regions selected the un-normalized amplitude variants.

We identify functions of the following form:

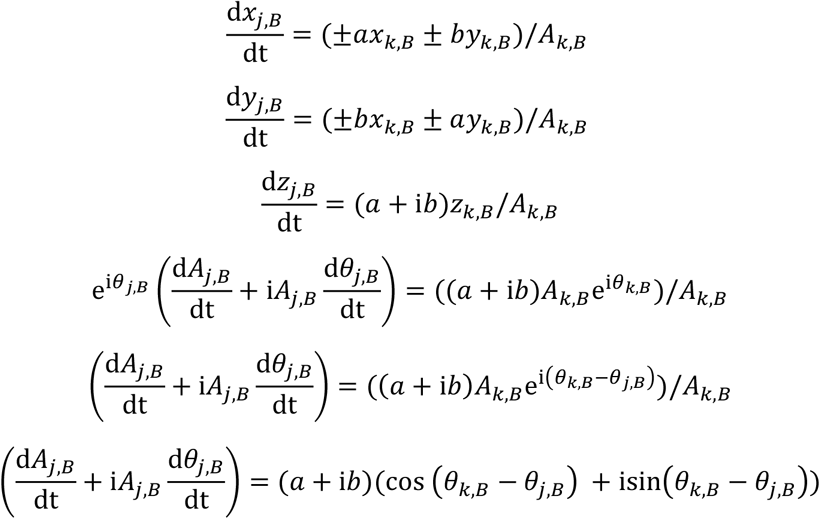

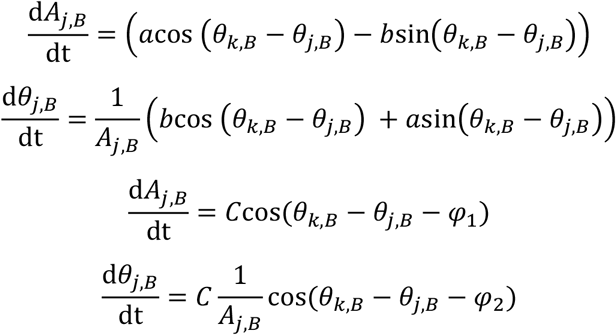

Where 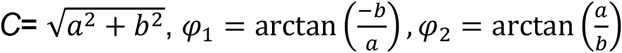

The steps for the non-amplitude normalized functions are the same as above but without a divisor of A_k,B_ resulting in coupling equations of the form:

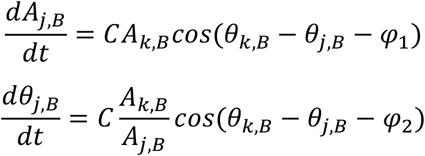

We note that the above functions exhibit phase dependent and amplitude dependent interaction terms. Notably the phase difference coupling portion of the terms is similar to those used in, ^38^ and by extension the Kuramoto model of phase coupled oscillators. Below we list all of the above coupling functions. We observe that while the majority of each function is dominated by those of the form described above with a proportionally small remainder of the form (E+Fi)*Re(z). By truncating the coefficients corresponding to the (E+Fi)*Re(z) remainder components and keeping only the (A+Bi)*z components we found they did not impact on model performance (0.19 vs 0.19).The following coefficients are divided by a normalizing scale factor of 230 to put them on a regular scale across terms.

**Supp. Table 5.**
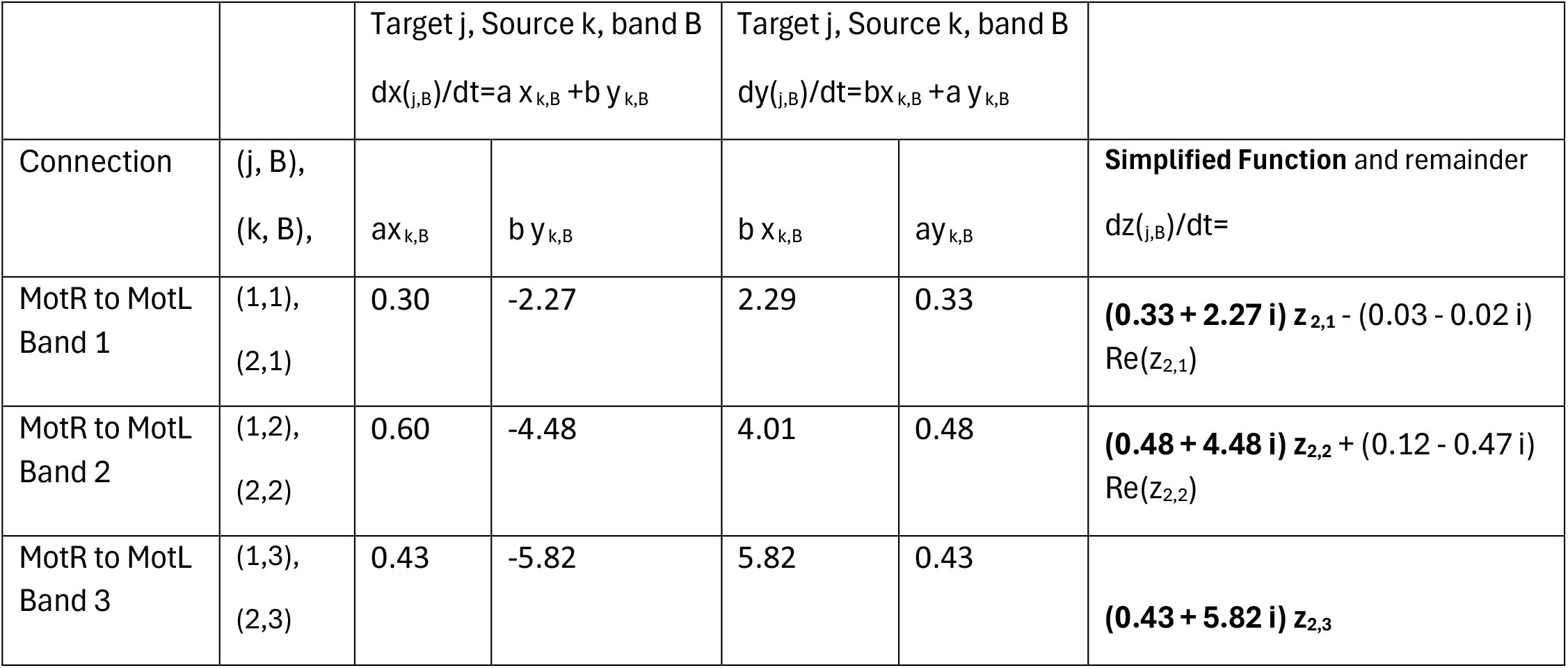
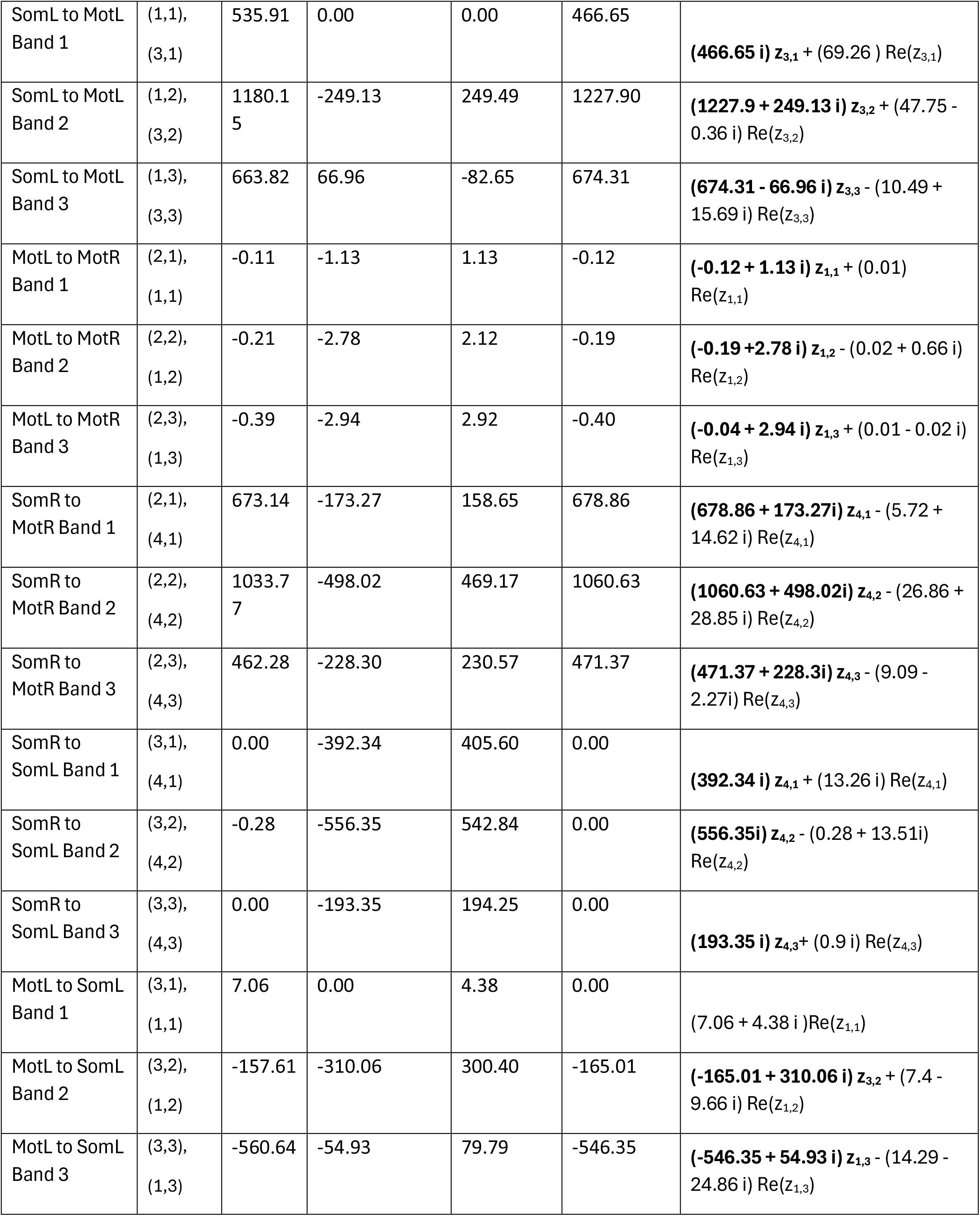
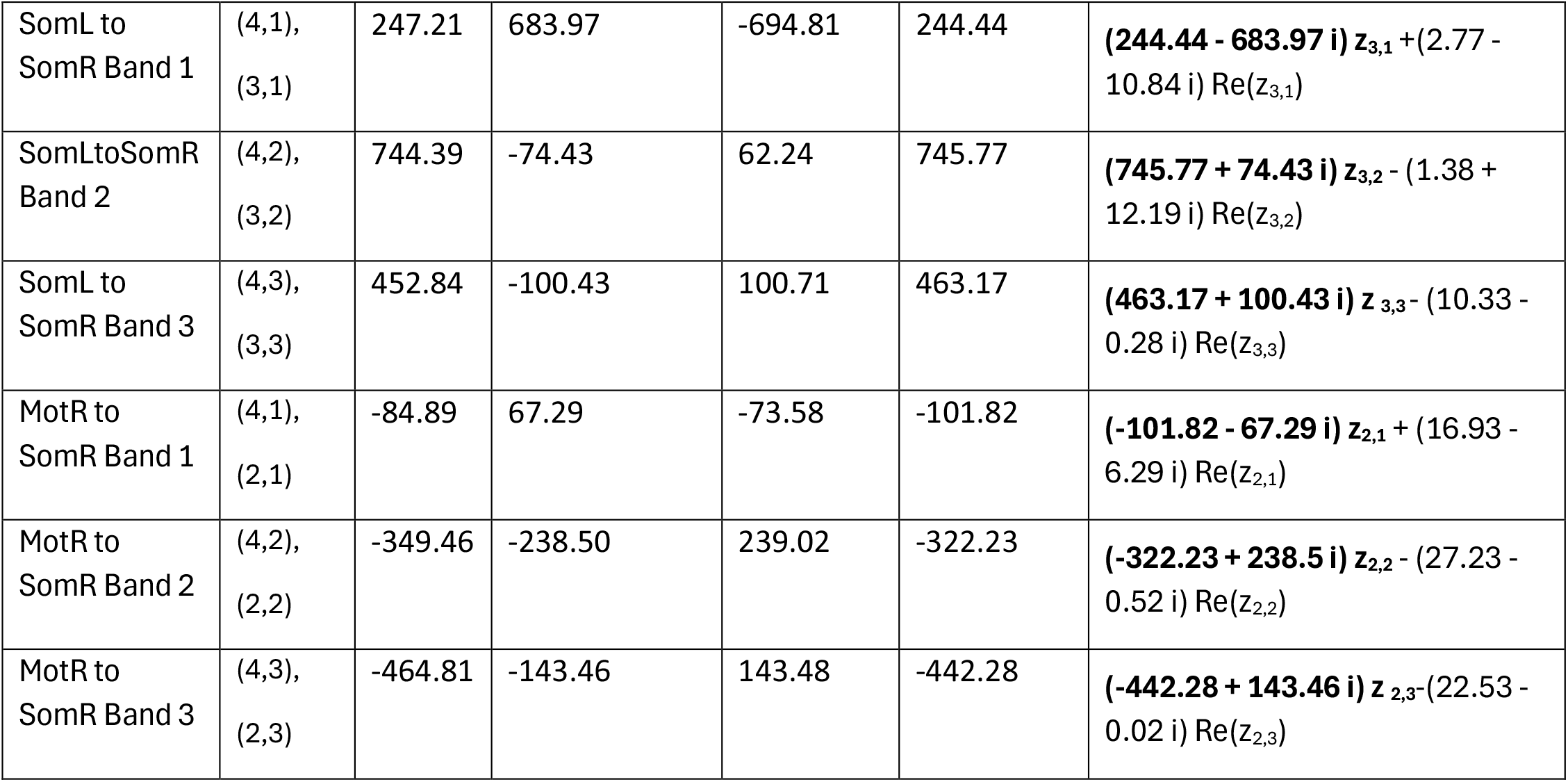
Cross regional coupling functions in the model from cross frequency library 1. Bold indicates simplified function in z.

| | | Target j, Source k, band B<br>$dx_{(j,B)}/dt = a x_{k,B} + b y_{k,B}$ | | Target j, Source k, band B<br>$dy_{(j,B)}/dt = b x_{k,B} + a y_{k,B}$ | | |
| --- | --- | --- | --- | --- | --- | --- |
| Connection | (j, B),<br>(k, B), | $a x_{k,B}$ | $b y_{k,B}$ | $b x_{k,B}$ | $a y_{k,B}$ | **Simplified Function** and remainder<br>$dz_{(j,B)}/dt =$ |
| MotR to MotL<br>Band 1 | (1,1),<br>(2,1) | 0.30 | -2.27 | 2.29 | 0.33 | **(0.33 + 2.27 i) $z_{2,1}$** - (0.03 - 0.02 i) $\text{Re}(z_{2,1})$ |
| MotR to MotL<br>Band 2 | (1,2),<br>(2,2) | 0.60 | -4.48 | 4.01 | 0.48 | **(0.48 + 4.48 i) $z_{2,2}$** + (0.12 - 0.47 i) $\text{Re}(z_{2,2})$ |
| MotR to MotL<br>Band 3 | (1,3),<br>(2,3) | 0.43 | -5.82 | 5.82 | 0.43 | **(0.43 + 5.82 i) $z_{2,3}$** |
| SomL to MotL<br>Band 1 | (1,1),<br>(3,1) | 535.91 | 0.00 | 0.00 | 466.65 | **(466.65 i) $z_{3,1}$ + (69.26 ) Re($z_{3,1}$)** |
| SomL to MotL<br>Band 2 | (1,2),<br>(3,2) | 1180.1<br>5 | -249.13 | 249.49 | 1227.90 | **(1227.9 + 249.13 i) $z_{3,2}$ + (47.75 - 0.36 i) Re($z_{3,2}$)** |
| SomL to MotL<br>Band 3 | (1,3),<br>(3,3) | 663.82 | 66.96 | -82.65 | 674.31 | **(674.31 - 66.96 i) $z_{3,3}$ - (10.49 + 15.69 i) Re($z_{3,3}$)** |
| MotL to MotR<br>Band 1 | (2,1),<br>(1,1) | -0.11 | -1.13 | 1.13 | -0.12 | **(-0.12 + 1.13 i) $z_{1,1}$ + (0.01) Re($z_{1,1}$)** |
| MotL to MotR<br>Band 2 | (2,2),<br>(1,2) | -0.21 | -2.78 | 2.12 | -0.19 | **(-0.19 + 2.78 i) $z_{1,2}$ - (0.02 + 0.66 i) Re($z_{1,2}$)** |
| MotL to MotR<br>Band 3 | (2,3),<br>(1,3) | -0.39 | -2.94 | 2.92 | -0.40 | **(-0.04 + 2.94 i) $z_{1,3}$ + (0.01 - 0.02 i) Re($z_{1,3}$)** |
| SomR to<br>MotR Band 1 | (2,1),<br>(4,1) | 673.14 | -173.27 | 158.65 | 678.86 | **(678.86 + 173.27i) $z_{4,1}$ - (5.72 + 14.62 i) Re($z_{4,1}$)** |
| SomR to<br>MotR Band 2 | (2,2),<br>(4,2) | 1033.7<br>7 | -498.02 | 469.17 | 1060.63 | **(1060.63 + 498.02i) $z_{4,2}$ - (26.86 + 28.85 i) Re($z_{4,2}$)** |
| SomR to<br>MotR Band 3 | (2,3),<br>(4,3) | 462.28 | -228.30 | 230.57 | 471.37 | **(471.37 + 228.3i) $z_{4,3}$ - (9.09 - 2.27i) Re($z_{4,3}$)** |
| SomR to<br>SomL Band 1 | (3,1),<br>(4,1) | 0.00 | -392.34 | 405.60 | 0.00 | **(392.34 i) $z_{4,1}$ + (13.26 i) Re($z_{4,1}$)** |
| SomR to<br>SomL Band 2 | (3,2),<br>(4,2) | -0.28 | -556.35 | 542.84 | 0.00 | **(556.35i) $z_{4,2}$ - (0.28 + 13.51i) Re($z_{4,2}$)** |
| SomR to<br>SomL Band 3 | (3,3),<br>(4,3) | 0.00 | -193.35 | 194.25 | 0.00 | **(193.35 i) $z_{4,3}$ + (0.9 i) Re($z_{4,3}$)** |
| MotL to SomL<br>Band 1 | (3,1),<br>(1,1) | 7.06 | 0.00 | 4.38 | 0.00 | **(7.06 + 4.38 i )Re($z_{1,1}$)** |
| MotL to SomL<br>Band 2 | (3,2),<br>(1,2) | -157.61 | -310.06 | 300.40 | -165.01 | **(-165.01 + 310.06 i) $z_{3,2}$ + (7.4 - 9.66 i) Re($z_{1,2}$)** |
| MotL to SomL<br>Band 3 | (3,3),<br>(1,3) | -560.64 | -54.93 | 79.79 | -546.35 | **(-546.35 + 54.93 i) $z_{1,3}$ - (14.29 - 24.86 i) Re($z_{1,3}$)** |
| SomL to SomR Band 1 | (4,1),<br>(3,1) | 247.21 | 683.97 | -694.81 | 244.44 | **(244.44 - 683.97 i) $z_{3,1}$** + (2.77 - 10.84 i) $\text{Re}(z_{3,1})$ |
| SomLtoSomR Band 2 | (4,2),<br>(3,2) | 744.39 | -74.43 | 62.24 | 745.77 | **(745.77 + 74.43 i) $z_{3,2}$** - (1.38 + 12.19 i) $\text{Re}(z_{3,2})$ |
| SomL to SomR Band 3 | (4,3),<br>(3,3) | 452.84 | -100.43 | 100.71 | 463.17 | **(463.17 + 100.43 i) $z_{3,3}$** - (10.33 - 0.28 i) $\text{Re}(z_{3,3})$ |
| MotR to SomR Band 1 | (4,1),<br>(2,1) | -84.89 | 67.29 | -73.58 | -101.82 | **(-101.82 - 67.29 i) $z_{2,1}$** + (16.93 - 6.29 i) $\text{Re}(z_{2,1})$ |
| MotR to SomR Band 2 | (4,2),<br>(2,2) | -349.46 | -238.50 | 239.02 | -322.23 | **(-322.23 + 238.5 i) $z_{2,2}$** - (27.23 - 0.52 i) $\text{Re}(z_{2,2})$ |
| MotR to SomR Band 3 | (4,3),<br>(2,3) | -464.81 | -143.46 | 143.48 | -442.28 | **(-442.28 + 143.46 i) $z_{2,3}$** - (22.53 - 0.02 i) $\text{Re}(z_{2,3})$ |

The remainder of selected functions are shown graphically in Fig. S4E. We note that the remaining functions contribute little to overall model performance or modulation of seizures (Fig. S4F and G). All coefficients with library selection steps are available in the accompanying MATLAB code available at https://github.com/hulljm/EmergentSeizures.

## Citations

1. Meeren H, van Luijtelaar G, da Silva FL, Coenen A. Evolving concepts on the pathophysiology of absence seizures: the cortical focus theory. Archives of neurology. 2005;62(3):371–6.

2. Polack P-O, Guillemain I, Hu E, Deransart C, Depaulis A, Charpier S. Deep layer somatosensory cortical neurons initiate spike-and-wave discharges in a genetic model of absence seizures. Journal of Neuroscience. 2007;27(24):6590–9.

3. McCafferty C, Gruenbaum BF, Tung R, Li J-J, Zheng X, Salvino P, Vincent P, Kratochvil Z, Ryu JH, Khalaf A. Decreased but diverse activity of cortical and thalamic neurons in consciousness-impairing rodent absence seizures. Nature Communications. 2023;14(1):117.

4. McCafferty C, David F, Venzi M, Lőrincz ML, Delicata F, Atherton Z, Recchia G, Orban G, Lambert RC, Di Giovanni G. Cortical drive and thalamic feed-forward inhibition control thalamic output synchrony during absence seizures. Nature neuroscience. 2018;21(5):744–56.

5. Dulko E, Kilianski S, Carns AG, Rajesh A, Lesmana I, Pikus M, Beenhakker M. Thalamic hubs as early sources of global neuronal synchrony in absence epilepsy. bioRxiv. 2025:2025.12.19.695246.

6. Khan W, Chopra S, Zheng X, Liu S, Paszkowski P, Valcarce-Aspegren M, Sieu L-A, Mcgill S, Mccafferty C, Blumenfeld H. Neuronal rhythmicity and cortical arousal in a mouse model of absence epilepsy. Experimental neurology. 2024;381:114925.

7. Blumenfeld H, McCormick DA. Corticothalamic inputs control the pattern of activity generated in thalamocortical networks. Journal of Neuroscience. 2000;20(13):5153–62.

8. Bal T, Debay D, Destexhe A. Cortical feedback controls the frequency and synchrony of oscillations in the visual thalamus. Journal of Neuroscience. 2000;20(19):7478–88.

9. Buzsáki G. The thalamic clock: emergent network properties. Neuroscience. 1991;41(2-3):351–64.

10. Huguenard J, Prince D. Intrathalamic rhythmicity studied in vitro: nominal T-current modulation causes robust antioscillatory effects. Journal of Neuroscience. 1994;14(9):5485–502.

11. Huguenard JR, McCormick DA. Thalamic synchrony and dynamic regulation of global forebrain oscillations. Trends in neurosciences. 2007;30(7):350–6.

12. Avoli M, Gloor P. Interaction of cortex and thalamus in spike and wave discharges of feline generalized penicillin epilepsy. Experimental neurology. 1982;76(1):196–217.

13. Angelaki DE, Gu Y, DeAngelis GC. Multisensory integration: psychophysics, neurophysiology, and computation. Current opinion in neurobiology. 2009;19(4):452–8.

14. Nicolelis MA, Baccala LA, Lin RC, Chapin JK. Sensorimotor encoding by synchronous neural ensemble activity at multiple levels of the somatosensory system. Science. 1995;268(5215):1353–8.

15. Lamme VA, Super H, Spekreijse H. Feedforward, horizontal, and feedback processing in the visual cortex. Current opinion in neurobiology. 1998;8(4):529–35.

16. Strogatz SH. Nonlinear dynamics and chaos: with applications to physics, biology, chemistry, and engineering (studies in nonlinearity): Westview press; 2001.

17. Luo TZ, Kim TD, Gupta D, Bondy AG, Kopec CD, Elliott VA, DePasquale B, Brody CD. Transitions in dynamical regime and neural mode during perceptual decisions. Nature. 2025:1–11.

18. Inagaki HK, Fontolan L, Romani S, Svoboda K. Discrete attractor dynamics underlies persistent activity in the frontal cortex. Nature. 2019;566(7743):212–7.

19. Gardner RJ, Hermansen E, Pachitariu M, Burak Y, Baas NA, Dunn BA, Moser M-B, Moser EI. Toroidal topology of population activity in grid cells. Nature. 2022;602(7895):123–8.

20. Vinograd A, Nair A, Kim JH, Linderman SW, Anderson DJ. Causal evidence of a line attractor encoding an affective state. Nature. 2024;634(8035):910–8.

21. Mattia M, Pani P, Mirabella G, Costa S, Del Giudice P, Ferraina S. Heterogeneous attractor cell assemblies for motor planning in premotor cortex. Journal of Neuroscience. 2013;33(27):11155–68.

22. Deco G, Jirsa VK. Ongoing cortical activity at rest: criticality, multistability, and ghost attractors. Journal of Neuroscience. 2012;32(10):3366–75.

23. Jirsa VK, Stacey WC, Quilichini PP, Ivanov AI, Bernard C. On the nature of seizure dynamics. Brain. 2014;137(8):2210–30.

24. Makinson CD, Tanaka BS, Sorokin JM, Wong JC, Christian CA, Goldin AL, Escayg A, Huguenard JR. Regulation of thalamic and cortical network synchrony by Scn8a. Neuron. 2017;93(5):1165–79.e6.

25. Cox DR. Regression models and life-tables. Journal of the Royal Statistical Society: Series B (Methodological). 1972;34(2):187–202.

26. Brunton SL, Proctor JL, Kutz JN. Discovering governing equations from data by sparse identification of nonlinear dynamical systems. Proceedings of the national academy of sciences. 2016;113(15):3932–7.

27. Kuramoto Y. International symposium on mathematical problems in theoretical physics. Lecture notes in Physics. 1975;30:420.

28. Ohno S, Kuramoto E, Furuta T, Hioki H, Tanaka YR, Fujiyama F, Sonomura T, Uemura M, Sugiyama K, Kaneko T. A morphological analysis of thalamocortical axon fibers of rat posterior thalamic nuclei: a single neuron tracing study with viral vectors. Cerebral cortex. 2012;22(12):2840–57.

29. Manita S, Suzuki T, Homma C, Matsumoto T, Odagawa M, Yamada K, Ota K, Matsubara C, Inutsuka A, Sato M. A top-down cortical circuit for accurate sensory perception. Neuron. 2015;86(5):1304–16.

30. Guo K, Yamawaki N, Barrett JM, Tapies M, Shepherd GM. Cortico-thalamo-cortical circuits of mouse forelimb S1 are organized primarily as recurrent loops. The Journal of Neuroscience. 2020;40(14):2849–58.

31. Suzuki M, Larkum ME. General anesthesia decouples cortical pyramidal neurons. Cell. 2020;180(4):666–76.e13.

32. Larkum M. A cellular mechanism for cortical associations: an organizing principle for the cerebral cortex. Trends in neurosciences. 2013;36(3):141–51.

33. Douglas RJ, Martin KA. Recurrent neuronal circuits in the neocortex. Current biology. 2007;17(13):R496–R500.

34. Inoue M, Duysens J, Vossen JM, Coenen AM. Thalamic multiple-unit activity underlying spike-wave discharges in anesthetized rats. Brain research. 1993;612(1-2):35–40.

35. Hansel D, Mato G, Meunier C. Phase dynamics for weakly coupled Hodgkin-Huxley neurons. Europhysics Letters. 1993;23(5):367.

36. Clusella P, Politi A, Rosenblum M. A minimal model of self-consistent partial synchrony. New Journal of Physics. 2016;18(9):093037.

37. Bick C, Timme M, Paulikat D, Rathlev D, Ashwin P. Chaos in symmetric phase oscillator networks. Physical review letters. 2011;107(24):244101.

38. Matthews PC, Mirollo RE, Strogatz SH. Dynamics of a large system of coupled nonlinear oscillators. Physica D: Nonlinear Phenomena. 1991;52(2-3):293–331.

39. Hull JM, Denomme N, Yuan Y, Booth V, Isom LL. Heterogeneity of voltage gated sodium current density between neurons decorrelates spiking and suppresses network synchronization in Scn1b null mouse models. Scientific Reports. 2023;13(1):8887.

40. Rich S, Chameh HM, Lefebvre J, Valiante TA. Loss of neuronal heterogeneity in epileptogenic human tissue impairs network resilience to sudden changes in synchrony. Cell reports. 2022;39(8).

41. Hull JM, O’Malley HA, Chen C, Yuan Y, Denomme N, Bouza AA, Anumonwo C, Lopez-Santiago LF, Isom LL. Excitatory and inhibitory neuron defects in a mouse model of Scn1b-linked EIEE52. Annals of clinical and translational neurology. 2020;7(11):2137–49.

42. Sohal VS, Huguenard JR. Inhibitory interconnections control burst pattern and emergent network synchrony in reticular thalamus. The Journal of neuroscience. 2003;23(26):8978–88.

43. Coulter DA, Huguenard JR, Prince DA. Characterization of ethosuximide reduction of low-threshold calcium current in thalamic neurons. Annals of Neurology: Official Journal of the American Neurological Association and the Child Neurology Society. 1989;25(6):582–93.

44. Lüttjohann A, van Luijtelaar G. Dynamics of networks during absence seizure’s on-and offset in rodents and man. Frontiers in physiology. 2015;6:16.

45. Stuart GJ, Häusser M. Dendritic coincidence detection of EPSPs and action potentials. Nature neuroscience. 2001;4(1):63–71.

46. Larkum ME, Zhu JJ, Sakmann B. A new cellular mechanism for coupling inputs arriving at different cortical layers. Nature. 1999;398(6725):338–41.

47. Larkum ME, Zhu JJ, Sakmann B. Dendritic mechanisms underlying the coupling of the dendritic with the axonal action potential initiation zone of adult rat layer 5 pyramidal neurons. The Journal of physiology. 2001;533(2):447–66.

48. Knowles JK, Xu H, Soane C, Batra A, Saucedo T, Frost E, Tam LT, Fraga D, Ni L, Villar K. Maladaptive myelination promotes generalized epilepsy progression. Nature neuroscience. 2022;25(5):596–606.

49. Hádinger N, Bősz E, Tóth B, Vantomme G, Lüthi A, Acsády L. Region-selective control of the thalamic reticular nucleus via cortical layer 5 pyramidal cells. Nature Neuroscience. 2023;26(1):116–30.

50. Palmer L, Murayama M, Larkum M. Inhibitory regulation of dendritic activity in vivo. Frontiers in neural circuits. 2012;6:26.

51. Mashour GA. Top-down mechanisms of anesthetic-induced unconsciousness. Frontiers in systems neuroscience. 2014;8:115.

52. Boly M, Garrido MI, Gosseries O, Bruno M-A, Boveroux P, Schnakers C, Massimini M, Litvak V, Laureys S, Friston K. Preserved feedforward but impaired top-down processes in the vegetative state. Science. 2011;332(6031):858–62.

53. Alkire MT, Hudetz AG, Tononi G. Consciousness and anesthesia. Science. 2008;322(5903):876–80.

54. Llinas R, Ribary U. Consciousness and the brain: The thalamocortical dialogue in health and disease. Annals of the New York Academy of Sciences. 2001;929(1):166–75.

55. Mashour GA, Roelfsema P, Changeux J-P, Dehaene S. Conscious processing and the global neuronal workspace hypothesis. Neuron. 2020;105(5):776–98.

56. Huang Z, Zhang J, Wu J, Mashour GA, Hudetz AG. Temporal circuit of macroscale dynamic brain activity supports human consciousness. Science advances. 2020;6(11):eaaz0087.

57. Boyd RW, Gaeta AL, Giese E. Nonlinear optics. Springer Handbook of Atomic, Molecular, and Optical Physics: Springer; 2008. p. 1097–110.

58. Mitchell-Heggs R, Prado S, Gava GP, Go MA, Schultz SR. Neural manifold analysis of brain circuit dynamics in health and disease. Journal of computational neuroscience. 2023;51(1):1–21.

59. Buzsáki G, Anastassiou CA, Koch C. The origin of extracellular fields and currents—EEG, ECoG, LFP and spikes. Nature reviews neuroscience. 2012;13(6):407–20.

60. Sadleir LG, Scheffer IE, Smith S, Carstensen B, Carlin J, Connolly MB, Farrell K. Factors influencing clinical features of absence seizures. Epilepsia. 2008;49(12):2100–7.

61. Ferguson B, Glick C, Huguenard JR. Prefrontal PV interneurons facilitate attention and are linked to attentional dysfunction in a mouse model of absence epilepsy. Elife. 2023;12:e78349.

62. Smyk MK, Coenen AM, Lewandowski MH, van Luijtelaar G. Endogenous rhythm of absence epilepsy: relationship with general motor activity and sleep–wake states. Epilepsy research. 2011;93(2-3):120–7.

63. Vantomme G, Devienne G, Hull JM, Huguenard JR. The reuniens thalamus recruits recurrent excitation in the medial prefrontal cortex. Proceedings of the National Academy of Sciences. 2025;122(11):e2500321122.

64. Siegle JH, López AC, Patel YA, Abramov K, Ohayon S, Voigts J. Open Ephys: an open-source, plugin-based platform for multichannel electrophysiology. Journal of neural engineering. 2017;14(4):045003.

65. Pikovsky A, Rosenblum M, Kurths J, Synchronization A. A universal concept in nonlinear sciences. Self. 2001;2(3):10.1017.

66. Bennetin G, Galgani L, Giorgilli A, Strelcyn J-M. Lyapunov characteristic exponents for smooth dynamical systems and for Hamiltonian systems: A method for computing all of them. Meccanica. 1980;15(9):27.

67. Shimada I, Nagashima T. A numerical approach to ergodic problem of dissipative dynamical systems. Progress of theoretical physics. 1979;61(6):1605–16.

68. Zheng P, Askham T, Brunton SL, Kutz JN, Aravkin AY. A unified framework for sparse relaxed regularized regression: SR3. IEEE Access. 2018;7:1404–23.

